# Forces driving transposable element load variation during Arabidopsis range expansion

**DOI:** 10.1101/2022.12.28.522087

**Authors:** Juan Jiang, Yong-Chao Xu, Zhi-Qin Zhang, Jia-Fu Chen, Xiao-Min Niu, Xing-Hui Hou, Xin-Tong Li, Li Wang, Yong Zhang, Song Ge, Ya-Long Guo

**Affiliations:** State Key Laboratory of Systematic and Evolutionary Botany, Institute of Botany, Chinese Academy of Sciences, Beijing 100093, China; China National Botanical Garden, Beijing 100093, China; University of Chinese Academy of Sciences, Beijing 100049, China; Agricultural Synthetic Biology Center, Agricultural Genomics Institute at Shenzhen, Chinese Academy of Agricultural Sciences, Shenzhen 518000, China; State Key Laboratory of Integrated Management of Pest Insects and Rodents & Key Laboratory of the Zoological Systematics and Evolution, Institute of Zoology, Chinese Academy of Sciences, Beijing 100101, China

**Keywords:** *Arabidopsis thaliana*, genetic architecture, genetic load, natural variation, transposable elements

## Abstract

Genetic load refers to the accumulated and potentially life-threatening deleterious mutations in populations. Understanding the mechanisms underlying genetic load variation of transposable elements (TEs), one major large-effect mutations, during range expansion is an intriguing question in biology. Here, we used 1,115 globally natural accessions of *Arabidopsis thaliana*, to study the driving forces of TE load variation during its range expansion. The TE load increased with range expansion, especially in the recently established Yangtze River basin population. The effective population size explained 62.0% of the variance in TE load, and high transposition rate and positive selection or hitch-hiking effect contributed to the accumulation of TEs in the expanded populations. We genetically mapped the candidate causal genes or TEs and revealed the genetic architecture of TE load. Overall, this study reveals the variation in the genetic load of TEs during Arabidopsis expansion and highlights the causes of TE load variation.

## INTRODUCTION

Range expansion is a common process of adaptive evolution in which genetic drift usually increases the frequency of deleterious mutations on expanding wave fronts and incurs expansion load, reducing fitness and affecting the persistence of newly colonizing populations ^1^. Understanding the genetic load and its causes during range expansion is crucial for understanding species adaptation to not only the local conditions but also climate change. Expansion load has been extensively studied in diverse species ^2,3^, especially in humans ^4,5^. However, most previous studies mainly focused on nonsynonymous or loss-of-function (LoF) mutations ^6^.

The genetic load of transposable elements (TEs), one of the main contributors to large-effect mutations ^7^, remains largely unknown. TEs are repetitive DNA fragments that can transpose across the genome and constitute a large portion of the genome in many organisms ^8^. TE gain or loss is fast, thus most TEs are restricted to only a single species or a few closely related species ^9^. The transposition rate of TEs is estimated to be 4– 5 orders of magnitude higher than that of single nucleotide base mutation ^10,11^. Transposition of TEs can affect gene function and genome stability, promote phenotypic divergence, and contribute to speciation and adaptation ^12–15^.

TEs are generally regarded as deleterious and can reduce fitness via three routes: gene disruption, ectopic TE recombination, and deleterious action of TE transcript and protein products ^16^. Consistently, in natural populations, most TEs are rare insertions and are depleted in genic regions ^16–18^. In particular, a single TE or TE family can compromise host fitness ^19^. Therefore, clarifying the genetic load of TEs during range expansion is important for revealing the mechanism of invasion and environmental adaptation.

Consistent with other mutations, the evolutionary dynamics of TEs are shaped by forces acting on the processes of mutation generation (transposition and excision) and mutation maintenance (selection and genetic drift) ^20^. The transposition rate and deletion rate differ among TE families and host genotypes ^10,11^, and can affect the TE load. In particular, features of TEs and hosts can affect the transposition rate, such as the transposition capacity ^21^ and robustness of the TE-silencing system ^22^. In addition, environmental factors can also affect the TE transposition rate ^23^.

In the TE mutation maintenance process, the strength of purifying selection and genetic drift is correlated with effective population size (*N_e_*); the smaller the *N_e_* value, the more relaxed the purifying selection. Therefore, TE load is associated with *N_e_*^24^. Demographic processes (such as range expansion and founder effects) that reduce the *N_e_* could lead to the accumulation of TEs. For example, TE number in the invasive populations of *Drosophila suzukii* is considerably higher than that in its native populations ^25^. Similarly, *Capsella rubella*, a close relative of *Arabidopsis thaliana*,originated through a bottleneck, which strongly reduced its *N_e_* and exhibits a much higher TE load than its sister species ^13^.

Arabidopsis is a self-pollinating species that occurs naturally in almost all parts of the world and has over 1,000 sequenced genomes ^26–30^. Arabidopsis underwent a postglacial spread of a human commensal non-relict group, which originated near the Balkans, expanded mainly along the east-west axis, and comprised 95% of the natural populations ^31^. Therefore, Arabidopsis is a great model system for understanding the TE load variation during range expansion. Here, we explored the genomes of 1,115 globally distributed natural Arabidopsis accessions, including 128 sequenced in this study. Given that 1) the spectrum of TE load in natural populations is important for understanding the dynamics of TE load, 2) the demographic history and genetic features potentially affect the TE load, and 3) the determinant loci of TE load are crucial for understanding the TE load variation, we investigated the dynamics of TE load during range expansion, the determinant factors of TE load, and the genetic architecture of TE load variation in Arabidopsis. Overall, we elucidate the variation in TE load during Arabidopsis expansion and highlight the causes of TE load variation.

## RESULTS

### Identification and characterization of TEs in natural Arabidopsis populations

To study the genetic load of TEs during Arabidopsis range expansion, the short-read sequencing data of 1,114 globally distributed Arabidopsis accessions were utilized (Figure 1A). While the whole genomes of 986 Arabidopsis accessions were sequenced in previous studies (Table S1) ^26–28^, those of 128 accessions from northwestern China and the Yangtze River basin were sequenced in this study (Table S2). A total of 31,189 TEs belonging to 320 families were identified in the Col-0 genome (TAIR10). The TEPID software ^17^, which combines split reads and discordant reads, was used to determine the presence-and-absence variation (PAV) of TEs in Arabidopsis, with Col-0 as the reference accession. Compared with Col-0, these accessions contained, on average, 441 TE insertions and 1,257 TE deletions (Figure S1A). In the Col-0 genome, 20.9% of TEs (6,516/31,189) were polymorphic (present in at least one accession but not in all accessions), and the highest fraction (26.7%) of polymorphic TEs were accounted for by retrotransposons (Figure S1B).

**Figure 1.**
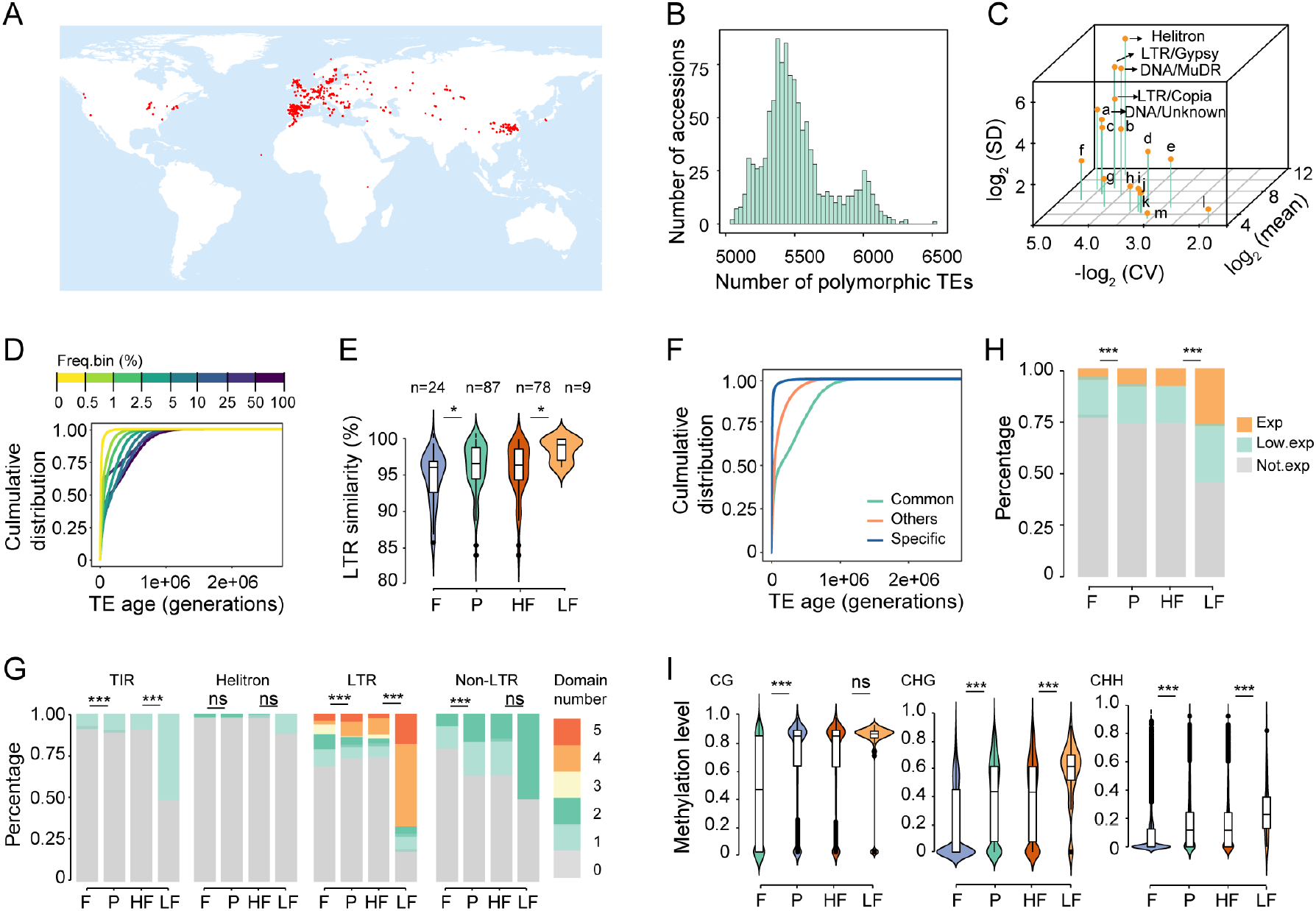
TE identification and characterization in natural Arabidopsis populations. Red dots indicate sample location. (A) Geographical distribution of 1,114 non-reference accessions. (B) Polymorphic TE numbers per accession. (C) Copy number variation of 18 TE superfamilies in 1,115 natural accessions. X-axis represents copy number variation in each TE superfamily across accessions, evaluated as coefficient of variation (CV); y-axis represents the average number of TEs in each superfamily; z-axis represents the overall contribution of each superfamily to the variation in total TE number, evaluated as standard deviation (SD). a, LINE/L1; b, DNA/En-Spm; c, DNA/HAT; d, DNA/Harbinger; e, RathE1_cons; f, DNA/Pogo; g, DNA/Mariner; h, SINE; i, Unassigned; j, DNA/Tc1; k, RathE3_cons; l, LINE?; m, RathE2_cons. (D) Age distribution of TEs in different frequency bins. (E) LTR similarity of LTR TEs in four TE categories: fixed (F; TEs with read coverage in all accessions), polymorphic (P), high frequency (HF; frequency ≥5%), and low frequency (LF; frequency <5%). Mann-Whitney U test was used for the significance test. ***, *p* < 0.001; **, *p* < 0.01; *, *p* < 0.05; ns, not significant. (F) Age distribution of TEs with different geographical distribution patterns. Common, TEs present in all populations; Others, TEs present in at least two, but not all, populations; Specific, population-specific TEs. (G) Proportion of TEs containing transposase domains in the four TE categories. Fisher’s exact test was used for the significance test. (H) Proportion of TEs in four categories, based on expression potential (the potential to express in TE-activated mutant): Exp, expressed and annotated; Low. exp, low expressed; Not. exp, not expressed. Fisher’s exact test. (I) DNA methylation levels of TEs in the four TE categories. n = 20,237 (F), 5,149 (P), 5,100 (HF), and 49 (LF). Mann-Whitney U test.

A total of 67,429 TE loci were identified among the 1,115 accessions, of which 42,756 were polymorphic. All analyses described below were conducted on polymorphic TEs unless stated otherwise. Most polymorphic TEs (80.2%) were at low frequency (frequency <5%), indicating recent mobilization or purging of TEs due to purifying selection (Figure S2). The number of polymorphic TEs per accession varied from 5,306 to 6,528 (Figure 1B), and the total number of TEs in each accession ranged from 29,979 to 31,201. Furthermore, based on the standard variation of TE numbers in diverse superfamilies across all 1,115 accessions, we estimated the contribution of each TE superfamily to the variation in TE number. The results revealed RC/Helitron, LTR/Gypsy, DNA/MuDR, LTR/Copia, and DNA/Unknown superfamilies as the top five major contributors to the variation in the total TE number (Figure 1C).

To estimate the age of polymorphic TEs, we used the Genealogical Estimation of Variant Age (GEVA), which relies on sequence divergence of regions around TEs ^32^. The results indicated that most of the polymorphic TEs transposed after the divergence of *A. thaliana* from its sister species *Arabidopsis lyrata* approximately 10 million years ago ^33^, which implies the rapid evolution and fast turnover of TEs (Figure 1D). TE frequency was commonly used to reflect TE age, of which low frequency TEs are much younger. Consistently, here the TE frequency is correlated with TE age (Figure 1D). The age estimation of long-terminal repeat (LTR) TEs, based on the diversification of two LTR sequences on either end of each intact LTR TE, also indicated that polymorphic TEs and low-frequency TEs were younger than fixed TEs and high-frequency TEs, respectively (Figure 1E). Additionally, in terms of the geographical distribution data, region-specific TEs were much younger than the globally distributed TEs (Figure 1F). Taken together, TE gain and loss is fast and the polymorphic TEs were transposed recently after speciation.

To characterize the transposition activity of polymorphic TEs, which are most probably active, we analyzed their structural integrity, transcription potential, and DNA methylation level. Given that data on these genetic features are mostly abundant in Col-0, all comparative analyses were based on Col-0. Firstly, to characterize the transposition potential of TEs, we searched the total number of transposition-related domains for each TE. Higher fraction of polymorphic TEs had transposition-related domains than that of fixed TEs, except Helitrons (whose transposition-related domains were similar between polymorphic and fixed TEs) and LTR TEs (Figure 1G). However, although a lower proportion of polymorphic LTR TEs contained transposition-related domains, the completeness of polymorphic LTR TEs was higher than that of fixed LTR TEs (Figure 1G). Secondly, given that most TEs are repressed by the host genome in Col-0, to evaluate the transcription potential of TEs, we utilized the published long-read TE transcriptome data of Col-0 triple mutants *(ddm1, rdr6*, and *pol V*), which lack multiple layers of TE repression and potentially reflect the transcription potential of TEs ^34^. Compared with fixed TEs, a much higher proportion of polymorphic TEs were expressed in the TE-activated mutants (Figure 1H). Thirdly, to clarify the extent of DNA methylation-induced TE repression, we calculated the DNA methylation levels of polymorphic and fixed TEs. The results showed that cytosines in all contexts were methylated to a higher degree in polymorphic TEs than in fixed TEs (Figure 1I).

Given that low-frequency TEs are a more accurate indicator of recent mobilization than high-frequency TEs, we compared the above three features between low-frequency (frequency <5%) and high-frequency (frequency ≥5%) TEs. Low-frequency TEs possessed more transposition-related domains, especially for terminal inverted repeat (TIR) and LTR TEs (Figure 1G), and showed higher transcription potential (Figure 1H) and higher CHG and CHH methylation levels than high-frequency TEs (Figure 1I). Taken together, these results suggest that polymorphic TEs, particularly low-frequency TEs, contain more transposition-related domains and are more likely to be transcribed, which help them or other nonautonomous members to transpose. However, at the same time, these recently active TEs tend to be silenced by DNA methylation, suggesting that hosts could identify and silence potentially active TEs.

### Deleterious effects of TEs and synergistic epistasis among TEs with large fitness effects

TEs have been demonstrated to be deleterious ^16,19,35^. However, genome-wide exploration of the deleterious effects of TEs is lacking. To detect the deleterious effects of TEs, we analyzed the site frequency spectrum (SFS) of TEs and other genetic variations. The more deleterious mutations are more skewed toward low frequency because of purifying selection ^36^. Four-fold degenerate sites (which are putatively neutral), nonsynonymous single nucleotide polymorphisms (nSNPs) that can be tolerated (tnSNPs) or have deleterious effects (dnSNPs) as predicted by Provean ^37^, and loss-of-function (LoF) mutations were used as comparisons to infer the potential fitness effect of TEs. The SFS of total TEs was more skewed toward the left than that of the less deleterious tnSNPs but less skewed than that of the more deleterious dnSNPs and LoF mutations (Figure 2A), indicating that the deleterious effect of TEs varies between that of tnSNPs and dnSNPs.

**Figure 2.**
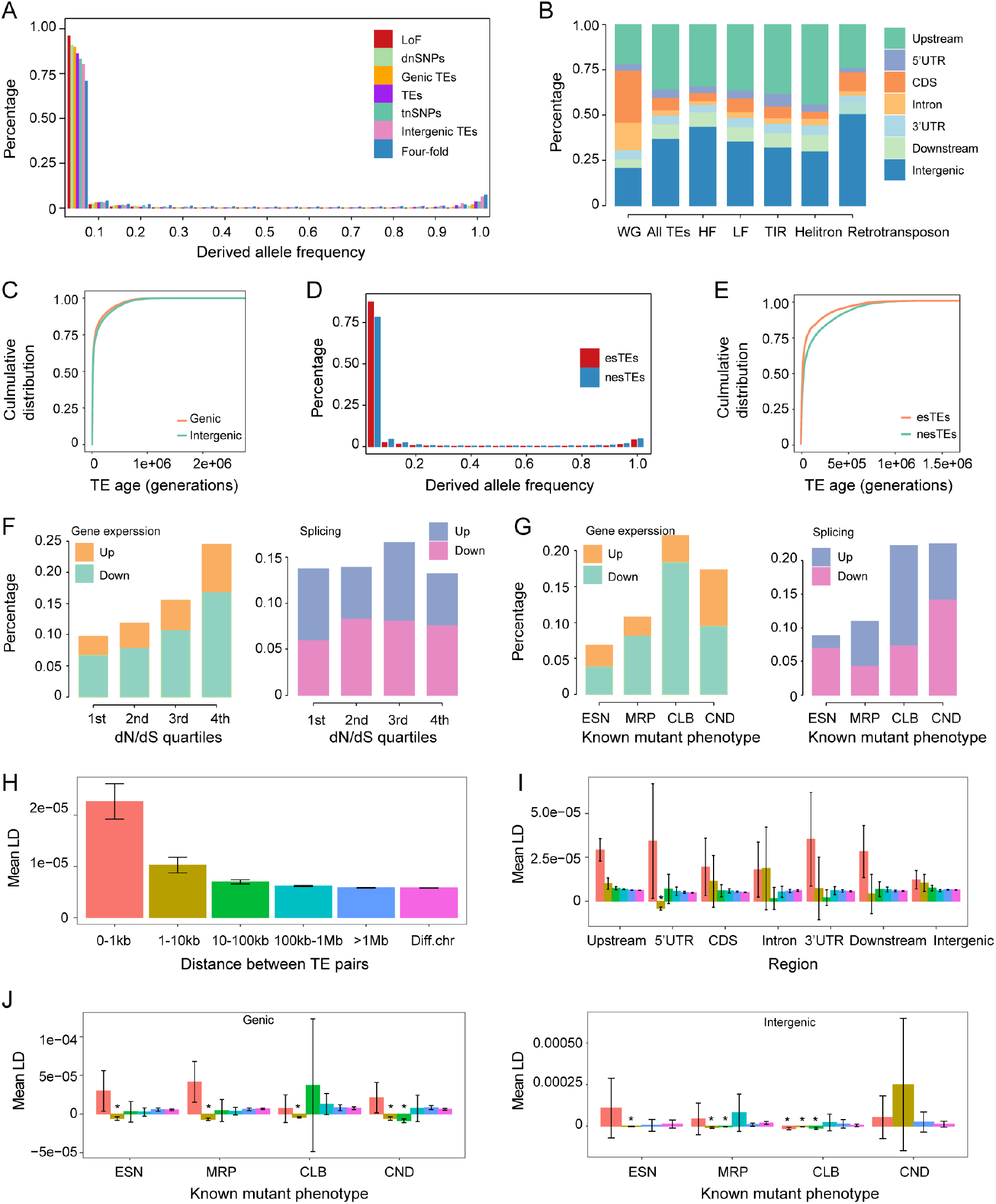
Evaluation of the deleterious effects of TEs. (A) Derived allele frequency of TEs and other variants. Four-fold, four-fold degenerate sites; dnSNPs, deleterious nonsynonymous SNPs; tnSNPs, tolerated nonsynonymous SNPs; LoF, loss-of-function variants; Genic TEs, TEs in the gene body and flanking region (2 kb upstream and 1 kb downstream); Intergenic TEs, TEs outside genic regions. (B) Distribution of TEs in different genomic regions. WG, whole genome; HF, high frequency; LF, low frequency; Upstream, 2 kb upstream of the start of the gene body; Downstream, 1 kb downstream of the end of the gene body. (C) Age distribution of genic and intergenic TEs. (D) Derived allele frequency of TEs capable of affecting gene expression and splicing (esTEs) and not capable of affecting gene expression and splicing (nesTEs). (E) Age distribution of esTEs or nesTEs. (F and G) Percentage of TEs associated with at least two-fold change in expression level or PSI value compared with the nearest genes grouped into four conservation categories based on their dN/dS quartiles (F) or mutant phenotypes (G). In (F), the first quartile (1st) was the most constrained, while the last quartile (4th) was the least constrained. In (G), the categories defined according to known mutant phenotypes were as follows: ESN, essential; MRP, morphological; CLB, cellular-biochemical; CND, conditional. (H) Mean LD of all TE pairs with different physical distances. Diff. chr, TE pairs located on different chromosomes. Error bar indicates 95% confidence interval. (I) Mean LD of TE pairs located in different genomic regions. Asterisk (*) denotes significant negative LD with a completely negative 95% confidence interval. Bar color denotes the physical distance between TEs of a pair. (J) Mean LD of TE pairs located within genic or intergenic regions of genes in different functional categories. For distance bins, whose number of TE pairs was not sufficient (see MATERIALS AND METHODS), are not shown.

To further characterize the deleterious effect of different types of TEs, we adopted three approaches that rely on the relationship of TEs with their inserted or adjacent genes.

The first approach is based on the location of TE insertion. TEs are enriched in pericentromeric regions, where genes are strongly depleted (Figure S3A). Similarly, TEs are enriched in intergenic regions as well (Figure 2B). In contrast, in genic regions (i.e., gene body, 2 kb upstream region, and 1 kb downstream region), TEs are largely depleted, especially in the coding sequence (CDS) and intronic regions (Figure 2B), which implies that TEs in genic regions are more deleterious than intergenic regions. Consistently, the SFS of genic TEs is more skewed than that of total TEs and tnSNPs but less skewed than that of dnSNPs and LoF mutations (Figure 2A), which implies that the deleterious effect of genic TEs varies between that of total TEs and dnSNPs. Similarly, based on the SFS, the potential fitness effect of intergenic TEs varies between that of four-fold degenerate sites and tnSNPs (Figure 2A), and intergenic TEs are less deleterious than genic TEs. Consistent with previous studies about SNPs, which showed that deleterious mutations are younger than neutral or benign mutations ^32,38^, we found that the more deleterious genic TEs were younger than the less deleterious intergenic TEs (Figure 2C, *p* = 2.4e-06, Mann-Whitney U test).

The second approach is based on the ability of TEs to affect gene expression and splicing. We used the RNA-seq data of 413 accessions ^39^ to evaluate the effects of TEs on gene transcription and splicing. The results showed that 23.1% and 14.2% of the TEs were associated with at least two-fold change in the expression level and percent spliced in (PSI) value of the inserted or adjacent genes, respectively. In general, TEs in the CDS or intronic regions were more likely to regulate gene expression and splicing than other regions, and mainly downregulated gene expression and inhibited splicing (Figure S3B and S3C). Given that genic TEs were generally deleterious (Figure 2A and 2B), and that stabilizing selection constrains the variation in gene expression ^40^, the TEs affecting gene expression or splicing were most probably deleterious. To validate this assumption, we compared the SFS of TEs altering gene expression or splicing (esTEs) with that of TEs incapable of regulating gene expression or splicing (nesTEs). As expected, the SFS of esTEs was more skewed toward rare variations than that of nesTEs, which implies that esTEs are more deleterious than nesTEs (Figure 2D). Consistent with our result that the more deleterious genic TEs were younger than the less deleterious intergenic TEs (Figure 2C), the more deleterious esTEs were much younger than the less deleterious nesTEs (Figure 2E, *p* < 2.2e-16, Kolmogorov-Smirnov test and Mann-Whitney U test). Thus, TEs altering gene expression or splicing are more likely to be deleterious.

The third approach relies on the functional importance of genes adjacent to or contained TEs. TEs located within or near functionally important genes and those affecting gene function would potentially be more deleterious and thus would be selected against. To verify this assumption, we categorized TEs based on the conservation of the nearest genes and tested if genes with more essential functions were depleted of TEs that affect gene function. The conservation categories were based on the dN/dS ratio; the smaller the dN/dS ratio, the more conserved and functionally essential the gene ^41,42^. Consistently, more constrained genes were depleted of TEs that could affect the gene expression level (Figure 2F). We also categorized the functional essentiality of genes based on their known mutant phenotypes ^43^. The functionally important genes were depleted of TEs that could affect the gene expression level and splicing (Figure 2G). These results suggest that TEs located close to functionally important genes are more deleterious than TEs near less important genes.

The classic theoretical framework predicts that purifying selection acting on the synergistic epistasis of deleterious TEs (whereby an additional TE copy exacerbates the reduction of fitness), rather than selection against insertional mutations, is predominantly responsible for TE abundance maintenance ^20,44^. To explore the possible effect of the synergistic epistasis of TEs, we utilized the repulsion linkage disequilibrium (LD) of TE pairs generated by purifying selection and computed the mean LD of TE pairs with different physical distances. Here, we used TEs with frequency < 1% to maximize the deleterious effect of TEs and thus improve the likelihood of the identification of the synergistic epistasis of TEs.

At the genome level, the mean LD and 95% confidence interval of all TE pairs with different physical distances were positive (Figure 2H). Because purifying selection is expected to generate more negative LD among the highly deleterious TE pairs, we focused on the mean LD of TE pairs with more deleterious effects. Among the potentially more deleterious genic TEs, the LD of TE pairs located 1–10 kb apart and in 5’ untranslated regions (5’UTRs) was significantly negative (Figure 2I). We also analyzed the LD of TE pairs inserted or adjacent to functionally essential genes. When genes were categorized according to their conservation score, we observed significantly negative LD values of TE pairs located in the intergenic regions (1–10 kb bin) of the most constrained genes (Figure S3D). When genes were categorized according to their mutant phenotype, the LD values of TE pairs located in genic regions were significantly negative for all phenotypic categories (1–10 kb bin) and conditional categories (10–100 kb bin) (Figure 2J). Interestingly, we also observed significantly negative LD for TE pairs located in intergenic regions when the mutant phenotype of the closest gene was categorized as essential (1–10 kb bin), morphological (1–10 and 10–100 kb bins), and cellular-biochemical (0–1, 1–10, and 10–100 kb bins) (Figure 2J). However, unlike in *Drosophila melanogaster* ^45^, we found significantly negative LD only between the more deleterious TE pairs but not at genome level, which might be ascribed to the weakly deleterious effects of TEs in *A. thaliana* (the SFS of dnSNPs and LoF mutations was more skewed to the left than that of TEs) compared with the strongly deleterious effects of TEs in *D. melanogaster* (the SFS of TEs is more skewed to the left than that of LoF mutations). Overall, these results suggest that the synergistic epistasis of TEs is present in *A. thaliana* and tends to be prevalent among TE pairs with more deleterious effects.

### TE load increased with the distance from the origin and was negatively correlated with *N_e_*

To study the TE load variation during the range expansion of *A. thaliana*, we focused on ten non-relict populations. The Balkans population was regarded as the origin of non-relicts as it is located near the predicted origin of non-relicts (Figure 3A) ^26,31^. Consistent with the expansion from Balkans, the genetic diversity of natural populations decreased with expansion from the origin (Figure 3B). All expanded populations, except the Spain population, showed significantly lower genetic diversity than the Balkans population (Figure 3B, *p* < 0.01, Mann-Whitney U test); the higher genetic diversity of the Spain population could have resulted from the introgression of Iberian relicts ^31^. Populations located on the margin of expansion, especially the Yangtze River basin population, had much lower genetic diversity (reduced by 60.7% relative to the Balkans) (Figure 3B).

**Figure 3.**
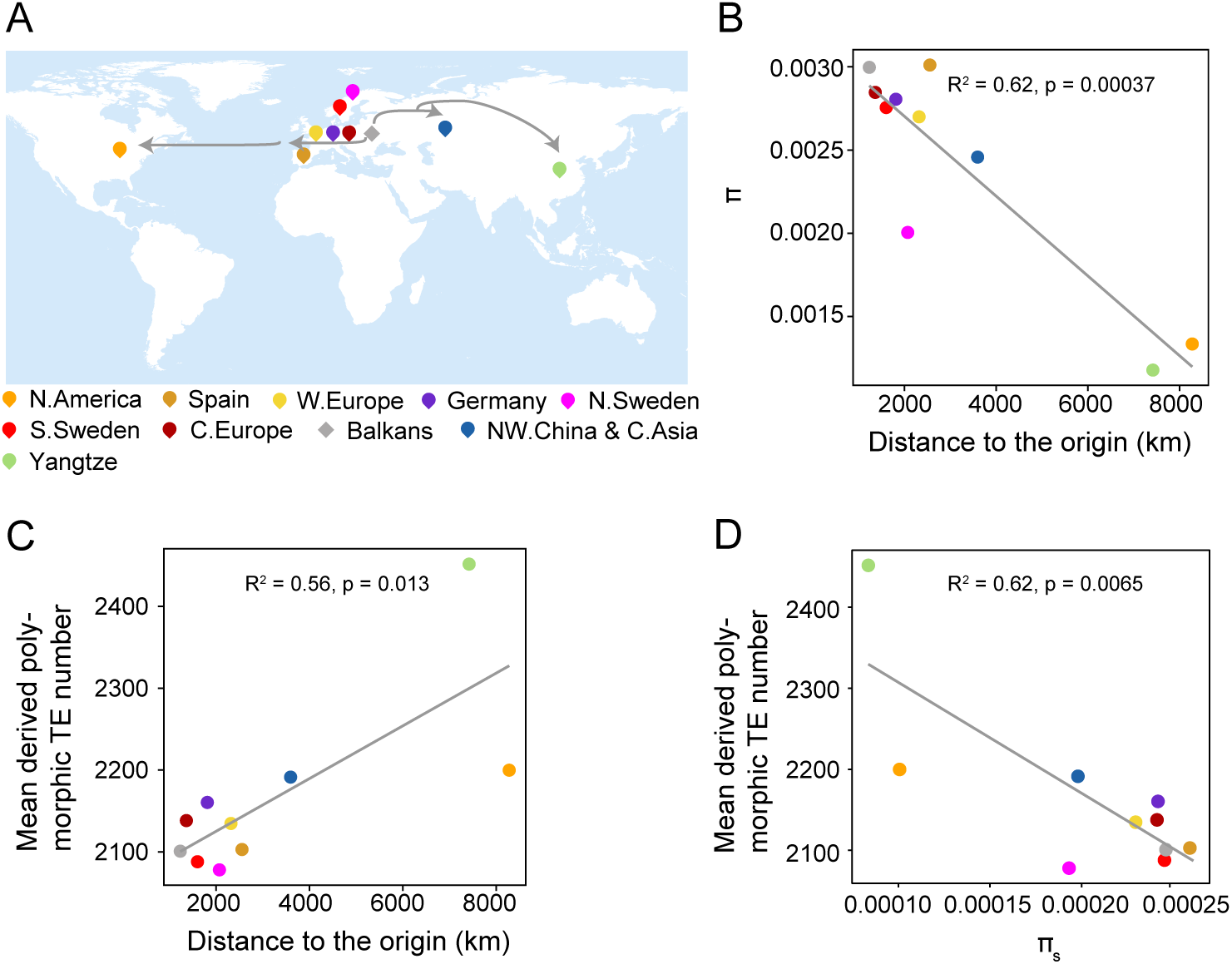
TE load variation during Arabidopsis range expansion. (A) Geographical representation of the range expansion of non-relicts. Locations of ten non-relict populations are indicated on the map. The putative expansion trace is indicated with arrows. (B) Correlation between genetic diversity (π; calculated in non-overlapping 10 kb windows) and the distance to the putative origin. (C) Correlation between TE load and the distance to the putative origin. (D) Correlation between the mean derived TE number and effective population size, used four-fold degenerate sites diversity (⊓_s_) as proxy. Genetic diversity was calculated in non-overlapping 10 kb windows.

To estimate TE load variation during range expansion, the derived polymorphic TE number per individual (TEs present in *A. thaliana* but absent in *A. lyrata* and *C. rubella)* was used as a load proxy. Consistent with the theoretical prediction of expansion load ^1^, TE load increased with the distance from the origin. Most expanded populations showed higher TE load than the origin Balkans population (*p* < 0.01, Mann-Whitney U test), and populations on the expansion wave front, especially the recently established Yangtze River basin population, exhibited the largest TE load (increased by 16.7% relative to the Balkans) (Figure 3C).

*N_e_*, reflected by nucleotide diversity at four-fold degenerate sites ^24^, was directly correlated with the selection coefficient. The linear regression between TE load and nucleotide diversity at four-fold degenerate sites in non-relict populations suggested that *N_e_* alone explained 62.0% of the TE load variation among the natural populations (Figure 3D). Consistently, populations on the expansion front had much lower *N_e_* and accumulated much higher TE load than the origin Balkans population (Figure 3A). The exception of the Spain population might be explained by the introgression of Iberian relicts, which increased the genetic diversity of this population ^31^. However, the reason for the exception of the two Swedish populations remains unclear and awaits further study.

Non-relicts were demonstrated to spread mainly along the east-west axis ^31^. Intriguingly, TE load differed between the western and eastern expansion fronts (Figure 3A). On the western expansion fronts, the reduction in *N_e_* was mild, and the TE load was only slightly higher in the expanded populations. In contrast to the western expansion fronts, the eastern expansion fronts showed a clear expansion trace ^28,46^. *N_e_* decreased with the distance to Balkans, and TE load increased along the expansion axis, reaching the highest at the most eastern expansion front. In subsequent analyses, we focused on gaining in-depth insights into the dynamics, causes, and consequences of the TE load on the eastern expansion fronts, especially the Yangtze River basin population.

### High transposition rate and positive selection or hitch-hiking effect contribute to high TE load in the Yangtze River basin population

To gain further insight into the TE load variation during range expansion, we focused on the Yangtze River basin population. The load of TEs with different deleterious effects, as predicted in Figure 2, was compared between the closely related northwestern China and Central Asia populations (NW. China & C. Asia population). Between the two populations, the load of TEs with different deleterious effects was much higher in the Yangtze River basin population than in the NW. China & C. Asia population (Figure 4A and Figure S4A, *p* < 0.01, Mann-Whitney U test).

**Figure 4.**
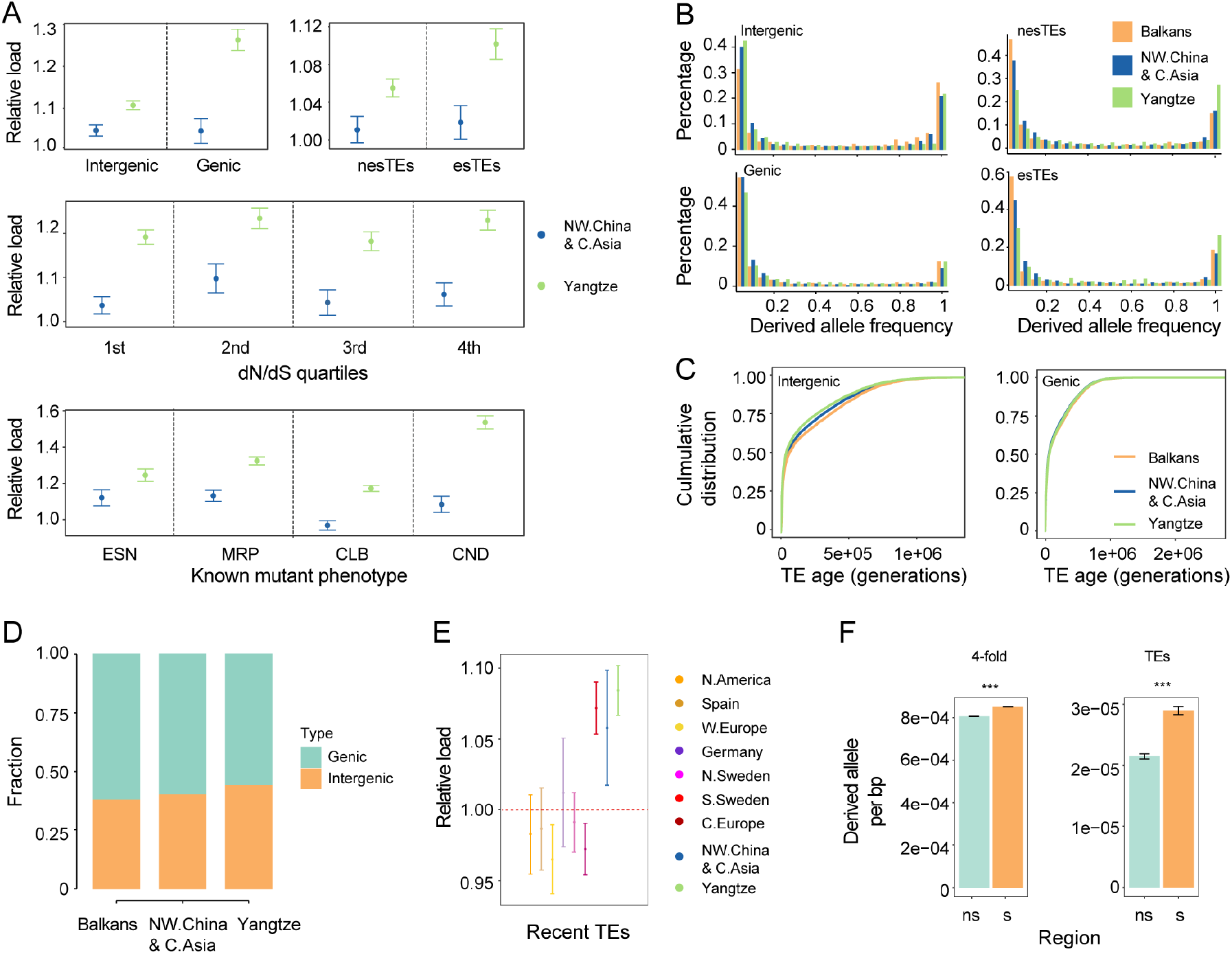
Causes of high TE load in the Yangtze River basin population. (A) Load of TEs with different deleterious effects as classified in Figure 2. The mean number of derived alleles in each variant category in the Balkans population was used as the standard for calculating the relative load. Error bars represent 95% confidence intervals. (B) Comparison of the SFS of different TEs among three populations. (C) Age distribution of TEs with different deleterious effects in the three populations. (D) Mutation type of genic and intergenic TEs in different populations. (E) Comparison of recent TE load (top 5% youngest TEs) among non-relict populations. The mean number of recent TEs in the Balkans population was used as a standard for calculating the relative load. Error bars represent 95% confidence intervals. (F) Derived allele counts per bp in selective sweep regions (s) and non-selective sweep regions (ns) of the Yangtze River basin population. Mann-Whitney U test was used for significance test. ***, *p* < 0.001.

Range expansion is assumed to cause genetic surfing, a phenomenon where allele frequency increases because of strong genetic drift ^47^. To test the effect of genetic surfing on the accumulation of TEs in the Yangtze River basin population, we compared the SFS of TEs in the Yangtze River basin population with that in the NW. China & C. Asia and Balkans populations. The Yangtze River basin population showed fewer low-frequency TEs and an excess of high-frequency and fixed TEs in all TE classes, except intergenic TEs, which supported the effect of genetic surfing on the accumulation of TEs in the Yangtze River basin population (Figure 4B and Figure S4B). As theoretically expected ^48^, genetic surfing was most likely caused by the lowest effective population size of the Yangtze River basin population (Figure 3D).

The exception that the SFS of intergenic TEs did not show a rightward shift compared with the Balkans population and its sister populations implied that other factors might contribute to the higher intergenic TE load in the Yangtze River basin population. Given the less efficient purifying selection in the Yangtze River basin population, the excess of rare intergenic TEs in this population suggested that these rare TEs were recent transpositions, indicating increased transposition rate. Intergenic TEs in the Yangtze River basin population were much younger than those in the NW. China & C. Asia and Balkans populations (Figure 4C, *p* < 0.01, Kolmogorov-Smirnov test and Mann-Whitney U test), while genic TEs in the Yangtze River basin population showed a similar age distribution as the NW. China & C. Asia and Balkans populations (Figure 4C, *p* > 0.05, Kolmogorov-Smirnov test). By contrast, both nesTEs and esTEs were older in the Yangtze River basin population than in the NW. China & C. Asia and Balkans populations (Figure S4C, *p* < 0.01, Kolmogorov-Smirnov test and Mann-Whitney U test). The mutation spectrum also suggested that the Yangtze River basin population had more intergenic TEs than each of the other two populations (Figure 4D, *p* < 0.01, Chi-square test). To further support this result, we compared the load of the most recent TEs (top 5%of the age distribution, less than 700 generations), which likely reflect recent mobilization and are less likely under selection, among non-relict populations. The most recent TE load in the Yangtze River basin population was also the highest (Figure 4E and Figure S4D, *p* < 0. 01, Mann-Whitney U test), indicating that high transposition rate is another contributor to the high TE load of Yangtze River basin population.

Hitch-hiking through positive selection has been demonstrated to contribute to the accumulation of deleterious mutations ^49^. To test if positive selection contributes to TE accumulation in the Yangtze River basin population, we compared the derived allele count per base-pair in selective sweep regions with that in regions not under selective sweep. Both the four-fold degenerate sites and TEs were significantly enriched in selective sweep regions (Figure 4F). The enrichment was stronger for TEs (1.35-fold enrichment) than for four-fold degenerate sites (1.06-fold enrichment) (Figure 4F). By contrast, in the NW. China & C. Asia population, at the syntenic region of the selective sweep in the Yangtze River basin population, both the four-fold degenerate sites and TEs were not enriched (*p* > 0.05, Mann-Whitney U test). Because TEs could be highly adaptive to new environments ^13^, it is likely that the enrichment of TEs in selective sweep regions was caused not only by the hitch-hiking effect but also by the TEs themselves as targets of positive selection. Thus, positive selection or hitch-hiking effect contributed to TE accumulation in the Yangtze River basin population.

### Genetic architecture of TE expression variation

To dissect the genetic architecture of natural variation in TE load, we focused on two stages of the TE transposition process (Figure 5A): the initiation stage of transcription and the final stage of copy number variation. We first aimed to disentangle the genetic factors responsible for TE expression.

**Figure 5.**
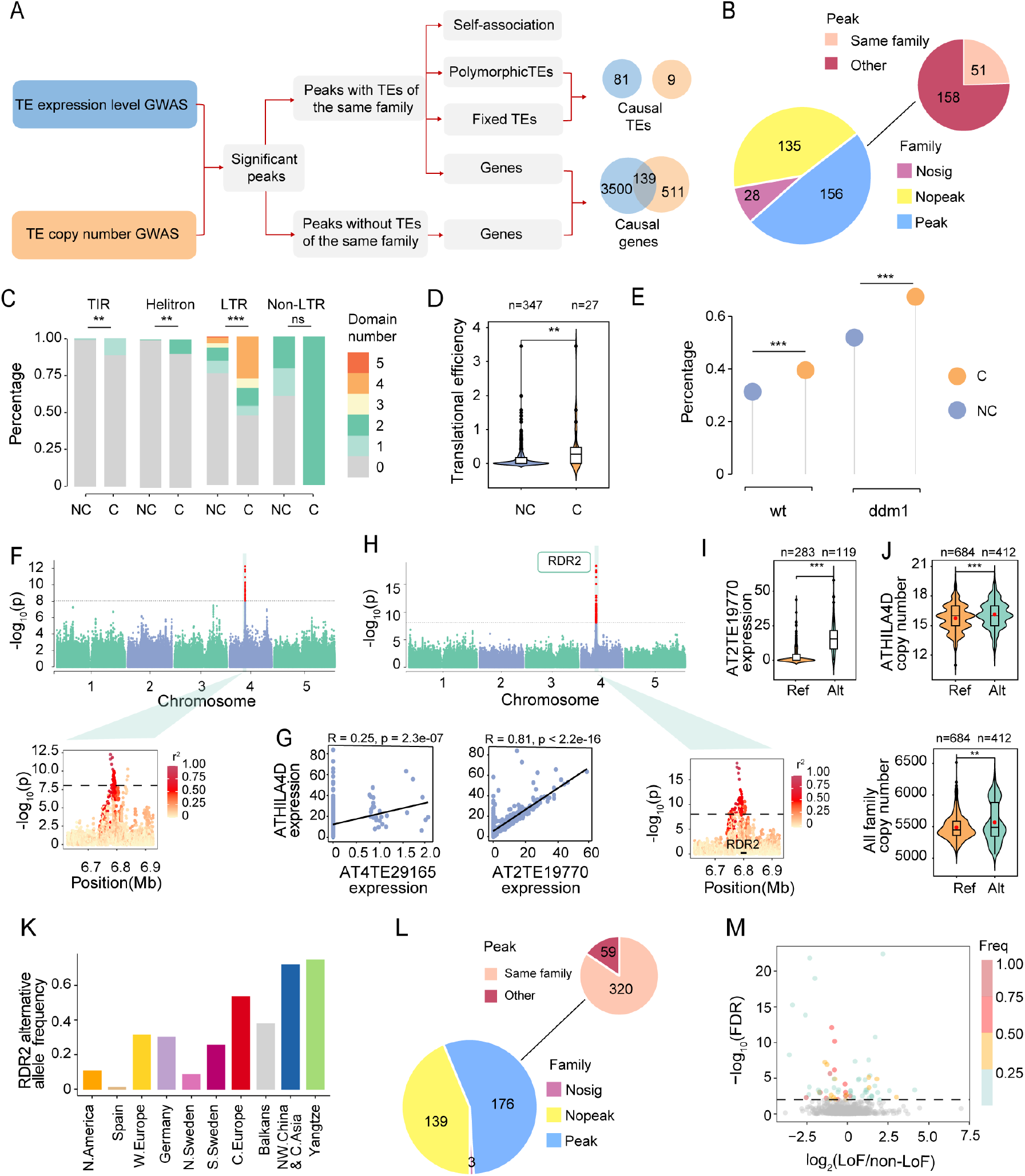
Genetic factors associated with TE family expression and copy number variation. (A) Pipeline and strategy used to identify causal loci associated with TE load variation. (B) Summary of the results of TE family expression level GWAS. Nosig, TE families without significant SNPs; Nopeak, TE families without significant peaks; Peak, TE families with significant peaks; Same family, peaks with TEs of the same family; Other, peaks without TEs of the same family. (C) Proportion of TEs containing the transposase domain. TEs were divided into two categories: non-causal (NC; TEs of the same family but not in the candidate interval) and causal (C). Fisher’s exact test was used for significance test. ***, *p* < 0.001; **, *p* < 0.01. (D) Translational efficiency of causal TEs. Mann-Whitney U test was used for significance test. (E) Proportion of LTR TEs capable of producing transposition intermediates in two genotypes: wild type (wt) and *ddm1* mutant (*ddm1*). Fisher’s exact test. (F) Manhattan plot and local Manhattan plot of GWAS of the ATHILA4D family expression level. (G) Correlation between the ATHILA4D family expression level and the expression level of two members of this family. (H) Manhattan plot and local Manhattan plot of the GWAS of AT2TE19770 expression level. (I) Expression levels of the two alleles of AT2TE19770. Alleles were grouped based on the lead SNP since the deleterious missense SNPs were in strong LD with the lead SNP (r^2^ = 0.88). Ref, reference allele; Alt, alternative allele. Mann-Whitney U test. (J) Copy number variation of the ATHILA4D family and polymorphic TEs between accessions with reference and non-reference *RDR2* alleles. Red dots indicate mean copy number. Mann-Whitney U test. (K) Frequency of the *RDR2* alternative allele in ten non-relict populations. (L) Summary of TE family copy number GWAS results. (M) Differences in TE abundance related phenotypes (TE family expression level and copy number) between accessions with LoF mutations and those without LoF mutations in candidate genes identified by GWAS. Horizontal line indicates 1% FDR. LoF/non-LoF indicates the ratio of TE abundance related phenotypes in accessions with LoF mutations to those in accessions without LoF mutations in TE regulatory genes. Freq indicates the frequency of LoF alleles in the Yangtze River basin population.

Based on the published high-coverage RNA-seq data of 414 Arabidopsis accessions (including Col-0) ^39^, we measured TE expression at both the family and locus levels. At the family level, by taking the TE superfamily size into account, the unassigned superfamilies showed the highest expression level, followed by Copia, LINE, SINE, and Harbinger superfamilies (Figure S5A). At the locus level, among the 31,189 TEs in the Col-0 (reference) genome, 16,974 (53.8%) were expressed in at least one accession (Figure S5B). A total of 13 TEs were expressed in all 414 accessions. Most TEs (81.1%) were expressed in only a few accessions (frequency < 0.05) (Figure S5C). Analysis of the percentage of expressed TEs of each type revealed that more retrotransposons were transcribed than DNA transposons (including TIR and Helitron) (Figure S5D).

The quantification of TE expression is much more accurate at the family level than at the single locus level because of sequence similarity among TEs within the same family. Therefore, expression level variation among TE families, instead of among the different TE loci, was utilized to identify the causal loci in the subsequent genome-wide association study (GWAS). Among the 319 TE families expressed in at least one accession, 291 families showed significant association signals (22,167 significant SNPs) (Figure 5B). Here, we focused on 209 significant peaks from the GWAS of 156 TE families since peaks most likely have large effects on phenotypes (Table S3). Among the 209 peaks, 51 peaks encompassed at least one TE belonging to the family under investigation (Figure 5A, Table S3). In total, we identified 40 fixed and 41 polymorphic candidate causal TEs in these 51 peaks (Table S4).

Among the 81 candidate causal TEs, the expression levels of 23 causal TEs were strongly correlated with those of their corresponding families (r > 0.75), indicating that these causal TEs were high-confidence causal candidates for the expression variation of their families (Table S5). The strong expression-level correlation between candidate TEs and their families suggested that most members of these families, except candidate TEs, were nearly not expressed. In addition, the association signal might result from TE sequence variation or TE PAV. Here, among the 40 fixed candidate causal TEs, we selected AT4TE52315, the causal locus of ATCOPIA10 family expression variation, for further analysis (Figure S6A). The expression level of ATCOPIA10 family (or AT4TE52315) was significantly lower in accessions with the reference allele (note: alleles were grouped based on the lead SNP) (Figure S6B and S6C). By contrast, in cases where polymorphic TEs acted as the causal loci, for example, the PAV of AT3TE63765 was responsible for the expression variation of the ATCOPIA62 family (Figure S7A and S7B). However, for some TE families, such as the ATCOPIA70 family, both the TE PAV and sequence variation contributed to its expression variation. We identified AT2TE29450 as the causal locus in the association analysis of ATCOPIA70 family expression level (Figure S8A); accessions with AT2TE29450 showed much higher ATCOPIA70 expression levels than accession without AT2TE29450 (Figure S8B). Additionally, AT2TE29450 was identified again in the association analysis of the ATCOPIA70 family expression level, which was performed using only 348 accessions with AT2TE29450 (Figure S8C). Accessions with the nonreference allele showed higher ATCOPIA70 family and AT2TE29450 expression levels than those with the reference allele (Figure S8D).

To characterize the transposition potential of candidate causal TEs, we compared the transposition potential of causal TEs with that of other TEs belonging to the same family. We divided the TEs into two categories: non-causal (TEs belonging to the same family but not in the peak) and causal. Compared with non-causal TEs, a higher proportion of causal TEs, especially TIR TEs, LTR TEs, and Helitrons, contained transposition related domains (Figure 5C). Furthermore, the translational efficiency of causal TE genes was significantly higher than that of non-causal TE genes (Figure 5D). In addition, we measured the transposition potential of causal TEs using the DNA-seq data of virus-like particles (VLPs), which reflect the capacity of LTR TEs to produce transposition intermediates ^50^. The results showed that a much higher proportion of causal LTR TEs was able to produce transposition intermediates (Figure 5E). Together, these results suggest that causal TEs have higher transposition potential than other TEs of the same family.

Despite 51 peaks containing TEs of the same family, it is possible that causal loci within some peaks are genes rather than TEs. In these 51 peaks, we identified a total of 1,373 genes (Table S6). For example, in the association analysis of the ATHILA4D family, *RNA-dependent RNA Polymerase 2 (RDR2)*, which functions in RNA-directed DNA methylation (RdDM) pathway by converting single-stranded RNA into double-stranded RNA ^51^, was identified at a peak on chromosome 4 (chr4: 6,777,205) (Figure 5F). This peak also contained a member of the ATHIL4D family, AT4TE29165; however, AT4TE29165 was silenced in most accessions, and its expression level was weakly correlated with that of the ATHIL4D family (Figure 5G). Instead, another member of the ATHIL4D family on chromosome 2, AT2TE19770, was highly expressed, and its expression level was strongly correlated with that of the ATHIL4D family (Figure 5G). We further conducted association analysis of the expression level of AT2TE19770 (unique mapping rate = 0.97) and identified the same peak (chr4: 6,777,205) in the GWAS of the family expression level (Figure 5H). These results indicate that *RDR2*, rather than AT4TE29165, is the candidate causal locus of this peak. However, the expression level of *RDR2* was similar between the two alleles. Additionally, five significant SNPs were identified in the *RDR2* gene, two of which were missense mutations, and the missense mutation (chr4: 6,784,172) that occurred in the functionally important C-terminal head domain ^52^ was deleterious as predicted by Provean. Accessions with the non-reference allele exhibited a higher expression level of AT2TE19770 (Figure 5I). In addition, the copy number of the ATHILA4D family and all TEs families at the genome level was significantly higher in accessions with the non-reference allele (Figure 5J). Interestingly, the non-reference allele of *RDR2* occurred at high frequency in the Yangtze River basin population (Figure 5K) and potentially contributed to TE expansion.

A total of 158 peaks did not contain TEs belonging to the same family (Table S3), and most of the casual loci in these peaks were probably genes. These 158 peaks contained 2,518 genes (Table S6), including genes involved in epigenetic regulation, such as the DNA demethylation gene *DML2* (Figure S9). In summary, we identified 81 candidate TEs and 3,640 candidate genes, including 22 well-known TE regulatory genes (Table S6), in the 209 peaks of GWAS. Gene Ontology enrichment analysis revealed no specific GO terms for these 3,640 candidate genes. Overall, these TEs and genes represent the candidate causal loci responsible for natural variation in the TE family expression level.

### Genetic architecture of TE copy number variation

To further investigate the mechanism of natural variation in TE load, we performed association analysis and determined the genetic architecture of TE family copy number variation among the 1,115 natural accessions. Among the 318 TE families with copy number variation, 315 showed significant association signals (156,995 significant SNPs) (Figure 5L). To identify the causal loci, we focused on 176 TE families with a total of 379 significant peaks (Table S7).

Out of 379 significant peaks, 320 peaks encompassed at least one TE belonging to the same TE family as that being studied (Figure 5L, Table S7). This association signal might have resulted from differences in the PAV or sequence variation of TEs or from the linkage between TE and the adjacent causal SNP. To identify candidate causal TEs (Figure 5A), we first excluded 315 peaks with polymorphic TEs belonging to the same family; this strategy was different from that adopted in the association study of TE family expression level. In the TE expression level GWAS, the PAV of actively transcribed TEs could affect the fate of the TE family through transcription and further transposition. However, the association in TE copy number GWAS probably originated from the PAV of TEs (self-associated, the presence of a TE is associated with a single copy number gain) rather than from the real contribution of active TEs (the presence of a TE is associated with multiple copy number gain). The remaining 5 out of 320 peaks contained fixed TEs, which were likely causal TEs (Table S8). For example, in the GWAS of the ATCOPIA68 family copy number, significant SNPs were detected in AT1TE62960, a member of this family (Figure S10A). Sequence variation of AT1TE62960 potentially contributed to its family copy number variation (Figure S10B). However, these causal TEs did not overlap with causal TEs identified in the family expression level GWAS, mostly probably because we ruled out the self-associated TEs in the copy number analyses, leaving behind only a few causal TEs (Figure 5A).

Although the 320 peaks contained TEs belonging to the same family, it is possible that the causal elements in some peaks were genes rather than TEs, especially in peaks without candidate causal TEs. We identified 88 candidate genes in the five peaks that contained fixed TEs of the same family (Table S9). In the 59 out of 379 peaks that did not contain any TEs belonging to the same family, genes were most likely the causal loci. These 59 peaks harbored a total of 590 genes (Table S9), two of which *(DRD1* and *NERD)* were previously reported to repress TEs ^53,54^.

In summary, among the 379 peaks of GWAS, 315 peaks were potentially self-associated, and 651 genes (Table S9) and 9 TEs (Table S8) were identified in the remaining 64 peaks as candidate causal loci of TE family copy number variation. However, similar to the TE expression association, no specific GO term enrichment was detected for these 651 genes. Intriguingly, 139 candidate genes identified at the copy number variation stage overlapped with those identified at the TE expression stage and were enriched in the salicylic acid signaling pathway (Figure 5A, Table S10). The observed number of overlapping genes was significantly higher than the expected number (*p* < 0.01), implying that some genes or pathways were involved in both TE expression and copy number variation.

### LoF mutations of candidate genes were associated with TE family expression and copy number variation

To verify the causality of candidate genes identified by the GWAS of TE expression and copy number, we utilized the natural LoF mutations and determined if the candidate genes contributed to TE load variation. Furthermore, we tested if accessions with LoF mutations in candidate genes differed from those with non-LoF mutations in TE family expression level or copy number.

In the 4,151 genes identified from the GWAS peaks, 2,363 harbored LoF mutations of which 864 possessed LoF mutations with minor allele frequency (MAF) > 5%. The accessions with LoF and non-LoF alleles for these 864 possessed LoF mutations with MAF > 5% provide the possibility to test the causality of these genes. In total, 61 genes exhibited a significant difference (false discovery rate [FDR] < 1%) in TE family expression level or copy number between accessions with LoF and non-LoF alleles, but no enriched GO terms were detected. Among these 61 genes, 30 exhibited elevated TE expression level or copy number; 30 displayed diminished TE expression level or copy number; and 1 showed dual roles, depending on the associated TE family (Figure 5M, Table S11).

## DISCUSSION

Understanding the genetic load as well as its causes during range expansion are long-standing questions, with important implications for human health ^55^, crop breeding ^2,56^, and conservation biology ^57^. Although TEs are a major component of the genomes of diverse species, the genetic load of TEs at the genome level in different natural populations remains largely unexplored. Moreover, the kind of demographic factors and molecular processes or mechanisms that determine the TE load need in-depth research ^58–60^.

The present study revealed the TE load variation among natural populations and pointed out the increased TE load at the expanding wave fronts. Although TEs are regarded as deleterious (Pasyukova et al. 2004; Barron et al. 2014; Hill et al. 2016), the deleterious effect of each specific TE is difficult to evaluate. By contrast, methods based on the evolutionary conservation across multispecies alignment, such as GERP ^61^, was previously used to classify the deleterious extent of base substitutions, which could not be used to evaluate the deleterious extent of TEs since TEs evolve fast and lack conservation among species. Here, based on the SFS of TEs in natural populations, we demonstrated that the deleterious effect of TEs varies probably between that of tnSNPs and dnSNPs. In addition, we adopted three approaches for inferring the deleterious effect of TEs, according to their insertion position and capacity of altering gene function and the functional importance of inserted or adjacent genes. These three methods enabled us to study the deleterious effects of TEs at high resolution in different natural populations.

More importantly, we clarified several factors that could affect TE load variation. Firstly, *N_e_* was identified a major contributor of the TE load variation and explained 62.0% of the variation in TE load along western and eastern expansion routes of Arabidopsis populations. Consistently, in *D. suzukii*, TE load is correlated with *N_e_* ^25^. Apparently, to maintain a relatively large population size, reducing or maintaining the deleterious mutations in natural populations, especially those of endangered species, would be an effective strategy in conservation biology. Secondly, consistent with a theoretical study, which pointed out that transposition rate also contributes to TE number variation ^20^, we found that higher transposition rate also contributed to higher TE load in the expanded populations. To infer the mutation rate of TEs in natural populations is difficult because of the purging of deleterious TEs. Here, we used intergenic TEs as well as youngest TEs, which are less likely to undergo selection-mediated purging. This approach revealed that the Yangtze River basin population has a higher transposition rate. Nevertheless, mutation accumulation lines (several generations of manipulated populations) could be used to estimate the mutation rate more accurately ^10,11^ in the future. Thirdly, positive selection or hitch-hiking effect could contribute to the accumulation of TEs in the expanded populations. TEs have been demonstrated to be adaptive in diverse species ^16,62–65^. It will be highly interesting to determine which TEs are under positive selection and which phenotypic traits are affected by these TEs.

The genetic architecture of TE load has been investigated in both Arabidopsis and Drosophila, based on TE copy number variation in TE families or whole genome ^18,25,66^. Here, to investigate the genetic architecture of natural transposition variation, besides considering the result of transposition as in previous studies, we studied both the initiation stage (TE expression) and final stage (TE copy number variation) of the TE transposition process. Transcription marks the initiation stage of TEs, particularly retrotransposons and autonomous transposons. It is crucial to identify determinants of TE expression, which are largely unknown. By contrast, TE copy number variation represents the final result of the TE transposition process. Thus, we addressed TE load variation from two different angles for the first time, and identified candidate TE-regulating genes and active TEs that might affect TE gain and loss rates. We demonstrated that *RDR2*, identified in the GWAS of TE expression level, was correlated with the total TE number at the genome level in natural populations, and the non-reference allele of *RDR2* occurred at high frequency in the Yangtze River basin population and potentially contributed to TE expansion.

Nevertheless, detailed functional analyses are needed to reveal the causal mutations responsible for TE load and epigenetic regulation, which are essential for understanding the mechanism of TE load variation. Besides TE expression level and copy number variation, approaches that focus on the DNA methylation variation of TEs could also identify potential TE load regulators. Based on the DNA methylation level of TEs, as determined in the 1001 genomes project of Arabidopsis, the causal genes of DNA methylation variation of TEs have been identified via GWAS ^67,68^. In addition, because of the repetitive nature of TEs and the inability to identify TEs by short-read sequencing, similar studies performed using long-read sequencing methods will be beneficial for the study of TE load variation mechanisms. Nevertheless, the results of this study and those of previous studies suggest that TE load is associated with many diverse molecular pathways. Similar to more than 7,000 human height-associated genomic segments discovered recently in the GWAS ^69^, the genomes of natural Arabidopsis populations potentially contain numerous TE load-associated variants. Overall, our study demonstrates the variation in TE load during Arabidopsis expansion and highlights the causes of TE load variation.

## ACKNOWLEDGMENTS

We thank Fei Lu (Institute of Genetics and Developmental Biology, Chinese Academy of Sciences) and Jinfeng Chen (Institute of Zoology, Chinese Academy of Sciences) for their valuable suggestions and discussions, and Fu-Min Zhang (Institute of Botany, Chinese Academy of Sciences) for helpful suggestions related to the analyses. This work was supported by the National Natural Science Foundation of China (31925004) and the Strategic Priority Research Program of the Chinese Academy of Sciences (XDB27010305).

## AUTHOR CONTRIBUTIONS

Y.-L.G. conceived the study; X.-H.H., and X.-M.N. performed the experiments; J.J., Y.-C.X, Z.-Q.Z., J.-F.C., X.-T.L., L.W., Y.Z., S.G. and Y.-L.G. analyzed and interpreted the data; J.J. and Y.-L.G wrote the paper with contribution from all authors.

## MATERIALS AND METHODS

### Plant material and high-throughput DNA sequencing

The paired-end resequencing data of 1,114 globally distributed non-reference accessions were obtained from four sources. The data of 810 accessions were obtained from the 1001 Genomes Project (NCBI SRP056687) ^26^, and those of 60 accessions were retrieved from the published data of Africa (ENA PRJEB19780) ^27^. Of the 244 accessions collected from China, 116 were sequenced by our laboratory and published previously (NCBI SRP062811) ^28^ (Table S1), and 128 were sequenced in this study (CNCB-NGDC, GSA: CRA008569). Coverage for each accession was over 10x.

The 128 Arabidopsis natural accessions sequenced in this study were collected from the Yangtze River basin and northwestern China (Table S2). DNA was extracted from leaves with the CTAB method. Paired-end sequencing libraries, with an insert size of approximately 350 bp, were constructed and sequenced on the Illumina HiSeq X Ten platform to generate 150 bp paired-end reads.

### Population structure

According to a previous study, the worldwide collection of Arabidopsis includes relicts (accessions that are distantly related to other accessions) and non-relicts ^26^. African accessions that are at least as divergent as relicts defined in the 1001 Genomes Project ^27^ were also treated as relicts. Two accessions from southwestern China, which are more divergent from non-relicts than those of relicts defined in the 1001 Genomes Project, were also treated as relicts (Figure S11A).

Non-relict accessions from the 1001 Genomes Project were classified into eight populations and an admixed group, as described previously ^26^. Because accessions from North America represent a newly colonized population ^70^, we designated these accessions as the North American population. Additionally, 38 accessions (including 11 accessions sequenced in this study) from northwestern China clustered with the Central Asia population of the 1001 Genomes Project (Figure S11B); together, they were grouped as the NW. China & C. Asia population. A total of 204 accessions from the Yangtze River basin (117 sequenced in this study) formed a cluster, which was designated as the Yangtze River basin population (Figure S11B). In total, all these 1,114 non-reference accessions were grouped into one relict, ten non-relict populations, and an admixed group.

### TE identification and feature analyses

TEPID ^17^ was employed for the detection of polymorphic TEs in 1,114 non-reference Arabidopsis accessions. Using the TE annotation of Col-0 (TAIR10) as a reference, the TE presence/absence calls for each accession were determined with tepid-map and tepid-discover algorithm, and the tepid-refine algorithm was further used to reduce false negative calls.

GEVA was used to estimate the age of polymorphic TEs with the parameters “--Ne 300000” and “--mut 7e-9”^32^. Full-length LTR TEs were annotated using LTRpred ^71^ in Col-0, and LTR similarity was calculated by LTRpred. CD-Search ^72^ was used to search for TE transposition related domains; a transposase for TIR transposons; RPA and helicase for Helitrons; reverse transcriptase and endonuclease for LINEs; and GAG, AP, RT, RNaseH, and INT for LTR TEs ^73^. The expression potential of TEs was measured using the published TE-transcript annotation data of Col-0 (TAIR10) ^34^, which was produced using TE-activated mutants. In the TE-transcript annotation, TEs were grouped into three categories (“Expressed and Annotated,” “Low expressed,” “.”) based on transcript abundance. The “Expressed and Annotated” and “Low expressed” categories were supported by reads, and reads were more abundant in “Expressed and Annotated” category than in the “Low expressed” category, while the “.” category had no read support. DNA methylation data were downloaded from NCBI (GSE43857), and the weighted methylation level of TEs was calculated as described previously ^74^.

### Evaluation of the deleterious effect of TEs

To determine the derived allele frequency spectrum of four-fold degenerate sites and deleterious mutations, only SNP sites with missing rate <10% were used. SNPs and indels were called using the GATK pipeline (GATK v2.1.8) ^75^ and annotated with SnpEff (version 4.3t) ^76^. Provean (Choi et al. 2012) was used to predict the deleterious effect of nSNPs against the NCBI nonredundant protein database. The nSNPs with score of Provean analysis ≤ −2.5 were defined as deleterious (dnSNPs), and those with score > −2.5 were defined as tolerated (tnSNPs). The LoF mutations (including stop-gain, splice site, and frameshift) were identified and filtered as described previously ^77^. However, three or more frameshift mutations found in the same gene of an accession were not excluded in the filtering step; only the frameshift mutations that restored the reading frame were filtered out. Ancestral state inference was based on the genome sequence alignment of Col-0 and its two close outgroups, *A. lyrata* (MN47) and *C. rubella* (MTE), using LASTZ. Alleles that matched the two outgroups were defined as ancestral alleles. Alleles different from the alleles of two outgroups, which were identical, were defined as derived alleles.

To determine the derived allele frequency spectrum of TEs, only polymorphic TE sites with missing rate <10% were used. The ancestral state inference of TEs was based on the genome sequence alignment of Col-0, *A. lyrata*, and *C. rubella* using AnchorWave ^78^, which is more sensitive in making TE presence/absence calls than LASTZ. TEs absent in the two outgroups but present in *A. thaliana* were defined as derived.

To study the effect of TEs on gene expression, the normalized transcriptome data of 413 accessions were downloaded from NCBI (GSE80744) ^39^. To calculate fold-change in the expression level of each gene with polymorphic TEs in or nearby, the gene expression level in accessions with TEs was normalized relative to that in accessions without TEs.

To study the effect of TEs on alternative splicing (AS), the transcript model of Col-0 was downloaded from TAIR (Araport11). SUPPA2 ^79^ was used to identify seven types of AS events (skipping exon [SE], alternative 5’splice sites [A5], alternative 3’ splice sites [A3], mutually exclusive exons [MX], retained introns [RI], alternative first exons [AF], alternative last exons [AL]). The TPM value of each transcript was measured with TopHat ^80^ and Salmon ^81^, and the RNA-seq data downloaded from NCBI (GSE80744) were used to calculate the TPM value of each transcript in 413 accessions. The PSI value of each AS event was calculated with SUPPA2 based on the TPM value of each accession. The PSI value indicates splicing efficacy; the larger the PSI value, the lower the splicing efficacy. To calculate fold-change in the PSI value of each gene with polymorphic TEs in or nearby, the gene PSI value in accessions with TEs was normalized relative to that in accessions without TEs.

The importance of gene function was measured based on sequence conservation and known mutant phenotypes. To evaluate gene sequence conservation, the dN/dS ratios of the orthologous genes of *A. thaliana* and *A. lyrata* were calculated with KaKs_calculator 3.0 ^82^. Genes were categorized into four bins, according to the dN/dS quartiles; the first quartile was the most constrained, and the fourth quartile was the least conserved. The mutant phenotypes of 2,400 genes were grouped into four categories (essential, morphological, cellular-biochemical, and conditional) according to their effect, as reported previously ^43^.

### Evaluation of the synergistic epistasis of TEs

To evaluate the synergistic epistasis of deleterious TEs, only TEs with frequency lower than 1%, which are more likely to be deleterious, were used. TEs in pericentromeric regions, which were determined based on genome-wide DNA accessibility analysis using DNase I sensitivity assays ^83^, were ruled out because pericentromeric regions have extensive LD and would likely induce a bias. PLINK (v1.90b4) ^84^ was used to calculate the correlation coefficient (r) of each TE pair, and the raw value of LD was back-calculated using the equation below:

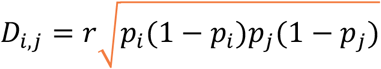

where *p_i_* and *p_j_* are the frequency of TEs *i* and *j*, respectively.

Each TE pair was categorized according to its physical distance (on the same chromosome or on different chromosomes) and deleterious effect, and the mean LD of TE pairs in each category was calculated. At least five TE pairs were required to be included in each category.

### Geographical distance, genetic diversity, and TE load calculation

The Haversine distance of each accession to the putative non-relict origin predicted previously was calculated using the “geosphere” package of R, and the mean distance of all accessions within a population was used as the distance of this population to the origin. Genetic diversity (π) was calculated using VCFtools ^85^ in non-overlapping 10 kb windows for each population. To estimate the TE load, only polymorphic TE sites with missing rate <10% were used. Derived polymorphic TE counts were used as load proxies and compared among ten non-relict populations.

### Selective sweep region identification

OmegaPlus (version 3.0.3) ^86^ was used to identify selective sweep regions in the Yangtze River basin and NW. China & C. Asia populations. OmegaPlus is based on LD, and the ω statistic was computed at 10 kb intervals with the parameters “-minwin 10000” and “-maxwin 100000.” The top 5% regions with high ω value were defined as selective sweep regions. To test the enrichment of TEs in selective sweep regions, the derived allele counts per base-pair of four-fold degenerate sites and TE sites were compared between selective sweep and non-sweep regions. Pericentromeric regions were removed from sweep and non-sweep regions when the derived allele count per base-pair was calculated since these regions exhibit high TE density and low gene density, which could bias the results.

### TE expression analysis

Because TEs are highly repetitive in nature, exhibit polymorphic insertions, and display co-transcription with genes, the quantification of TE expression using short-read data has been a challenging task for a long time. Recently, many tools have been developed to quantify TE expression using RNA-seq data, at both the family and locus levels ^87^. However, accurate counting at the locus level is still challenging because of ambiguous mapping, especially for young TEs. Therefore, we focused on quantifying TE family expression using a modified version of TEtranscripts pipeline ^88,89^, which largely excludes the transcripts that co-transcribed with genes.

High-coverage RNA-seq data of 414 natural Arabidopsis accessions (including Col-0) were downloaded from NCBI (GSE80744). RNA-seq reads were mapped to TAIR10 using STAR ^90^ with the parameters “— winAnchorMultimapNmax 100 --outFilterMultimapNmax 100”. A modified version of TEtranscripts ^88,89^ was used to quantify TE expression level. Two output files (unique mapping and multiple mapping) for each TE family and individual TEs were generated with the parameters “-mode uniq” and “-mode multi,” respectively. In the “uniq” mode, only reads uniquely mapped to TEs were counted, and in the “multi” mode, all reads mapped to TEs were counted. The multiple mapping file was used to measure the transcription level of each TE family, and the unique mapping file was used to quantify the transcripts of each individual TE. The scaling factor for each sample was calculated using edgeR ^91^ with the RLE method and was then used to normalize the expression data of each TE family and individual TE. To compare the expression of 18 TE superfamilies, the expression level of each superfamily was further normalized relative to the corresponding superfamily size. The unique mapping rate of each TE was calculated based on raw read counts and defined as the percentage of uniquely mapped reads relative to all mapped reads.

### GWAS analysis

Biallelic SNPs with MAF > 5% and missing rate <10% were used in GWAS analysis. In the TE family expression GWAS analysis, expression levels of 319 expressed TE families were used as phenotypes. In the TE family copy number GWAS analysis, the copy number of 318 TE families with polymorphic TEs was used as phenotypes. GWAS was performed with EMMAX (linear mixed model) for each phenotype using principal components and kinship matrix to control for population structure ^92^. Significant thresholds were set based on Bonferroni correction (0.01/number of passed SNPs); candidate interval was determined based on the lead SNP of a peak and the SNPs linked to the lead SNP (r^2^ > 0.2), and pairwise LD was calculated using PLINK (v1.90b4) ^84^. The candidate intervals were then intersected with gene annotation (Araport11), reference TE annotation (TAIR10), and polymorphic TEs to identify the causal element. GO enrichment analysis was conducted with agriGO ^93^.

### Analysis of the transposition potential of candidate causal TEs

To estimate the capacity of TEs to produce transposition intermediates, the Oxford Nanopore Technology (ONT) long-read VLP DNA-seq data of Col-0 were retrieved from NCBI (GSE128932) ^50^. The long-read data of two genotypes (wild type and *ddm1* mutant) were mapped to the reference genome (TAIR10) using Minimap2 ^94^. The number of reads mapped to annotated TEs (TAIR10) was calculated using featureCounts ^95^. LTR TEs with at least one read were regarded as LTR TEs capable of producing transposition intermediates. To calculate the translational efficiency of TE genes, polysomal RNA-seq and RNA-seq data of the *ddm1* mutant (Col-0 background) were retrieved from NCBI (GSE128932) ^50^. The translational efficiency of TE genes (Araport 11) was calculated as the ratio of the FPKM value of polysomal RNA to that of total RNA.

### Analysis of LoF mutations in candidate genes

To validate the causality of candidate genes identified by GWAS, we utilized the natural LoF mutations of these genes and determined if their respective phenotypes (expression level or copy number) differed with the PAV of LoF allele. Among the LoF alleles with MAF > 5%, Mann-Whitney U test was used to test the difference in TE expression level or copy number between LoF and non-LoF alleles. After multiple test correction, the LoF alleles with FDR < 0.01 were defined as significant. Candidate genes with significant LoF alleles were designated as causal genes.

### Statistical analyses

All statistical analyses were performed in R (http://www.r-project.org/).

## Data accessibility

The raw sequence data reported in this paper have been deposited in the Genome Sequence Archive in National Genomics Data Center, China National Center for Bioinformation (GSA: CRA008569) that are publicly accessible at https://ngdc.cncb.ac.cn/gsa.

