## Supplemental Information for "Forces driving transposable element load variation during Arabidopsis range expansion"

Beijing 100093, China

PH +86-10-62836298; FX +86-10-62590843

EM

Running title: The genetic load of transposons

Key words: *Arabidopsis thaliana*, genetic architecture, genetic load, natural variation, transposable elements

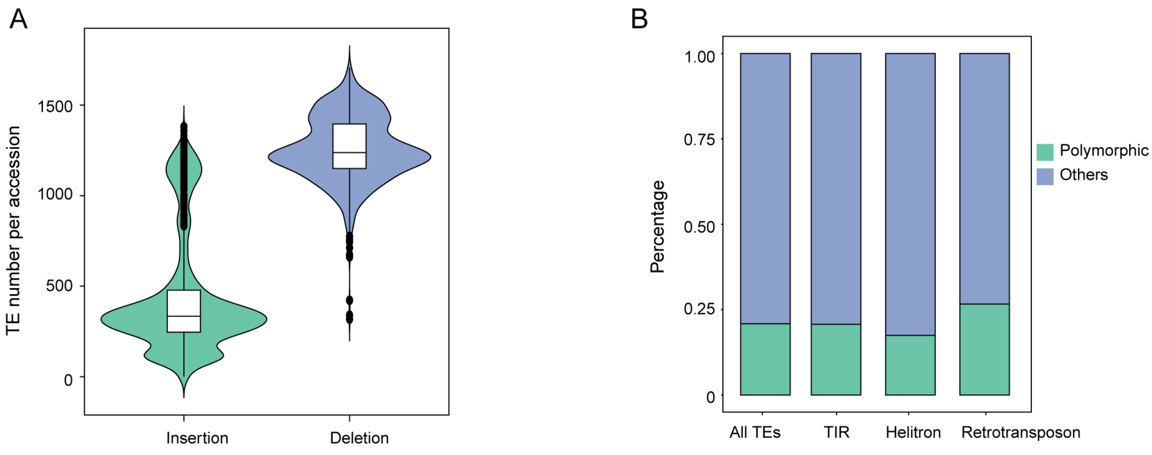


Figure S1. TEs identified in 1,115 accessions. (A) Number of TE insertion and deletion per accession comparing to reference genome. (B) Percentage of polymorphic TEs for all TEs and different type TEs in Col-0. Others indicate TEs that are not identified as absence in any accessions, including fixed TEs and TEs without reads mapping in flanking regions at some accessions.

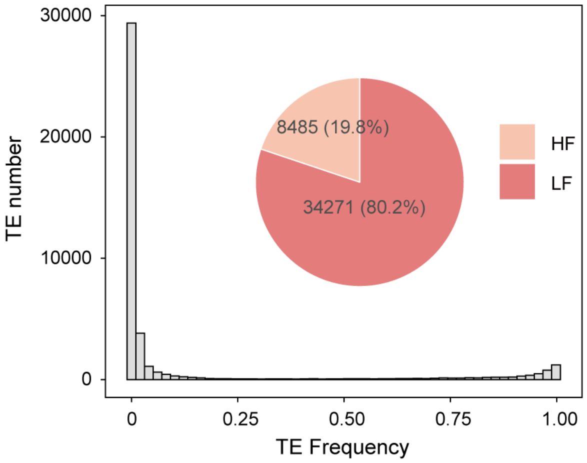


Figure S2. Frequency distribution of polymorphic TEs. HF: high frequency TEs (frequency ≥5%); LF: low frequency TEs (frequency <5%).

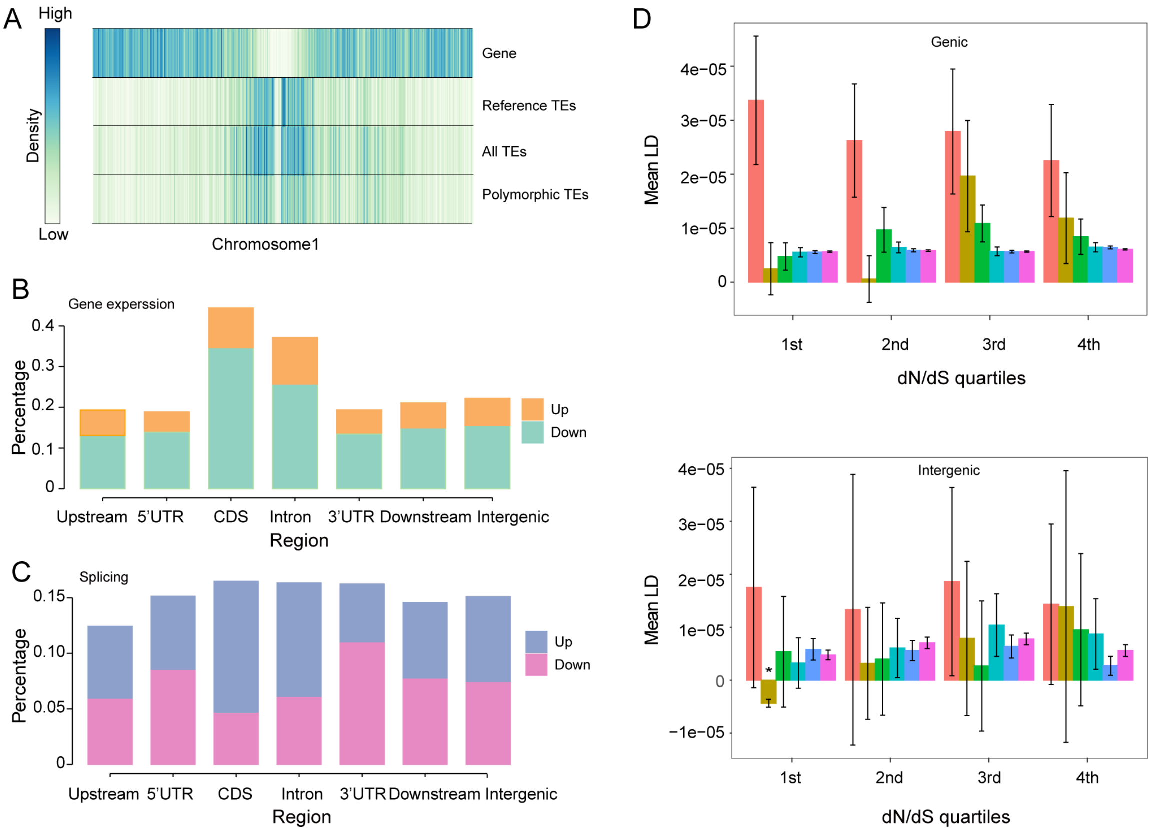


Figure S3. Classifying the deleterious extent of TEs. (A) Genomic distribution of genes and TEs, taking chromosome1 as an example. All TEs, polymorphic TEs and non-polymorphic reference TEs. (B) Percentage of TEs located in different genomic regions that are associated with more than two-fold gene expression changes. Up/Down, upregulate/downregulate gene expression level. (C) Percentage of TEs located in different genomic regions that are associated with more than two-fold PSI value changes. Larger PSI indicates lower splicing efficiency. Up/Down, inhibit/promote splicing. (D) Mean LD of TE pairs located within genic or intergenic regions of genes with different conservation categories (dN/dS quartiles, the first quartile is the most constrained, the last quartile is the least constrained). The * denotes significant negative LD of which the 95% confidence interval is completely negative. Color of bars denotes different physical distance between TE pairs.

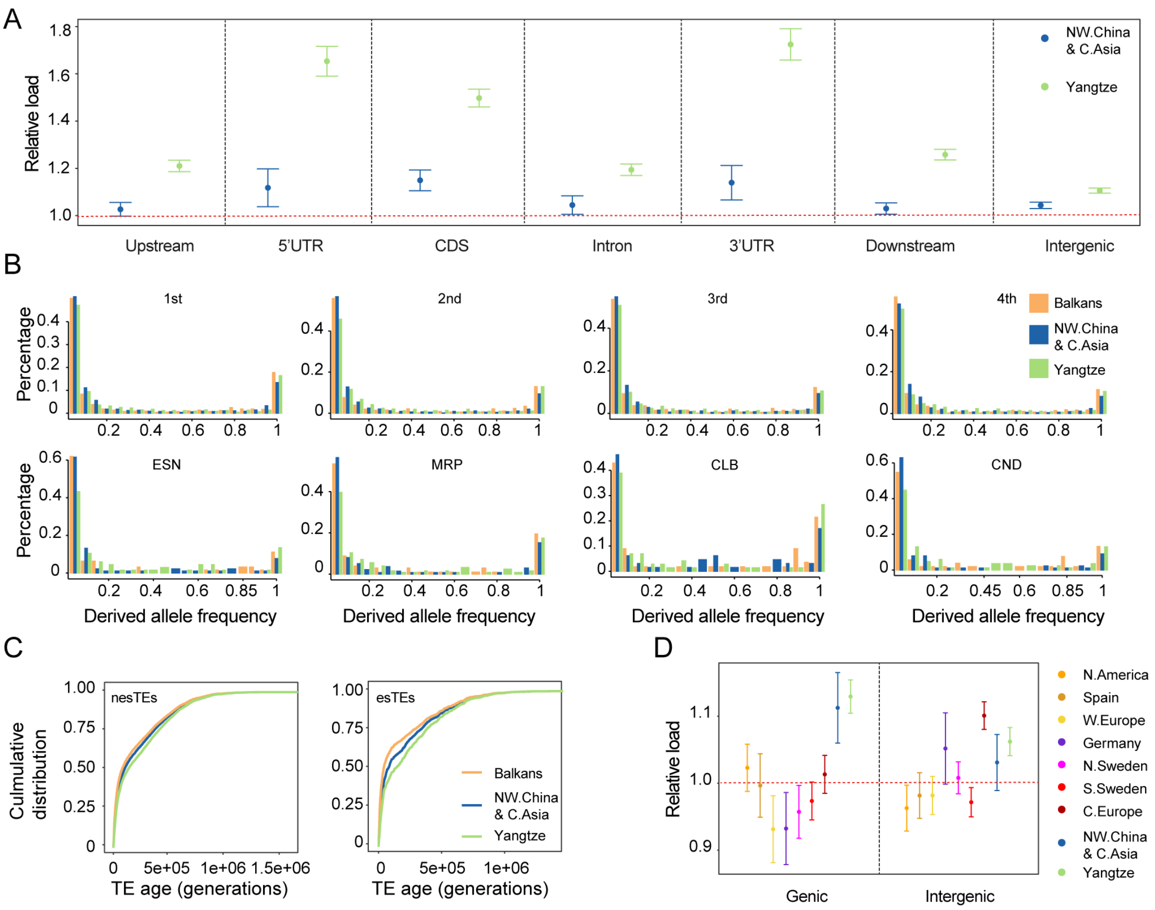


Figure S4. Causes and consequences of TE load accumulation in Yangtze River basin population. (A) Load of TEs with different deleterious effects as classified in Figure 2. For each variants categories, the mean number of derived alleles of Balkans was used as standard to calculate the relative load. Error bars represent 95% confidence interval. (B) Comparison of the SFS of different TEs in three populations. TEs were classified based on the function essentiality of their closest gene. The upper functional categories are based on the evolutionary conservation of genes, the bottom functional categories are based on the known mutant phenotype of genes. (C) Age distribution of TEs with different deleterious effects in three populations. (D) Comparison of recent TE load (the top 5% youngest TEs) located in different regions among non-relict populations.

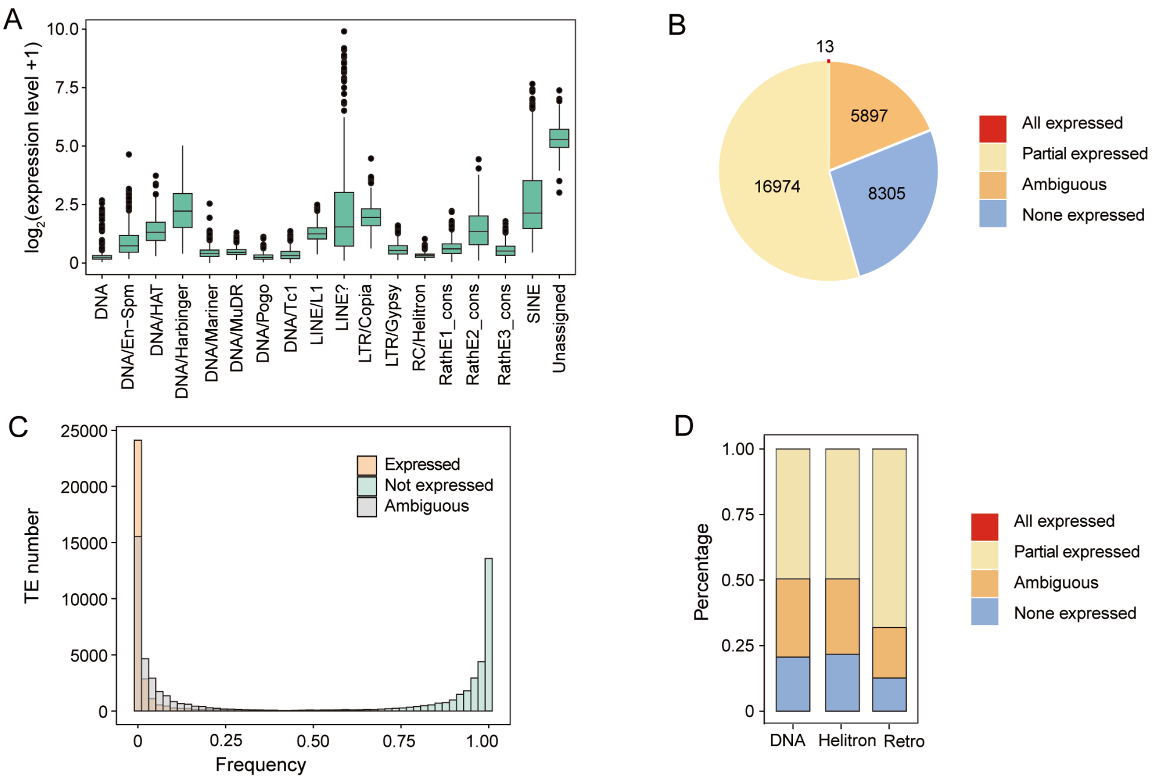


Figure S5. TE expression in different accessions at family level and locus level. (A) Expression level of 18 TEs superfamilies. Y axis represents the expression level of TE superfamilies, we use $\log_2$ (expression level + 1) as some superfamilies are not expressed in some accessions. (B) Percentage of expressed TE. All expressed, TEs with unique mapping reads in all (414) accessions; Partially expressed, TEs with unique mapping reads in at least one accession; None expressed, TEs without any reads in any accessions; Unresolved, TEs with multiple mapping reads but no unique mapping reads. (C) TE number distribution of different expression frequency. Frequency = expressed accessions (or not expressed, others)/all accessions. Expressed, TEs with unique mapping reads; Not expressed, TEs have no reads mapping; Unresolved, TEs with multiple mapping reads but no unique mapping reads. (D) Percentage of expressed TE for different TE types. Retro, retrotransposon.

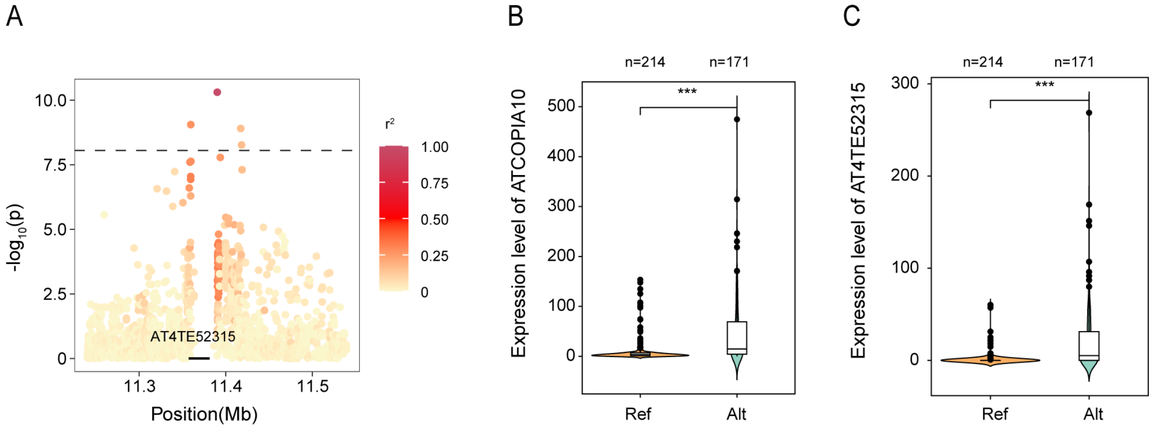


Figure S6. GWAS of ATCOPIA10 family expression level. (A) Local Manhattan plot of ATCOPIA10 family expression level. $r^2$ represents the LD between lead SNP and other SNPs. (B) ATCOPIA10 family expression level in two alleles (grouped by the lead SNP). ***, $p < 0.001$. Mann-Whitney U test was used for significance test. Ref, reference allele; Alt, non-reference allele. (C) AT4TE52315 expression level in two alleles (grouped by lead SNP). Mann-Whitney U test.

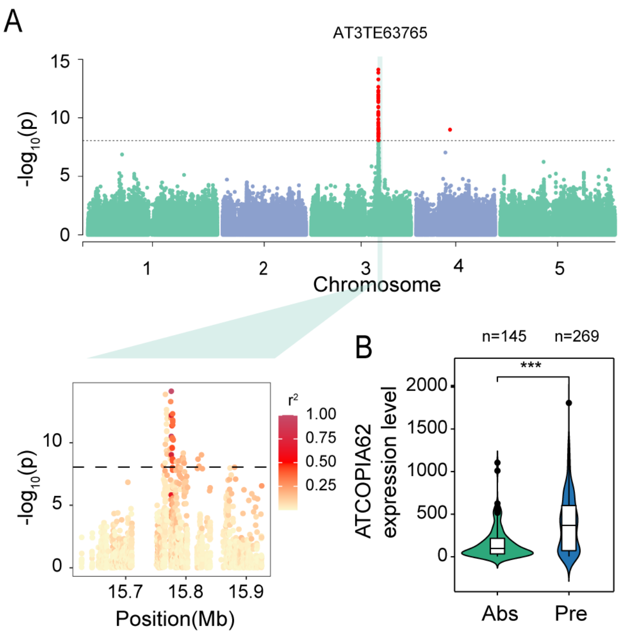


Figure S7. GWAS of ATCOPIA62 family expression level. (A) Manhattan plot and local Manhattan plot of ATCOPIA62 family expression level. $r^2$ represents the LD between lead SNP and other SNPs. (B) Expression level of ATCOPIA62 with AT3TE63765 presence and absence variation. Absence, TE absence allele; Presence, TE presence allele. ***, $p < 0.001$. Mann-Whitney U test was used for significance test.

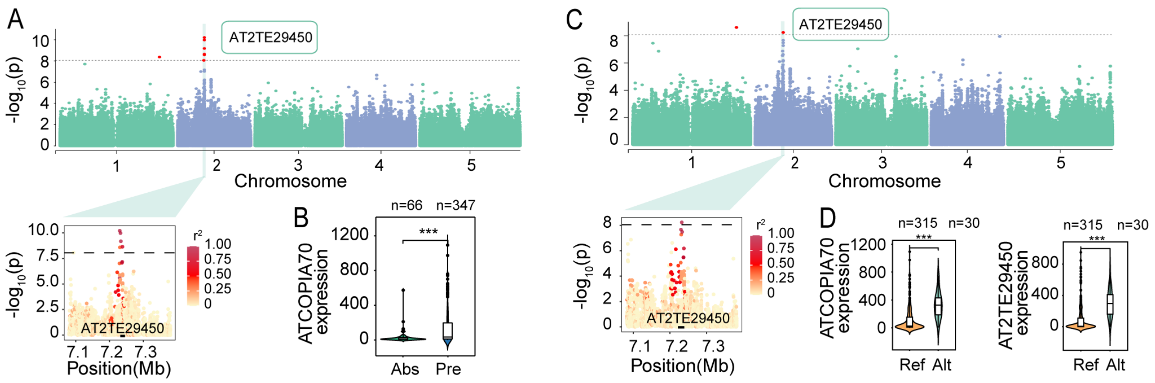


Figure S8. GWAS on the expression level of ATCOPIA70 family. (A) Manhattan plot and local Manhattan plot of GWAS for ATCOPIA70 family expression GWAS. $r^2$ represents the LD between lead SNP and other SNPs. (B) Expression level of ATCOPIA70 family with AT2TE29450 presence and absence variation. Abs, TE absence allele; Pre, TE presence allele. ***, $p < 0.001$. Mann-Whitney U test was used for significance test. (C) Manhattan plot and local Manhattan plot of ATCOPIA70 expression level GWAS based only on accessions with AT2TE29450. (D) ATCOPIA70 expression level and AT2TE29450 expression level in two alleles (grouped by lead SNP). Ref, reference allele; Alt, non-reference allele.

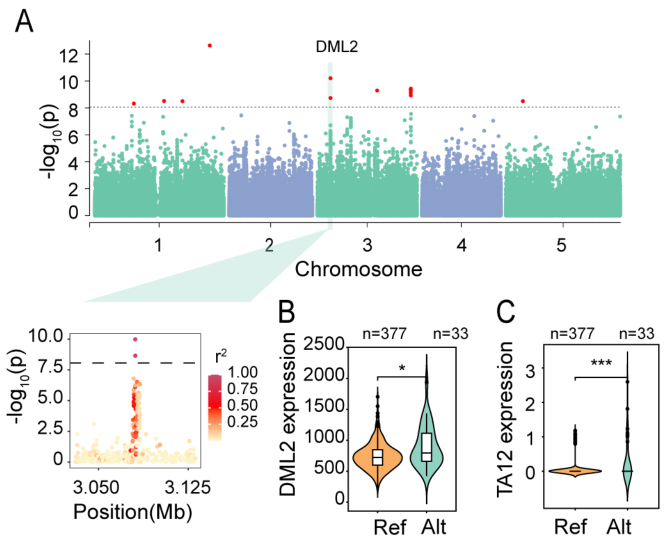


Figure S9. GWAS results of TA12 family expression level. (A) Manhattan plot and local Manhattan plot of TA12 family expression level. $r^2$ represents the LD between lead SNP and other SNPs. (B) *DML2* expression level of different alleles. ***, *p* < 0.001; *, *p* < 0.01. Mann-Whitney U test was used for significance test. Ref, reference allele; Alt, non-reference allele. (C) TA12 family expression level of different alleles.

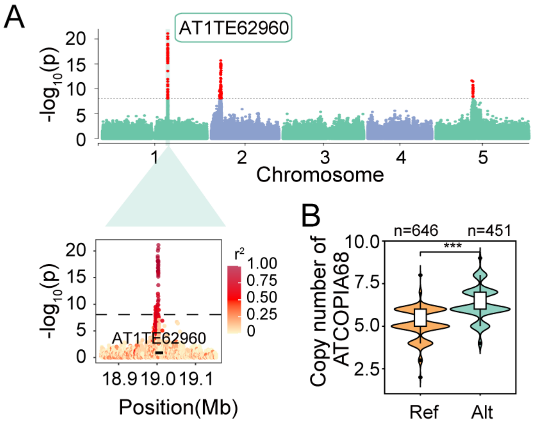


Figure S10. GWAS results of ATCOPIA68 copy number. (A) Manhattan plot and local Manhattan plot of GWAS for ATCOPIA68 family copy number. (B) ATCOPIA68 copy number in two alleles (grouped by lead SNP). ***, $p < 0.001$. Mann-Whitney U test was used for significance test.

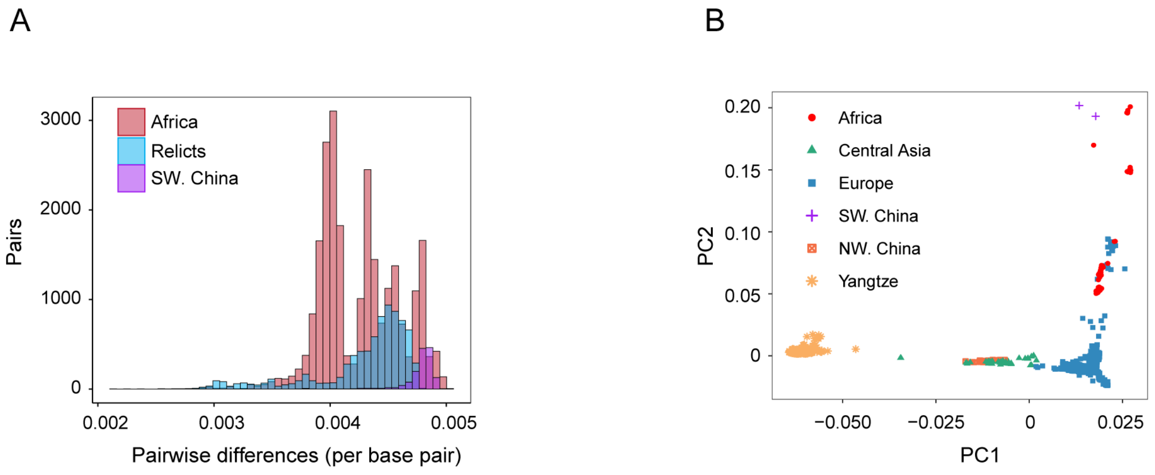


Figure S11. Population structure of 1,115 accessions. (A) Per base pairwise differences between 2 accessions from southwestern China (SW. China) and non-relict accessions. The per base pairwise difference between relicts defined in 1001 Genomes Project and non-relicts, Africa accessions and non-relicts are also showed. (B) PCA of 1,115 natural accessions based on SNP. Europe represents accessions from 1001 Genomes Project, in which Central Asia accessions were extracted to denote their relationship with accessions from northwestern China. SW. China, southwestern China. NW. China, northwestern China. Yangtze, accessions from Yangtze River basin.

Table S1. Summary of the published natural accessions used in this study.

| Source | Population | accessions |
|---|---|---|
| 1001 | North America | 1612,1622,1651,1652,1676,1684,1756,1757,1793,1797,1820,1829,1834,1835,1851,1852,1872,1890,1925,2016,2017,2031,2053,2057,2108,2171,2191,2212,2239,2240,2278,2285,2286,506,544,546,628,630,6739,6740,6744,6749,6750,6805,6806,680,6814,681,685,687,6926,6927,7248,728,7350,7356,7358,7359,7383,7416,7475,7514,7515,7525,7529,7530,7568,7717,7757,7767,801,8037,8057,8171,8233,8246,8464,8483,853,854,867,868,870,9027,932 |
| 1001 | NW. China & C. Asia | 14312,14313,14314,14315,14318,14319,15560,18694,18696,6929,6931,6938,7183,7323,763,766,772,8424,9125,9128,9130,9131,9133,9134,9609,9611,9612,9616,9620,9621,9625,9626,9629,9630,9631,9635,9636,9639,9640,9641 |
| 1001 | Central Europe | 10020,10027,15591,15593,19949,19950,19951,403,410,424,428,430,5837,5874,5890,5893,5907,5921,5950,5984,5993,6008,6296,6390,6396,6424,6445,6903,6919,6951,6956,6957,6975,6976,6979,6984,7025,7067,7120,7177,7186,7203,7207,7236,7273,7347,7372,7396,7411,7413,7520,7521,8235,8236,8284,8285,8290,8311,8365,8386,8419,9644,9664,9665,9669,9670,9671,9672,9673,9677,9678,9679,9682,9683,9684,9685,9686,9689,9690,9692,9693,9694,9695,9696,9730,9731,9732,9733,9756,9768,9769,9770,9771,9772,9774,9775,9776,9777,9778,9779,9780,9781,9782,9784,9785,9786,9787,9788,9789,9791,9792,9793,9794,9795,9796,9797,9798,9799,9800,9801,9802,9803,9804,9805,9806,9807,9808,9812,9815 |
| 1001 | Germany | 5486,5748,5800,6180,6252,6268,6276,6915,6940,6945,6973,6997,7002,7008,7014,7036,7062,7102,7117,7119,7133,7143,7161,7165,7192,7199,7202,7208,7258,7276,7282,7287,7322,7419,7461,7516,8238,8312,9442,9476 |
| 1001 | Balkans | 7077,9067,9069,9070,9075,9078,9081,9084,9085,9089,9091,9095,9100,9102,9103,9104,9105,9106,9111,9113,9114,9115,9121,9613,9647,9649,9653,9656,9660,9698,9699,9700,9701,9703,9705,9708,9710,9711,9712,9713,9717,9720,9722,9726,9755 |
| 1001 | North Sweden | 1254,1257,1552,5856,5860,6009,6010,6011,6012,6013,6016,6017,6025,6046,6064,6069,6070,6071,6153,6154,6163,6166,6169,6172,6173,6174,6177,6184,6209,6210,6214,6216,6217,6218,6220,6221,6235,6238,6240,6244,6900,6901,6913,6917,6918,6968,6969,8227,8351,8376,9321,9323,9332,9363,9371,9386,9388,9427,9433 |
| 1001 | Admixed | 10022,10023,1313,1317,265,5720,5757,5768,5772,5784,5811,6039,6042,6086,6258,6434,6830,6907,6920,6932,6981,6992,7003,7028,7058,7061,7096,7111,7126,7127,7147,7163,7169,7181,7209,7213,7288,7305,7314,7316,7342,7353,7373,7384,7394,7404,8256,8334,8366,9058,9336,9343,9381,9382,9383,9394,9506,9508,9513,9523,9527,9529,9530,9536,9548,9551,9552,9558,9559,9561,9565,9576,9579,9581,9591,9595,9596,9663,9743,9754,9764,9783,9831,9835,9839,9849,9881,9890,9892,9894,9933 |
| 1001 | Relicts | 6911,9533,9542,9543,9545,9549,9550,9554,9555,9574,9583,9598,9600,9606,9832,9837,9871,9879,9887 |
| 1001 | South Sweden | 1002,1006,1061,1062,1063,1066,1070,1158,1166,5830,5831,5832,5836,5865,5867,6019,6020,6021,6022,6023,6024,6034,6036,6038,6040,6041,6073,6074,6076,6077,6085,6087,6088,6090,6091,6092,6094,6095,6096,6097,6098,6099,6100,6101,6102,6104,6105,6106,6107,6109,6111,6112,6113,6114,6115,6118,6119,6122,6123,6124,6125,6126,6128,6131,6132,6133,6134,6136,6137,6138,6140,6141,6142,6145,6148,6149,6150,6151,6188,6189,6191,6192,6193,6194,6195,6201,6202,6203,6242,6284,6413,6974,7013,7346,7354,8222,8230,8231,8234,8237,8240,8241,8242,8247,8249,8258,8259,8283,8306,8307,8326,8335,8369,8422,8426,8427,9057,9339,9352,9353,9369,9370,9380,9390,9391,9392,9399,9402,9404,9405,9407,9408,9409,9412,9413,9416,9421,9436,9451,9452,9453,9454,9455,9470,9471,9481,991,992,997 |
| 1001 | Spain | 6933,6961,6971,7081,7327,7328,8264,8357,9507,9509,9510,9511,9512,9514,9515,9518,9519,9520,9521,9522,9524,9525,9526,9531,9532,9534, |

| | | |
|---|---|---|
| | | 9535,9537,9540,9541,9544,9546,9547,9553,9556,9557,9560,9562,9564,9567,9568,9573,9577,9582,9584,9586,9587,9588,9589,9593,9594,9597,9601,9602,9817,9819,9820,9821,9822,9825,9833,9834,9836,9840,9841,9843,9845,9846,9848,9850,9852,9855,9868,9870,9873,9874,9876,9878,9880,9888,9891,9895,9898,9900,9901,9902 |
| 1001 | Western Europe | 108,139,159,350,351,4807,4840,4857,4939,4958,5210,5249,5253,5276,5279,5349,5353,5395,5577,5644,5651,5717,5718,5726,5741,5776,5779,5798,6108,6904,6908,6923,6943,6944,6958,6959,6960,6966,6967,6986,7026,7064,7071,7092,7130,7217,7387,8214,8243,8244,8297,8337,88,9569,9571,9578,9585,9590,9599,9809,9826,9827,9847,9851,9853,9854,9875 |
| Africa | Relicts | Agl0,Agl1,Agl2,Agl3,Agl5,Agl9,Ait9,Ait14,Arb0,Arb2,Azr0,Azr5,Azr7,Azr11,Azr13,Azr16,Bab0,Bab3,Bba0,Bba2,Bbe0,Elh2,Elh10,Elh15,Elh20,Elh23,Elh27,Elh33,Elh39,Elh46,Elk1,Elk3,Elk20,Elk28,IFr0,Ifr3,Ifr4,Ifr6,Ket10,Ket12,Khe0,Khe32,Meh0,Meh4,Meh7,Oua0,Set0,Set6,Tah0,Tah4,Tanz-1,Taz0,Taz11,Taz16,Taz18,Til2,Tiz0,Tiz7,Zin4,Zin9 |
| China | NW. China & C. Asia | 39-16,40-2,41-10,41-11,42-3,42-8,43-12,43-7,45-20,45-23,46-28,46-31,48-1,50-3,50-5,51-1,51-4,51-8,52-10,52-13,53-18,53-19,54-5,54-6,55-12,55-13,55-22 |
| China | Relicts | 36-31,87 |
| China | Yangtze | 10-1,10-3,10-5,11-4,12-8,13-13,1-32,13-5,14-62,15-11,17-2,17-5,18-4,18-7,1-8,19-33,20-2,20-36,23-19,23-34,24-10,25-16,2-5,26-5,27-8,27-9,28-6,29-8,30-1,30-6,31-26,32-28,3-2,33-46,33-4,34-16,3-4,35-10,35-1,37-10,38-2,38-4,4-15,5-15,57-1,58-7,59-12,60-1,61-1,62-1,6-28,63-1,64-5,65-1,66-3,67-1,68-1,69-7,6-9,70-1,71-5,72-2,7-2,73-5,74-14,75-6,76-3,76-4,76-5,77-2,77-3,78-4,78-5,79-2,79-4,8-17,82-1,82-3,83-2,83-5,84-3,84-4,85-3,85-5,86,8-6,9-5 |

Note: NW. China & C. Asia, northwestern China and central Asia population; Yangtze, Yangtze River basin population.

Table S2. Summary of the newly-sequenced 128 natural accessions in this study.

| Population | Accession | Region | Latitude | Longitude |
|---|---|---|---|---|
| NW. China & C. Asia | 39-6 | Qinghe, Xinjiang | 46.75 | 90.27 |
| NW. China & C. Asia | 39-7 | Qinghe, Xinjiang | 46.75 | 90.27 |
| NW. China & C. Asia | 41-7 | Altay, Xinjiang | 47.59 | 88.77 |
| NW. China & C. Asia | 42-4 | Emin, Xinjiang | 46.83 | 84.18 |
| NW. China & C. Asia | 43-1 | Qinghe, Xinjiang | 46.74 | 90.34 |
| NW. China & C. Asia | 45-25 | Qinghe, Xinjiang | 46.75 | 90.34 |
| NW. China & C. Asia | 46-27 | Qinghe, Xinjiang | 46.74 | 90.34 |
| NW. China & C. Asia | 50-4 | Fuyun, Xinjiang | 47.36 | 89.65 |
| NW. China & C. Asia | 53-21 | Altay, Xinjiang | 47.77 | 88.38 |
| NW. China & C. Asia | 54-4 | Altay, Xinjiang | 47.73 | 88.22 |
| NW. China & C. Asia | 55-7 | Altay, Xinjiang | 47.73 | 88.22 |
| Yangtze | 11-1 | Jinggangshan, Jiangxi | 26.75 | 114.3 |
| Yangtze | 11-5 | Jinggangshan, Jiangxi | 26.75 | 114.3 |
| Yangtze | 12-11 | Jiujiang, Jiangxi | 29.59 | 115.91 |
| Yangtze | 12-3 | Jiujiang, Jiangxi | 29.59 | 115.91 |
| Yangtze | 13-10 | Nanfeng, Jiangxi | 26.99 | 116.24 |
| Yangtze | 14-5 | Xinjian, Jiangxi | 28.74 | 115.74 |
| Yangtze | 16-62 | Dongyang, Zhejiang | 29.08 | 120.43 |
| Yangtze | 17-11 | Jiande, Zhejiang | 29.54 | 119.49 |
| Yangtze | 18-5 | Kaihua, Zhejiang | 29.11 | 118.38 |
| Yangtze | 19-61 | Lin'an, Zhejiang | 30.05 | 119.91 |
| Yangtze | 19-66 | Lin'an, Zhejiang | 30.05 | 119.91 |
| Yangtze | 20-21 | Tonglu, Zhejiang | 29.69 | 119.67 |
| Yangtze | 21-42 | Lin'an,Zhejiang | 30.05 | 119.91 |
| Yangtze | 21-54 | Lin'an, Zhejiang | 30.05 | 119.91 |
| Yangtze | 21-64 | Lin'an, Zhejiang | 30.05 | 119.91 |
| Yangtze | 2-16 | Qianshan, Anhui | 30.75 | 117.62 |
| Yangtze | 2-1 | Qianshan, Anhui | 30.75 | 117.62 |
| Yangtze | 22-10 | Pan'an, Zhejiang | 28.96 | 120.37 |
| Yangtze | 22-20 | Pan'an, Zhejiang | 28.96 | 120.37 |
| Yangtze | 23-40 | Nanjing, Jiangsu | 32.05 | 118.83 |
| Yangtze | 24-14 | Wuchang, Hubei | 30.52 | 114.47 |
| Yangtze | 24-20 | Wuchang, Hubei | 30.52 | 114.47 |
| Yangtze | 25-30 | Beibei, Chongqing | 29.79 | 106.48 |
| Yangtze | 25-34 | Beibei, Chongqing | 29.79 | 106.48 |
| Yangtze | 26-1 | Tongliang, Chongqing | 29.82 | 106.06 |
| Yangtze | 26-3 | Tongliang, Chongqing | 29.82 | 106.06 |
| Yangtze | 27-5 | Wenxian, Gansu | 32.72 | 105.12 |
| Yangtze | 28-10 | Mianxian, Shaanxi | 33.15 | 106.75 |
| Yangtze | 28-17 | Mianxian, Shaanxi | 33.15 | 106.75 |
| Yangtze | 29-50 | Chenggu, Shaanxi | 32.93 | 107.21 |
| Yangtze | 29-6 | Chenggu, Shaanxi | 32.93 | 107.21 |
| Yangtze | 30-7 | Hong'an, Hubei | 31.28 | 115.02 |

| | | | | |
|---|---|---|---|---|
| Yangtze | 3-1 | Qingyang, Anhui | 30.62 | 117.76 |
| Yangtze | 32-15 | Hanyang, Hubei | 30.46 | 114.09 |
| Yangtze | 32-27 | Hanyang, Hubei | 30.46 | 114.09 |
| Yangtze | 33-53 | Zhangjiajie, Hunan | 29.41 | 110.44 |
| Yangtze | 34-11 | Yuanling, Hunan | 28.52 | 110.72 |
| Yangtze | 34-2 | Yuanling, Hunan | 28.52 | 110.72 |
| Yangtze | 35-3 | Yinjiang, Guizhou | 27.94 | 108.61 |
| Yangtze | 37-3 | Xinyang, Henan | 32.05 | 113.98 |
| Yangtze | 37-8 | Xinyang, Henan | 32.05 | 113.98 |
| Yangtze | 38-3 | Yichang, Hubei | 31.1 | 110.95 |
| Yangtze | 4-17 | Shexian, Anhui | 29.51 | 118.23 |
| Yangtze | 4-7 | Shexian, Anhui | 29.51 | 118.23 |
| Yangtze | 5-23 | Taihu, Anhui | 30.28 | 117.18 |
| Yangtze | 57-6 | Dongxiang, Jiangxi | 28.23 | 116.59 |
| Yangtze | 57-7 | Dongxiang, Jiangxi | 28.23 | 116.59 |
| Yangtze | 58-2 | Dongxiang, Jiangxi | 28.28 | 116.61 |
| Yangtze | 58-5 | Dongxiang, Jiangxi | 28.28 | 116.61 |
| Yangtze | 59-2 | Xinjian, Jiangxi | 28.75 | 115.82 |
| Yangtze | 59-4 | Xinjian, Jiangxi | 28.75 | 115.82 |
| Yangtze | 60-2 | Nanchang, Jiangxi | 28.78 | 115.84 |
| Yangtze | 60-7 | Nanchang, Jiangxi | 28.78 | 115.84 |
| Yangtze | 61-2 | Pengze, Jiangxi | 29.86 | 116.52 |
| Yangtze | 61-5 | Pengze, Jiangxi | 29.86 | 116.52 |
| Yangtze | 62-2 | Pengze, Jiangxi | 29.92 | 116.57 |
| Yangtze | 62-3 | Pengze, Jiangxi | 29.92 | 116.57 |
| Yangtze | 63-2 | Pengze, Jiangxi | 29.88 | 116.52 |
| Yangtze | 63-5 | Pengze, Jiangxi | 29.88 | 116.52 |
| Yangtze | 64-2 | Pengze, Jiangxi | 29.87 | 116.53 |
| Yangtze | 64-3 | Pengze, Jiangxi | 29.87 | 116.53 |
| Yangtze | 65-2 | Jingdezhen, Jiangxi | 29.57 | 117.58 |
| Yangtze | 65-4 | Jingdezhen, Jiangxi | 29.57 | 117.58 |
| Yangtze | 66-5 | Jingdezhen, Jiangxi | 29.34 | 117.34 |
| Yangtze | 66-7 | Jingdezhen, Jiangxi | 29.34 | 117.34 |
| Yangtze | 68-3 | Jingdezhen, Jiangxi | 29.32 | 117.31 |
| Yangtze | 68-5 | Jingdezhen, Jiangxi | 29.32 | 117.31 |
| Yangtze | 6-8 | Wuhu, Anhui | 31.39 | 118.38 |
| Yangtze | 69-1 | Wuning, Jiangxi | 28.97 | 114.85 |
| Yangtze | 69-5 | Wuning, Jiangxi | 28.97 | 114.85 |
| Yangtze | 70-2 | Wuning, Jiangxi | 28.99 | 114.86 |
| Yangtze | 70-5 | Wuning, Jiangxi | 28.99 | 114.86 |
| Yangtze | 71-1 | Wuning, Jiangxi | 29 | 114.87 |
| Yangtze | 71-4 | Wuning, Jiangxi | 29 | 114.87 |
| Yangtze | 7-15 | Xiuning, Anhui | 29.8 | 118.18 |
| Yangtze | 7-1 | Xiuning, Anhui | 29.8 | 118.18 |
| Yangtze | 72-4 | Wuning, Jiangxi | 29.24 | 115.1 |
| Yangtze | 72-6 | Wuning, Jiangxi | 29.24 | 115.1 |
| Yangtze | 73-9 | Xingzi, Jiangxi | 29.29 | 115.98 |

| | | | | |
|---|---|---|---|---|
| Yangtze | 74-10 | Jiujiang, Jiangxi | 29.61 | 115.93 |
| Yangtze | 74-8 | Jiujiang, Jiangxi | 29.61 | 115.93 |
| Yangtze | 75-3 | Jiujiang, Jiangxi | 29.6 | 115.93 |
| Yangtze | 75-5 | Jiujiang, Jiangxi | 29.6 | 115.93 |
| Yangtze | 77-5 | Ningxiang, Hunan | 28.23 | 112.55 |
| Yangtze | 78-1 | Taojiang, Hunan | 28.5 | 112.14 |
| Yangtze | 79-6 | Chenxi, Hunan | 28.03 | 110.21 |
| Yangtze | 80-1 | Tongbai, Henan | 32.38 | 113.39 |
| Yangtze | 80-2 | Tongbai, Henan | 32.38 | 113.39 |
| Yangtze | 81-10 | Tongbai, Henan | 32.38 | 113.39 |
| Yangtze | 81-2 | Tongbai, Henan | 32.38 | 113.39 |
| Yangtze | 81-3 | Tongbai, Henan | 32.38 | 113.39 |
| Yangtze | 82-6 | Shangcheng, Henan | 31.82 | 115.39 |
| Yangtze | 83-3 | Gushi, Henan | 32.17 | 115.66 |
| Yangtze | 84-1 | Huangchuan, Henan | 32.14 | 115.06 |
| Yangtze | 85-1 | Huangchuan, Henan | 32.14 | 115.06 |
| Yangtze | 88-1 | Wuxue, Hubei | 29.98 | 115.62 |
| Yangtze | 88-2 | Wuxue, Hubei | 29.98 | 115.62 |
| Yangtze | 88-3 | Wuxue, Hubei | 29.98 | 115.62 |
| Yangtze | 8-8 | Yixian, Anhui | 29.96 | 117.96 |
| Yangtze | 89-1 | Qichun, Hubei | 30.27 | 115.41 |
| Yangtze | 89-2 | Qichun, Hubei | 30.27 | 115.41 |
| Yangtze | 89-5 | Qichun, Hubei | 30.27 | 115.41 |
| Yangtze | 90-1 | Hanchuan, Hubei | 30.53 | 113.81 |
| Yangtze | 90-2 | Hanchuan, Hubei | 30.53 | 113.81 |
| Yangtze | 90-6 | Hanchuan, Hubei | 30.53 | 113.81 |
| Yangtze | 9-10 | Yuexi, Anhui | 30.71 | 116.26 |
| Yangtze | 91-1 | Hanchuan, Hubei | 30.53 | 113.81 |
| Yangtze | 9-15 | Yuexi, Anhui | 30.71 | 116.26 |
| Yangtze | 92-2 | Jingmen, Hubei | 31 | 112.12 |
| Yangtze | 92-4 | Jingmen, Hubei | 31 | 112.12 |
| Yangtze | 92-6 | Jingmen, Hubei | 31 | 112.12 |
| Yangtze | 93-2 | Xiangyang, Hubei | 32.04 | 112.09 |
| Yangtze | 93-5 | Xiangyang, Hubei | 32.04 | 112.09 |
| Yangtze | 93-6 | Xiangyang, Hubei | 32.04 | 112.09 |
| Yangtze | 94-1 | Xiangyang, Hubei | 32.01 | 112.1 |
| Yangtze | 94-2 | Xiangyang, Hubei | 32.01 | 112.1 |
| Yangtze | 95-3 | Yixian, Anhui | 30 | 118.03 |

Note: NW. China & C. Asia, northwestern China and central Asia; Yangtze, Yangtze River basin population.

Table S3. Summary of significant peaks in TE family expression level GWAS analysis.

| Lead SNP | TE family | P value | Peak type |
|---|---|---|---|
| chr2:545516 | ARNOLD1 | 7.26E-60 | same family |
| chr2:9835396 | ARNOLD2 | 4.57E-11 | others |
| chr3:13493764 | ARNOLD3 | 3.53E-37 | same family |
| chr3:15869093 | ARNOLDY1 | 3.60E-17 | others |
| chr1:21736735 | ARNOLDY2 | 6.38E-09 | others |
| chr5:26428195 | AT9MU1 | 4.07E-39 | others |
| chr5:26428195 | AT9NMU1 | 1.22E-23 | same family |
| chr4:11390163 | ATCOPIA10 | 4.90E-11 | same family |
| chr3:15102607 | ATCOPIA11 | 2.40E-10 | others |
| chr4:11241351 | ATCOPIA11 | 5.24E-09 | others |
| chr3:11118309 | ATCOPIA14 | 3.14E-19 | others |
| chr5:13350109 | ATCOPIA15 | 3.26E-13 | same family |
| chr5:21852849 | ATCOPIA16 | 6.31E-09 | others |
| chr4:6702682 | ATCOPIA17 | 3.52E-58 | same family |
| chr5:16048359 | ATCOPIA18A | 1.40E-10 | others |
| chr5:2442479 | ATCOPIA18 | 7.35E-30 | same family |
| chr3:2207658 | ATCOPIA18 | 3.45E-09 | others |
| chr3:17161175 | ATCOPIA19 | 1.49E-19 | others |
| chr1:24186784 | ATCOPIA20 | 9.56E-16 | others |
| chr1:27601701 | ATCOPIA21 | 4.94E-11 | others |
| chr2:7386454 | ATCOPIA20 | 5.98E-10 | others |
| chr3:22347547 | ATCOPIA22 | 1.41E-11 | others |
| chr1:3627108 | ATCOPIA23 | 7.64E-17 | others |
| chr5:13684120 | ATCOPIA24 | 3.73E-13 | same family |
| chr3:11117812 | ATCOPIA27 | 1.29E-11 | others |
| chr5:14686374 | ATCOPIA28 | 6.54E-12 | same family |
| chr4:10793777 | ATCOPIA28 | 1.70E-10 | others |
| chr4:10339233 | ATCOPIA29 | 5.65E-10 | others |
| chr5:26121244 | ATCOPIA29 | 1.25E-09 | others |
| chr2:7996680 | ATCOPIA30 | 4.04E-47 | others |
| chr3:22903228 | ATCOPIA31 | 2.53E-22 | others |
| chr4:6382376 | ATCOPIA31 | 1.13E-36 | others |
| chr2:4007630 | ATCOPIA32B | 3.41E-14 | same family |
| chr1:540339 | ATCOPIA32B | 1.25E-10 | others |
| chr2:4114169 | ATCOPIA32 | 2.23E-14 | others |
| chr5:7896154 | ATCOPIA32 | 8.54E-12 | others |
| chr3:15507824 | ATCOPIA34 | 1.25E-17 | same family |
| chr5:18849826 | ATCOPIA35 | 1.36E-18 | others |
| chr1:12817581 | ATCOPIA36 | 1.98E-15 | same family |
| chr1:13055327 | ATCOPIA3 | 7.14E-10 | others |
| chr5:15809686 | ATCOPIA3 | 1.16E-10 | others |
| chr1:13210172 | ATCOPIA43 | 3.99E-36 | others |

| | | | |
|---|---|---|---|
| chr2:13194438 | ATCOPIA44 | 3.35E-09 | others |
| chr4:14244202 | ATCOPIA46 | 1.45E-19 | same family |
| chr4:17321503 | ATCOPIA46 | 4.61E-12 | others |
| chr3:2239972 | ATCOPIA48 | 3.70E-12 | others |
| chr3:10494234 | ATCOPIA50 | 1.57E-09 | same family |
| chr4:11390942 | ATCOPIA53 | 4.30E-26 | others |
| chr1:12817581 | ATCOPIA57 | 3.01E-14 | same family |
| chr1:19484366 | ATCOPIA59 | 5.36E-19 | others |
| chr5:19235647 | ATCOPIA61 | 1.20E-17 | same family |
| chr3:15775672 | ATCOPIA62 | 7.66E-15 | same family |
| chr3:6774161 | ATCOPIA63 | 2.26E-10 | others |
| chr4:7839065 | ATCOPIA63 | 8.41E-42 | others |
| chr2:18205978 | ATCOPIA65A | 1.01E-10 | others |
| chr2:11414196 | ATCOPIA65 | 1.54E-10 | others |
| chr3:15866287 | ATCOPIA69 | 1.42E-15 | same family |
| chr2:7230176 | ATCOPIA70 | 6.17E-11 | same family |
| chr1:10120008 | ATCOPIA71 | 4.43E-10 | others |
| chr2:7454593 | ATCOPIA71 | 2.71E-12 | same family |
| chr1:16273576 | ATCOPIA72 | 1.37E-11 | others |
| chr2:1683943 | ATCOPIA72 | 1.61E-18 | others |
| chr3:10494234 | ATCOPIA74 | 7.29E-22 | same family |
| chr4:2656886 | ATCOPIA76 | 8.97E-24 | others |
| chr2:6023757 | ATCOPIA77 | 4.04E-18 | same family |
| chr1:19586307 | ATCOPIA81 | 2.37E-26 | others |
| chr5:20204392 | ATCOPIA82 | 1.22E-19 | same family |
| chr5:17958733 | ATCOPIA82 | 2.53E-14 | others |
| chr2:1454491 | ATCOPIA87 | 4.40E-11 | others |
| chr5:11408000 | ATCOPIA88 | 1.20E-09 | others |
| chr3:3937527 | ATCOPIA89 | 3.20E-14 | others |
| chr1:18022839 | ATCOPIA91 | 9.89E-11 | others |
| chr5:14142812 | ATCOPIA91 | 2.00E-13 | same family |
| chr5:10298842 | ATCOPIA92 | 1.98E-12 | same family |
| chr4:737265 | ATCOPIA92 | 3.67E-12 | others |
| chr5:15189300 | ATDNA12T3_2 | 5.02E-42 | same family |
| chr5:18105117 | ATDNA2T9B | 6.29E-13 | others |
| chr2:17291660 | ATDNAI27T9C | 3.41E-45 | same family |
| chr1:11586171 | ATDNATA1 | 6.50E-10 | others |
| chr5:6928072 | ATENSPM10 | 1.53E-13 | others |
| chr2:9255919 | ATENSPM11 | 1.20E-12 | others |
| chr1:24432078 | ATENSPM1 | 2.75E-11 | others |
| chr5:15254733 | ATENSPM6 | 1.72E-13 | others |
| chr2:9277465 | ATENSPM7 | 7.91E-13 | others |
| chr1:23535159 | ATENSPM7 | 3.17E-16 | others |
| chr1:21222585 | ATENSPM9 | 1.31E-11 | others |
| chr2:10852535 | ATENSPM9 | 1.63E-10 | others |
| chr3:7016283 | ATGP10 | 9.75E-12 | others |
| chr2:2148268 | ATGP1 | 1.79E-11 | others |

| | | | |
|---|---|---|---|
| chr1:8242240 | ATGP2 | 2.26E-12 | others |
| chr1:4148612 | ATGP3 | 2.37E-09 | others |
| chr3:6015124 | ATGP3 | 1.26E-14 | others |
| chr4:7881096 | ATGP3 | 2.15E-11 | others |
| chr2:5800623 | ATHAT10 | 1.12E-09 | others |
| chr1:16317933 | ATHAT1 | 6.84E-10 | others |
| chr1:27652049 | ATHAT3 | 4.36E-09 | others |
| chr4:13977304 | ATHAT3 | 5.50E-09 | others |
| chr2:1584389 | ATHAT7 | 9.62E-18 | others |
| chr2:4943212 | ATHATN1 | 3.35E-16 | others |
| chr5:9263425 | ATHATN1 | 2.69E-14 | others |
| chr5:4967255 | ATHATN4 | 5.16E-11 | others |
| chr3:15677944 | ATHATN6 | 5.93E-30 | same family |
| chr3:5328864 | ATHATN6 | 2.30E-10 | others |
| chr1:16120148 | ATHILA2 | 9.36E-11 | same family |
| chr2:19142628 | ATHILA2 | 2.06E-12 | others |
| chr1:16975207 | ATHILA3 | 4.28E-14 | others |
| chr2:1714434 | ATHILA3 | 1.37E-10 | others |
| chr1:29398093 | ATHILA4C | 8.34E-12 | others |
| chr1:22601456 | ATHILA4C | 5.21E-10 | others |
| chr4:711215 | ATHILA4C | 7.92E-15 | others |
| chr4:6777205 | ATHILA4D_LTR | 5.57E-13 | same family |
| chr1:27706514 | ATHILA6A | 1.50E-11 | others |
| chr2:2151176 | ATHILA6A | 2.65E-11 | others |
| chr5:4957575 | ATHILA6B | 3.51E-11 | others |
| chr2:767986 | ATHILA8A | 3.86E-09 | others |
| chr4:15344984 | ATHILA8A | 2.22E-12 | others |
| chr3:4453663 | ATHPOGO | 4.06E-09 | others |
| chr1:20175651 | ATHPOGON3 | 3.17E-12 | same family |
| chr2:2380483 | ATIS112A | 1.43E-10 | others |
| chr3:14167801 | ATLANTYS1 | 6.95E-28 | same family |
| chr1:6985756 | ATLANTYS2 | 2.31E-14 | others |
| chr3:16840925 | ATLANTYS2 | 2.33E-15 | others |
| chr3:23091510 | ATLANTYS3 | 2.14E-12 | same family |
| chr4:6031362 | ATLINE1_1 | 4.56E-10 | others |
| chr4:7956078 | ATLINE1_5 | 3.64E-12 | others |
| chr3:15392159 | ATLINE1_6 | 1.68E-09 | same family |
| chr5:10476340 | ATLINE1A | 2.21E-11 | others |
| chr4:10815748 | ATLINEIII | 3.03E-10 | others |
| chr2:9843129 | ATMU6 | 8.47E-13 | others |
| chr5:10846186 | ATMU6N1 | 4.75E-14 | others |
| chr1:16536079 | ATMU9 | 6.12E-09 | others |
| chr1:13383948 | ATMUN2 | 9.03E-11 | others |
| chr2:14680602 | ATMUN2 | 6.07E-10 | others |
| chr5:17958733 | ATMUNX1 | 5.02E-11 | others |
| chr2:16748195 | ATN9_1 | 2.29E-09 | others |
| chr3:23071659 | ATRE1 | 1.85E-12 | others |

| | | | |
|---|---|---|---|
| chr2:8230956 | ATRE1 | 3.21E-09 | others |
| chr1:19434047 | ATREP10C | 8.64E-09 | others |
| chr1:9402186 | ATREP10D | 7.18E-09 | others |
| chr4:6736283 | ATREP13 | 7.43E-21 | others |
| chr1:22226311 | ATREP14 | 4.82E-10 | others |
| chr2:11050952 | ATREP16 | 8.23E-09 | others |
| chr1:26513099 | ATREP2 | 4.71E-10 | others |
| chr1:13210878 | ATREP3 | 2.02E-09 | same family |
| chr1:24997099 | ATREP4 | 3.95E-09 | others |
| chr1:11311394 | ATREP5 | 1.77E-11 | same family |
| chr4:7990444 | ATREP5 | 5.44E-11 | same family |
| chr1:17853243 | ATREP6 | 7.53E-10 | others |
| chr4:7842365 | ATREP9 | 3.95E-09 | others |
| chr2:19062038 | ATSINE2A | 6.32E-10 | others |
| chr5:17403448 | ATSINE2A | 7.74E-17 | others |
| chr3:18730206 | ATSINE4 | 1.04E-13 | same family |
| chr4:11586610 | ATSINE4 | 1.48E-10 | others |
| chr4:10809686 | ATSINE4 | 6.04E-14 | others |
| chr1:21733864 | ATTIR16T3A | 1.54E-13 | others |
| chr5:22750179 | ATTIR16T3A | 6.11E-12 | others |
| chr3:16128233 | ATTIRX1C | 7.11E-11 | others |
| chr2:1730606 | BOMZH2 | 5.46E-09 | others |
| chr1:13950766 | BRODYAGA1A | 9.65E-11 | same family |
| chr2:13189113 | BRODYAGA1A | 9.11E-14 | others |
| chr5:20202759 | BRODYAGA1 | 4.37E-09 | others |
| chr4:13391455 | DT1 | 5.57E-12 | others |
| chr3:10761315 | DT1 | 1.91E-09 | others |
| chr1:13255134 | ENDOVIR1 | 8.72E-19 | others |
| chr2:2303637 | ENDOVIR1 | 8.85E-17 | others |
| chr1:13352257 | HELITRON1 | 3.42E-12 | same family |
| chr3:17983921 | HELITRON1 | 1.35E-30 | others |
| chr2:12255099 | HELITRON2 | 9.41E-14 | others |
| chr4:11341911 | HELITRONY1B | 4.14E-09 | others |
| chr2:7995404 | HELITRONY1C | 9.79E-17 | same family |
| chr4:972969 | HELITRONY1C | 6.43E-13 | others |
| chr5:22246184 | HELITRONY1E | 6.62E-37 | same family |
| chr2:18276687 | HELITRONY1E | 1.43E-10 | same family |
| chr4:5530600 | HELITRONY3 | 2.05E-10 | others |
| chr2:305938 | HELITRONY1D | 4.09E-09 | others |
| chr3:17117971 | LIMPET1 | 4.99E-12 | same family |
| chr3:7794973 | RathE2_cons | 5.43E-27 | others |
| chr5:10236863 | ROMANIAT5 | 6.44E-17 | others |
| chr5:15191802 | RP1_AT | 2.27E-45 | same family |
| chr3:21416229 | SADHU | 4.59E-11 | others |
| chr5:5818285 | SIMPLEHAT2 | 3.34E-09 | others |
| chr3:10978600 | TA11 | 1.13E-10 | same family |
| chr3:22014347 | TA12 | 3.63E-10 | others |

| | | | |
|---|---|---|---|
| chr3:3080703 | TA12 | 6.21E-11 | others |
| chr4:6409588 | TAG1 | 2.08E-21 | others |
| chr3:15507824 | TNAT1A | 1.48E-15 | same family |
| chr5:6991437 | TNAT1A | 1.52E-09 | others |
| chr3:16601301 | TNAT2A | 3.80E-38 | others |
| chr1:25354273 | TSCL | 2.84E-15 | others |
| chr1:27638039 | TSCL | 8.27E-14 | others |
| chr4:11307964 | TSCL | 7.47E-12 | others |
| chr1:2480692 | VANDAL10 | 5.63E-11 | others |
| chr2:1557162 | VANDAL10 | 3.38E-11 | others |
| chr5:25473574 | VANDAL10 | 3.90E-10 | others |
| chr4:2139727 | VANDAL11 | 8.67E-10 | others |
| chr2:7298605 | VANDAL14 | 8.32E-11 | others |
| chr3:15887401 | VANDAL16 | 2.18E-13 | same family |
| chr5:7670238 | VANDAL18NB | 8.98E-16 | others |
| chr5:15486428 | VANDAL18NB | 2.34E-16 | others |
| chr3:22047798 | VANDAL22 | 1.05E-09 | others |
| chr2:10736036 | VANDAL2N1 | 2.51E-11 | same family |
| chr1:29398093 | VANDAL4 | 3.73E-12 | others |
| chr4:11398157 | VANDAL4 | 2.05E-10 | others |
| chr4:1559762 | VANDAL5 | 9.44E-17 | others |
| chr4:2718352 | VANDAL5 | 8.10E-51 | same family |
| chr1:11453796 | VANDAL6 | 9.06E-14 | others |
| chr3:8003382 | VANDAL7 | 7.01E-10 | others |
| chr3:23156856 | VANDAL7 | 1.07E-09 | others |
| chr1:23588573 | VANDALNX2 | 4.47E-32 | same family |

Note: in peak type, same family represents a peak contain TEs of same family, others represents a peak not contain TEs of the same family.

Table S4. Summary of candidate causal TEs associated with TE family expression level variation.

| Lead SNP | TE family | Candidate TE | TE type |
|---|---|---|---|
| chr2:545516 | ARNOLD1 | AT2TE02465 | polymorphic |
| chr3:13493764 | ARNOLD3 | chr3:13534057:13534400:ARNOLD3-DNA/MuDR | polymorphic |
| chr3:13493764 | ARNOLD3 | AT3TE54815 | polymorphic |
| chr5:26428195 | AT9NMU1 | AT5TE94370 | polymorphic |
| chr5:26428195 | AT9NMU1 | AT5TE94375 | polymorphic |
| chr5:26428195 | AT9NMU1 | AT5TE94380 | polymorphic |
| chr4:11390163 | ATCOPIA10 | AT4TE52315 | fixed |
| chr5:13350109 | ATCOPIA15 | AT5TE47100 | polymorphic |
| chr4:6702682 | ATCOPIA17 | AT4TE28870 | polymorphic |
| chr5:2442479 | ATCOPIA18 | AT5TE08870 | polymorphic |
| chr5:13684120 | ATCOPIA24 | AT5TE48930 | polymorphic |
| chr5:14686374 | ATCOPIA28 | AT5TE52880 | fixed |
| chr2:4007630 | ATCOPIA32B | AT2TE17870 | fixed |
| chr3:15507824 | ATCOPIA34 | chr3:15506098:15506159:ATCOPIA34-LTR/Copia | polymorphic |
| chr1:12817581 | ATCOPIA36 | AT1TE41800 | fixed |
| chr4:14244202 | ATCOPIA46 | AT4TE67490 | polymorphic |
| chr3:10494234 | ATCOPIA50 | AT3TE43605 | fixed |
| chr3:10494234 | ATCOPIA50 | AT3TE43635 | fixed |
| chr1:12817581 | ATCOPIA57 | chr1:12841985:12842079:ATCOPIA57-LTR/Copia | polymorphic |
| chr1:12817581 | ATCOPIA57 | chr1:12959513:12959601:ATCOPIA57-LTR/Copia | polymorphic |
| chr1:12817581 | ATCOPIA57 | AT1TE41795 | fixed |
| chr5:19235647 | ATCOPIA61 | AT5TE69245 | polymorphic |
| chr3:15775672 | ATCOPIA62 | AT3TE63760 | polymorphic |
| chr3:15775672 | ATCOPIA62 | AT3TE63765 | polymorphic |
| chr3:15866287 | ATCOPIA69 | chr3:15839119:15839204:ATCOPIA69-LTR/Copia | polymorphic |
| chr3:15866287 | ATCOPIA69 | chr3:15839470:15839525:ATCOPIA69-LTR/Copia | polymorphic |
| chr2:7230176 | ATCOPIA70 | AT2TE29450 | polymorphic |
| chr2:7454593 | ATCOPIA71 | AT2TE30555 | polymorphic |
| chr2:7454593 | ATCOPIA71 | AT2TE30560 | polymorphic |
| chr3:10494234 | ATCOPIA74 | AT3TE43610 | fixed |
| chr2:6023757 | ATCOPIA77 | AT2TE24515 | fixed |
| chr2:6023757 | ATCOPIA77 | AT2TE24520 | fixed |
| chr2:6023757 | ATCOPIA77 | AT2TE24530 | fixed |
| chr5:20204392 | ATCOPIA82 | chr5:20205280:20205410:ATCOPIA82-LTR/Copia | polymorphic |
| chr5:14142812 | ATCOPIA91 | AT5TE50380 | polymorphic |
| chr5:10298842 | ATCOPIA92 | AT5TE37810 | fixed |
| chr5:10298842 | ATCOPIA92 | AT5TE37815 | fixed |
| chr5:10298842 | ATCOPIA92 | AT5TE37820 | fixed |
| chr5:15189300 | ATDNA12T3_2 | AT5TE55005 | fixed |

| | | | |
|---|---|---|---|
| chr2:17291660 | ATDNAI27T9C | AT2TE78945 | fixed |
| chr3:15677944 | ATHATN6 | AT3TE63400 | polymorphic |
| chr1:16120148 | ATHILA2 | chr1:16517493:16517580:ATHILA2-LTR/Gypsy | polymorphic |
| chr1:16120148 | ATHILA2 | AT1TE53410 | polymorphic |
| chr4:6777205 | ATHILA4D_LTR | AT4TE29165 | fixed |
| chr1:20175651 | ATHPOGON3 | chr1:20172844:20172917:ATHPOGON3-DNA/Pogo | polymorphic |
| chr1:20175651 | ATHPOGON3 | AT1TE66580 | fixed |
| chr3:14167801 | ATLANTYS1 | chr3:14083570:14083718:ATLANTYS1-LTR/Gypsy | polymorphic |
| chr3:14167801 | ATLANTYS1 | AT3TE57970 | fixed |
| chr3:14167801 | ATLANTYS1 | AT3TE58560 | fixed |
| chr3:14167801 | ATLANTYS1 | AT3TE58580 | fixed |
| chr3:14167801 | ATLANTYS1 | AT3TE57885 | fixed |
| chr3:23091510 | ATLANTYS3 | chr3:23135754:23135822:ATLANTYS3-LTR/Gypsy | polymorphic |
| chr3:15392159 | ATLINE1_6 | AT3TE62895 | fixed |
| chr3:15392159 | ATLINE1_6 | AT3TE62785 | polymorphic |
| chr1:13210878 | ATREP3 | chr1:13217358:13217426:ATREP3-RC/Helitron | polymorphic |
| chr1:13210878 | ATREP3 | AT1TE43145 | fixed |
| chr1:11311394 | ATREP5 | AT1TE36580 | fixed |
| chr4:7990444 | ATREP5 | AT4TE35190 | fixed |
| chr3:18730206 | ATSINE4 | chr3:18739736:18739786:ATSINE4-SINE | polymorphic |
| chr3:18730206 | ATSINE4 | AT3TE76015 | fixed |
| chr1:13950766 | BRODYAGA1A | chr1:13904892:13904966:BRODYAGA1A-DNA/MuDR | polymorphic |
| chr1:13950766 | BRODYAGA1A | AT1TE45730 | polymorphic |
| chr1:13950766 | BRODYAGA1A | AT1TE45760 | fixed |
| chr1:13950766 | BRODYAGA1A | AT1TE45910 | fixed |
| chr1:13950766 | BRODYAGA1A | AT1TE46055 | fixed |
| chr1:13950766 | BRODYAGA1A | AT1TE46060 | fixed |
| chr1:13352257 | HELITRON1 | AT1TE43740 | polymorphic |
| chr2:7995404 | HELITRONY1C | AT2TE33290 | fixed |
| chr2:18276687 | HELITRONY1E | AT2TE82920 | fixed |
| chr5:22246184 | HELITRONY1E | AT5TE80455 | fixed |
| chr3:17117971 | LIMPET1 | chr3:17116464:17116519:LIMPET1-DNA/MuDR | polymorphic |
| chr5:15191802 | RP1_AT | AT5TE55010 | fixed |
| chr3:10978600 | TA11 | AT3TE45620 | polymorphic |
| chr3:15507824 | TNAT1A | AT3TE62385 | fixed |
| chr3:15887401 | VANDAL16 | chr3:15952104:15952237:VANDAL16-DNA/MuDR | polymorphic |
| chr2:10736036 | VANDAL2N1 | chr2:10736424:10736533:VANDAL2N1-DNA/MuDR | polymorphic |
| chr4:2718352 | VANDAL5 | AT4TE13020 | fixed |
| chr4:2718352 | VANDAL5 | AT4TE13025 | fixed |
| chr4:2718352 | VANDAL5 | AT4TE13045 | polymorphic |
| chr4:2718352 | VANDAL5 | AT4TE13030 | fixed |
| chr1:23588573 | VANDALNX2 | AT1TE77670 | fixed |

Note: TE id in TAIR10 (such as AT3TE63765) was used if a TE is present in the reference genome, TE position (such as chr2:10736424:10736533: VANDAL2N1-DNA/MuDR, which including the chromosome, start position, end position and TE family) was used if a TE is absent in the reference genome.

Table S5. The correlation coefficients of candidate causal TE expression levels and their family expression levels.

| **Lead SNP** | **TE family** | **Causal TE** | **R** |
|---|---|---|---|
| chr2:545516 | ARNOLD1 | AT2TE02465 | 0.098435694 |
| chr3:13493764 | ARNOLD3 | AT3TE54815 | 0.742142287 |
| chr5:26428195 | AT9NMU1 | AT5TE94370 | 0.037169895 |
| chr5:26428195 | AT9NMU1 | AT5TE94375 | 0.024613388 |
| chr5:26428195 | AT9NMU1 | AT5TE94380 | NA |
| chr4:11390163 | ATCOPIA10 | AT4TE52315 | 0.830672823 |
| chr5:13350109 | ATCOPIA15 | AT5TE47100 | 0.967443457 |
| chr4:6702682 | ATCOPIA17 | AT4TE28870 | 0.956734141 |
| chr5:2442479 | ATCOPIA18 | AT5TE08870 | 0.999577777 |
| chr5:13684120 | ATCOPIA24 | AT5TE48930 | 0.998150705 |
| chr5:14686374 | ATCOPIA28 | AT5TE52880 | 0.162248194 |
| chr2:4007630 | ATCOPIA32B | AT2TE17870 | 0.990345418 |
| chr1:12817581 | ATCOPIA36 | AT1TE41800 | 0.994997409 |
| chr4:14244202 | ATCOPIA46 | AT4TE67490 | 0.990261062 |
| chr3:10494234 | ATCOPIA50 | AT3TE43605 | 0.734486501 |
| chr3:10494234 | ATCOPIA50 | AT3TE43635 | 0.674897728 |
| chr1:12817581 | ATCOPIA57 | AT1TE41795 | 0.982444703 |
| chr5:19235647 | ATCOPIA61 | AT5TE69245 | 0.969497496 |
| chr3:15775672 | ATCOPIA62 | AT3TE63760 | NA |
| chr3:15775672 | ATCOPIA62 | AT3TE63765 | 0.945872814 |
| chr2:7230176 | ATCOPIA70 | AT2TE29450 | 0.97944912 |
| chr2:7454593 | ATCOPIA71 | AT2TE30555 | 0.997712453 |
| chr2:7454593 | ATCOPIA71 | AT2TE30560 | NA |
| chr3:10494234 | ATCOPIA74 | AT3TE43610 | 0.993807552 |
| chr2:6023757 | ATCOPIA77 | AT2TE24515 | NA |
| chr2:6023757 | ATCOPIA77 | AT2TE24520 | 0.996930073 |
| chr2:6023757 | ATCOPIA77 | AT2TE24530 | 0.134334584 |
| chr5:14142812 | ATCOPIA91 | AT5TE50380 | 0.998384176 |
| chr5:10298842 | ATCOPIA92 | AT5TE37810 | NA |
| chr5:10298842 | ATCOPIA92 | AT5TE37815 | NA |
| chr5:10298842 | ATCOPIA92 | AT5TE37820 | 0.926121972 |
| chr5:15189300 | ATDNA12T3_2 | AT5TE55005 | 0.945574209 |
| chr2:17291660 | ATDNAI27T9C | AT2TE78945 | 0.045733922 |
| chr3:15677944 | ATHATN6 | AT3TE63400 | 0.842708471 |
| chr1:16120148 | ATHILA2 | AT1TE53410 | NA |
| chr4:6777205 | ATHILA4D_LTR | AT4TE29165 | 0.250992082 |
| chr1:20175651 | ATHPOGON3 | AT1TE66580 | NA |
| chr3:14167801 | ATLANTYS1 | AT3TE57970 | NA |
| chr3:14167801 | ATLANTYS1 | AT3TE58560 | 0.987130681 |
| chr3:14167801 | ATLANTYS1 | AT3TE58580 | 0.238307044 |
| chr3:14167801 | ATLANTYS1 | AT3TE57885 | 0.186826545 |
| chr3:15392159 | ATLINE1_6 | AT3TE62895 | 0.070348676 |
| chr3:15392159 | ATLINE1_6 | AT3TE62785 | 0.628678387 |

| | | | |
|---|---|---|---|
| chr1:13210878 | ATREP3 | AT1TE43145 | NA |
| chr1:11311394 | ATREP5 | AT1TE36580 | 0.04013176 |
| chr4:7990444 | ATREP5 | AT4TE35190 | NA |
| chr3:18730206 | ATSINE4 | AT3TE76015 | 0.928533251 |
| chr1:13950766 | BRODYAGA1A | AT1TE45730 | -0.00834212 |
| chr1:13950766 | BRODYAGA1A | AT1TE45760 | NA |
| chr1:13950766 | BRODYAGA1A | AT1TE45910 | 0.05194346 |
| chr1:13950766 | BRODYAGA1A | AT1TE46055 | NA |
| chr1:13950766 | BRODYAGA1A | AT1TE46060 | NA |
| chr1:13352257 | HELITRON1 | AT1TE43740 | 0.188352186 |
| chr2:7995404 | HELITRONY1C | AT2TE33290 | 0.070850025 |
| chr2:18276687 | HELITRONY1E | AT2TE82920 | NA |
| chr5:22246184 | HELITRONY1E | AT5TE80455 | -7.52E-05 |
| chr5:15191802 | RP1_AT | AT5TE55010 | 0.998960632 |
| chr3:10978600 | TA11 | AT3TE45620 | 0.628085453 |
| chr3:15507824 | TNAT1A | AT3TE62385 | 0.129499683 |
| chr4:2718352 | VANDAL5 | AT4TE13020 | NA |
| chr4:2718352 | VANDAL5 | AT4TE13025 | NA |
| chr4:2718352 | VANDAL5 | AT4TE13045 | 0.048403416 |
| chr4:2718352 | VANDAL5 | AT4TE13030 | 0.002305425 |
| chr1:23588573 | VANDALNX2 | AT1TE77670 | 0.879999491 |

Note: NA occurs when the given TE is not expressed in any accession. 17 non-reference TEs were not shown as we could not calculate the expression level of these TEs.

Table S6. Summary of candidate genes associated with TE family expression level variation.

| Lead SNP | TE family | Gene |
|---|---|---|
| chr2:1683943 | ATCOPIA72-LTR/Copia | AT2G04780,AT2G04790,AT2G04795,AT2G04810,AT2G04830,AT2G04840,AT2G04842,AT2G04850,AT2G04860,AT2G04800,AT2G04845 |
| chr1:13210172 | ATCOPIA43-LTR/Copia | AT1G35660,AT1G35680,AT1G35720,AT1G35730,AT1G35670,AT1G35710,AT1G35750 |
| chr1:13950766 | BRODYAGA1A-DNA/MuDR | AT1G36730,AT1G36745,AT1G36756,AT1G36920,AT1G36922,AT1G36925,AT1G36940,AT1G36942,AT1G36950,AT1G36960,AT1G36970,AT1G36990,AT1G37000,AT1G37020,AT1G37010,AT1G36980 |
| chr4:711215 | ATHILA4C-LTR/Gypsy | AT4G01600,AT4G01610,AT4G01630,AT4G01640,AT4G01660,AT4G01670,AT4G01671,AT4G01680,AT4G01690,AT4G01700,AT4G01703,AT4G01650 |
| chr1:4148612 | ATGP3-LTR/Gypsy | AT1G12160,AT1G12180,AT1G12190,AT1G12200,AT1G12210,AT1G12211,AT1G12230,AT1G12240,AT1G12244,AT1G12250,AT1G12260,AT1G12170,AT1G12220 |
| chr3:15866287 | ATCOPIA69-LTR/Copia | AT3G44050,AT3G44060,AT3G44070,AT3G44080,AT3G44090,AT3G44100,AT3G44110,AT3G44115,AT3G44120 |
| chr3:23091510 | ATLANTYS3-LTR/Gypsy | AT3G62530,AT3G62550,AT3G62560,AT3G62570,AT3G62580,AT3G62600,AT3G62610,AT3G62615,AT3G62620,AT3G62630,AT3G62540,AT3G62590 |
| chr3:5328864 | ATHATN6-DNA/HAT | AT3G15650,AT3G15670,AT3G15680,AT3G15690,AT3G15700,AT3G15720,AT3G15730,AT3G15740,AT3G15750,AT3G15760,AT3G15780,AT3G15790,AT3G15800,AT3G15810,AT3G15820,AT3G15830,AT3G15850,AT3G15860,AT3G15660,AT3G15710,AT3G15770,AT3G15840 |
| chr3:13493764 | ARNOLD3-DNA/MuDR | AT3G32896,AT3G32904,AT3G32920,AT3G32930,AT3G32940,AT3G32960,AT3G32980,AT3G33187,AT3G33293,AT3G33393,AT3G33494,AT3G33528,AT3G33520(ARP6),AT3G41762,AT3G42050,AT3G42075,AT3G42130,AT3G42140,AT3G42148,AT3G42150,AT3G42153,AT3G42155,AT3G42160,AT3G42170,AT3G42180,AT3G42310,AT3G42390,AT3G42473,AT3G42550,AT3G42560,AT3G42565,AT3G42570,AT3G42628,AT3G42630,AT3G33530,AT3G42060 |

| | | |
|---|---|---|
| chr2:6023757 | ATCOPIA77-LTR/Copia | AT2G14210,AT2G14247,AT2G14255,AT2G14260,AT2G14265,AT2G14282,AT2G14270 |
| chr2:2380483 | ATIS112A-Harbinger | AT2G06090,AT2G06095,AT2G06105,AT2G06166,AT2G06200,AT2G06210 |
| chr2:18276687 | HELITRONY1E-RC/Helitron | AT2G44150,AT2G44160,AT2G44175,AT2G44180,AT2G44190,AT2G44195,AT2G44198,AT2G44200,AT2G44210,AT2G44230,AT2G44240,AT2G44220 |
| chr2:1584389 | ATHAT7-DNA/HAT | AT2G04495,AT2G04500,AT2G04515,AT2G04520,AT2G04530,AT2G04540,AT2G04560,AT2G04570,AT2G04550 |
| chr5:5818285 | SIMPLEHAT2-DNA/HAT | AT5G17610,AT5G17620,AT5G17630,AT5G17640,AT5G17650,AT5G17660,AT5G17670,AT5G17680,AT5G17690,AT5G17700 |
| chr5:15486428 | VANDAL18NB-DNA/MuDR | AT5G38620,AT5G38640,AT5G38660,AT5G38650,AT5G38670,AT5G38680,AT5G38700,AT5G38710,AT5G38720,AT5G38730,AT5G38740,AT5G38743,AT5G38747,AT5G38750,AT5G38760,AT5G38770,AT5G38780,AT5G38790,AT5G38800,AT5G38810,AT5G38820,AT5G38830,AT5G38840,AT5G38850,AT5G38860,AT5G38865,AT5G38890,AT5G38895,AT5G38900,AT5G38910,AT5G38920,AT5G38930,AT5G38940,AT5G38950,AT5G38960,AT5G38980,AT5G38990,AT5G39010,AT5G39020,AT5G39024,AT5G39030,AT5G39040,AT5G39050,AT5G39080,AT5G39090,AT5G38690,AT5G38880,AT5G39000,AT5G38630,AT5G38970 |
| chr5:10846186 | ATMU6N1-DNA/MuDR | AT5G28810,AT5G28820,AT5G28823,AT5G28830,AT5G28780,AT5G28840 |
| chr1:24186784 | ATCOPIA20-LTR/Copia | AT1G65020,AT1G65030,AT1G65032,AT1G65040,AT1G65050,AT1G65060,AT1G65070,AT1G65080,AT1G65110,AT1G65113,AT1G65120,AT1G65140,AT1G65150,AT1G65160,AT1G65165,AT1G65170,AT1G65180,AT1G65190,AT1G65200,AT1G65210,AT1G65220,AT1G65230,AT1G65250,AT1G65260,AT1G65270,AT1G65290,AT1G65295,AT1G65300,AT1G65310,AT1G65320,AT1G65330,AT1G65340,AT1G65342,AT1G65350,AT1G65346,AT1G65349,AT1G65352,AT1G65360,AT1G65370,AT1G65380,AT1G65390,AT1G65420,AT1G65425,AT1G65430,AT1G65450,AT1G65470(FAS1),AT1G65090,AT1G65130,AT1G65240,AT1G65410,AT1G65440,AT1G65280 |
| chr3:2239972 | ATCOPIA48-LTR/Copia | AT3G06850,AT3G06868,AT3G06870,AT3G06880,AT3G06890,AT3G06895,AT3G06920,AT3G06930,AT3G06950,AT3G06970,AT3G06980,AT3G06985,AT3G06990,AT3G07000,AT3G07005,AT3G07010,AT3G07020,AT3G07030,AT3G07050,AT3G07060,AT3G07070,A |

| | | |
|---|---|---|
| | | T3G07080,AT3G07100,AT3G07110,AT3G07120,AT3G07130,AT3 G06860,AT3G06910,AT3G06960,AT3G07090,AT3G07040 |
| chr1:2173386 4 | ATTIR16T3A-DNA | AT1G58450,AT1G58460,AT1G58470,AT1G58525,AT1G58520,AT1 G58602 |
| chr5:2275017 9 | ATTIR16T3A-DNA | AT5G56160,AT5G56170,AT5G56190,AT5G56200,AT5G56210,AT5 G56230,AT5G56240,AT5G56180,AT5G56220 |
| chr1:1802283 9 | ATCOPIA91-LTR/Copia | AT1G48660,AT1G48670,AT1G48690,AT1G48700,AT1G48720,AT1 G48725,AT1G48730,AT1G48740,AT1G48745,AT1G48750,AT1G4 8770,AT1G48780,AT1G48790,AT1G48760 |
| chr4:1139094 2 | ATCOPIA53-LTR/Copia | AT4G21366,AT4G21380,AT4G21390,AT4G21400,AT4G21410 |
| chr5:1810511 7 | ATDNA2T9B-DNA/MuDR | AT5G44770,AT5G44780,AT5G44790,AT5G44820,AT5G44830,AT5 G44840,AT5G44850,AT5G44860,AT5G44870,AT5G44785,AT5G4 4800 |
| chr1:2763803 9 | TSCL-LINE? | AT1G73220,AT1G73240,AT1G73250,AT1G73260,AT1G73270,AT1 G73280,AT1G73300,AT1G73310,AT1G73320,AT1G73325,AT1G7 3330,AT1G73350,AT1G73360,AT1G73370,AT1G73390,AT1G7340 0,AT1G73410,AT1G73440,AT1G73450,AT1G73470,AT1G73480,A T1G73490,AT1G73510,AT1G73530,AT1G73540,AT1G73550,AT1 G73560,AT1G73570,AT1G73580,AT1G73590,AT1G73230,AT1G7 3290,AT1G73340,AT1G73380,AT1G73430,AT1G73460,AT1G7350 0 |
| chr2:1454491 | ATCOPIA87-LTR/Copia | AT2G04220,AT2G04230,AT2G04235,AT2G04240 |
| chr2:2303637 | ENDOVIR1-LTR/Copia | AT2G05900,AT2G05910,AT2G05915,AT2G05920,AT2G05940,AT2 G05970,AT2G05990,AT2G06000,AT2G06005,AT2G06010,AT2G0 5850 |
| chr4:6702682 | ATCOPIA17-LTR/Copia | AT4G10610,AT4G10613,AT4G10620,AT4G10630,AT4G10640,AT4 G10650,AT4G10660,AT4G10670,AT4G10680,AT4G10695,AT4G1 0700,AT4G10720,AT4G10730,AT4G10740,AT4G10760,AT4G1076 7,AT4G10770,AT4G10790,AT4G10800,AT4G10810,AT4G10820,A T4G10840,AT4G10843,AT4G10850,AT4G10860,AT4G10870,AT4 G10880,AT4G10890,AT4G10895,AT4G10910,AT4G10920,AT4G1 0925,AT4G10930,AT4G10950,AT4G10955,AT4G10970,AT4G1100 0,AT4G11020,AT4G11030,AT4G11040,AT4G11060,AT4G11070,A T4G11080,AT4G11090,AT4G11100,AT4G11110,AT4G11120,AT4 G11140,AT4G11150,AT4G11160,AT4G11175,AT4G11180,AT4G1 1190,AT4G11210,AT4G11211,AT4G11220,AT4G11230,AT4G1125 |

| | | |
|---|---|---|
| | | 0,AT4G11260,AT4G11270,AT4G11290,AT4G11300,AT4G11310,AT4G11320,AT4G11330,AT4G11350,AT4G11360,AT4G11370,AT4G11380,AT4G11385,AT4G11393,AT4G11400,AT4G11402,AT4G11420,AT4G11430,AT4G11440,AT4G11450,AT4G11460,AT4G11470,AT4G11480,AT4G11485,AT4G11510,AT4G11521,AT4G11530,AT4G11540,AT4G11543,AT4G11547,AT4G11550,AT4G11560,AT4G11570,AT4G11580,AT4G11590,AT4G11600,AT4G11610,AT4G11630,AT4G11640,AT4G11650,AT4G11653,AT4G11655,AT4G11660,AT4G11680,AT4G11690,AT4G11700,AT4G11720,AT4G11730,AT4G11740,AT4G10710(SPT16),AT4G10750,AT4G10780,AT4G10960,AT4G11010,AT4G11050,AT4G11130(RDR2),AT4G11240,AT4G11280,AT4G11340,AT4G11373,AT4G11390,AT4G11490,AT4G11670,AT4G11170,AT4G11410 |
| chr2:1714434 | ATHILA3-LTR/Gypsy | AT2G04830,AT2G04840,AT2G04842,AT2G04850,AT2G04860,AT2G04865,AT2G04870,AT2G04890,AT2G04900,AT2G04910,AT2G04920,AT2G04880,AT2G04845 |
| chr1:11311394 | ATREP5-RC/Helitron | AT1G31550,AT1G31555,AT1G31580,AT1G31600,AT1G31620,AT1G31630,AT1G31640,AT1G31650 |
| chr1:13352257 | HELITRON1-RC/Helitron | AT1G35880,AT1G35890,AT1G35910,AT1G35895 |
| chr5:6991437 | TNAT1A-DNA | AT5G20640,AT5G20650,AT5G20660,AT5G20670,AT5G20680,AT5G20690,AT5G20700,AT5G20635 |
| chr5:17958733 | ATCOPIA82-LTR/Copia | AT5G44510,AT5G44520,AT5G44540,AT5G44550,AT5G44560,AT5G44563,AT5G44565,AT5G44567,AT5G44568,AT5G44566,AT5G44570,AT5G44572,AT5G44574,AT5G44575,AT5G44578,AT5G44580,AT5G44582,AT5G44585,AT5G44590,AT5G44530 |
| chr5:17958733 | ATMUNX1-DNA/MuDR | AT5G44510,AT5G44520,AT5G44540,AT5G44550,AT5G44560,AT5G44563,AT5G44565,AT5G44567,AT5G44568,AT5G44566,AT5G44570,AT5G44572,AT5G44574,AT5G44575,AT5G44578,AT5G44580,AT5G44582,AT5G44585,AT5G44530 |
| chr5:15254733 | ATENSPM6-DNA/En-Spm | AT5G38195,AT5G38197,AT5G38200,AT5G38210,AT5G38190,AT5G38220 |
| chr3:3080703 | TA12-LINE/L1 | AT3G09980,AT3G09990,AT3G10000,AT3G10010(DML2),AT3G10020,AT3G10030 |
| chr3:22014347 | TA12-LINE/L1 | AT3G59520,AT3G59530,AT3G59540,AT3G59550,AT3G59580,AT3G59590,AT3G59600,AT3G59620,AT3G59630,AT3G59640,AT3G59570,AT3G59610 |

| | | |
|---|---|---|
| chr4:11390163 | ATCOPIA10-LTR/Copia | AT4G21323,AT4G21326,AT4G21330,AT4G21350,AT4G21366,AT4G21380,AT4G21390,AT4G21400,AT4G21430,AT4G21445,AT4G21450,AT4G21460,AT4G21470,AT4G21340,AT4G21440,AT4G21410 |
| chr1:13255134 | ENDOVIR1-LTR/Copia | AT1G35625,AT1G35630,AT1G35660,AT1G35680,AT1G35720,AT1G35730,AT1G35620,AT1G35670,AT1G35710,AT1G35750 |
| chr1:3627108 | ATCOPIA23-LTR/Copia | AT1G10770,AT1G10780,AT1G10790,AT1G10800,AT1G10810,AT1G10830,AT1G10840,AT1G10850,AT1G10865,AT1G10875,AT1G10880,AT1G10890,AT1G10910,AT1G10920,AT1G10940,AT1G10760,AT1G10820,AT1G10870,AT1G10900,AT1G10930 |
| chr2:8230956 | ATRE1-LTR/Copia | AT2G18969,AT2G18970,AT2G18980,AT2G18990,AT2G19010,AT2G18960,AT2G19000 |
| chr2:18205978 | ATCOPIA65A-LTR/Copia | AT2G43920,AT2G43930,AT2G43940,AT2G43945,AT2G43960,AT2G43970,AT2G43980,AT2G43990,AT2G44000,AT2G44010,AT2G44030,AT2G44040,AT2G44050,AT2G44060,AT2G44065,AT2G44070,AT2G43910,AT2G43950,AT2G44020 |
| chr5:6928072 | ATENSPM10-DNA/En-Spm | AT5G20110,AT5G20120,AT5G20130,AT5G20150,AT5G20160,AT5G20165,AT5G20170,AT5G20180,AT5G20200,AT5G20220,AT5G20230,AT5G20240,AT5G20250,AT5G20260,AT5G20270,AT5G20290,AT5G20300,AT5G20310,AT5G20330,AT5G20340,AT5G20360,AT5G20370,AT5G20380,AT5G20390,AT5G20400,AT5G20420(CLSY2),AT5G20430,AT5G20440,AT5G20447,AT5G20450,AT5G20460,AT5G20470,AT5G20480,AT5G20500,AT5G20510,AT5G20520,AT5G20540,AT5G20550,AT5G20560,AT5G20570,AT5G20580,AT5G20590,AT5G20600,AT5G20610,AT5G20620,AT5G20630,AT5G20640,AT5G20650,AT5G20660,AT5G20670,AT5G20680,AT5G20690,AT5G20700,AT5G20720,AT5G20730,AT5G20740,AT5G20790,AT5G20820,AT5G20830,AT5G20850,AT5G20870,AT5G20885,AT5G20890,AT5G20900,AT5G20920,AT5G20930,AT5G20935,AT5G20940,AT5G20960,AT5G20970,AT5G20990,AT5G20995,AT5G21010,AT5G21020,AT5G21040,AT5G21050,AT5G21060,AT5G21070,AT5G21090,AT5G21100,AT5G21120,AT5G21125,AT5G21130,AT5G21140,AT5G21160,AT5G21170,AT5G21274,AT5G21326,AT5G21430,AT5G21482,AT5G21900,AT5G21930,AT5G21940,AT5G21950,AT5G21970,AT5G21280,AT5G22000,AT5G22010(RFC1),AT5G22020,AT5G22040,AT5G22050,AT5G22060,AT5G22070,AT5G22080,AT5G22090,AT5G22100,AT5G22110,AT5G22120,AT5G22140,AT5G22145,AT5G22150,AT5G22160,AT5G22170,AT5G22180,AT5G22190,AT5G22210,AT5G22220,AT5G22240,AT5G22250,AT5G2226 |

| | | |
|---|---|---|
| | | 0,AT5G22280,AT5G22290,AT5G22300,AT5G22310,AT5G22330,AT5G22340,AT5G22350,AT5G22355,AT5G22370,AT5G22380,AT5G20140,AT5G20280,AT5G20320(DCL4),AT5G20350,AT5G20410,AT5G20490,AT5G20635,AT5G20710,AT5G20810,AT5G20840,AT5G20910,AT5G20950,AT5G20980,AT5G21030,AT5G21080,AT5G21105,AT5G21150(AGO9),AT5G21910,AT5G21920,AT5G21960,AT5G21990,AT5G22030,AT5G22200,AT5G22270,AT5G22320,AT5G22360,AT5G20190,AT5G20860,AT5G21222,AT5G22130 |
| chr4:6031362 | ATLINE1_1-LINE/L1 | AT4G09490,AT4G09500,AT4G09510,AT4G09520,AT4G09545,AT4G09550,AT4G09560,AT4G09530 |
| chr1:16975207 | ATHILA3-LTR/Gypsy | AT1G44890,AT1G44895,AT1G44910,AT1G44920,AT1G44940,AT1G44941,AT1G44960,AT1G44970,AT1G44980,AT1G44990,AT1G45000,AT1G45010,AT1G45015,AT1G45050,AT1G45063,AT1G45100,AT1G45110,AT1G45130,AT1G44900 |
| chr3:11118309 | ATCOPIA14-LTR/Copia | AT3G29130,AT3G29140,AT3G29152,AT3G29160,AT3G29170 |
| chr4:737265 | ATCOPIA92-LTR/Copia | AT4G01660,AT4G01670,AT4G01671,AT4G01680,AT4G01690,AT4G01700,AT4G01703,AT4G01710,AT4G01720,AT4G01730 |
| chr4:6777205 | ATHILA4D_LTR-LTR/Gypsy | AT4G11080,AT4G11090,AT4G11100,AT4G11110,AT4G11120,AT4G11140,AT4G11150,AT4G11160,AT4G11175,AT4G11180,AT4G11190,AT4G11210,AT4G11211,AT4G11220,AT4G11230,AT4G11250,AT4G11260,AT4G11130(RDR2),AT4G11240,AT4G11170 |
| chr2:9255919 | ATENSPM11-DNA/En-Spm | AT2G21595,AT2G21600,AT2G21610,AT2G21620,AT2G21630,AT2G21640,AT2G21650,AT2G21655,AT2G21660,AT2G21680,AT2G21690,AT2G21720,AT2G21725,AT2G21727,AT2G21730,AT2G21740,AT2G21590,AT2G21710 |
| chr1:2480692 | VANDAL10-DNA/MuDR | AT1G07910,AT1G07920,AT1G07930,AT1G07940,AT1G07950,AT1G07960,AT1G07970,AT1G07985,AT1G07990,AT1G08000,AT1G08005,AT1G08010,AT1G08035,AT1G08040,AT1G07980,AT1G08030 |
| chr3:17983921 | HELITRON1-RC/Helitron | AT3G48110,AT3G48120,AT3G48131,AT3G48140,AT3G48150,AT3G48160,AT3G48170,AT3G48180,AT3G48187,AT3G48195,AT3G48200,AT3G48205,AT3G48208,AT3G48209,AT3G48210,AT3G48220,AT3G48230,AT3G48240,AT3G48250,AT3G48270,AT3G48280,AT3G48290,AT3G48300,AT3G48298,AT3G48310,AT3G48330,AT3G48340,AT3G48343,AT3G48344,AT3G48346,AT3G48350,AT3G48360,AT3G48380,AT3G48390,AT3G48420,AT3G48425(APE1L),AT3G48440,AT3G48450,AT3G48460,AT3G48475,AT3G48480,AT3G |

| | | |
|---|---|---|
| | | 48490,AT3G48500,AT3G48510,AT3G48520,AT3G48540,AT3G48550,AT3G48100,AT3G48185,AT3G48231,AT3G48260,AT3G48320,AT3G48410,AT3G48400,AT3G48430,AT3G48530,AT3G48470,AT3G48190 |
| chr1:27652049 | ATHAT3-DNA/HAT | AT1G73370,AT1G73390,AT1G73400,AT1G73410,AT1G73440,AT1G73450,AT1G73470,AT1G73480,AT1G73490,AT1G73510,AT1G73530,AT1G73540,AT1G73550,AT1G73560,AT1G73570,AT1G73580,AT1G73590,AT1G73380,AT1G73430,AT1G73460,AT1G73500 |
| chr2:545516 | ARNOLD1-DNA/MuDR | AT2G01260,AT2G01270,AT2G01275,AT2G01290,AT2G01300,AT2G01310,AT2G01320,AT2G01330,AT2G01340,AT2G01350,AT2G01360,AT2G01370,AT2G01379,AT2G01390,AT2G01400,AT2G01410,AT2G01430,AT2G01450,AT2G01460,AT2G01480,AT2G01490,AT2G01500,AT2G01510,AT2G01520,AT2G01530,AT2G01540,AT2G01554,AT2G01560,AT2G01570,AT2G01580,AT2G01590,AT2G01600,AT2G01610,AT2G01630,AT2G01640,AT2G01650,AT2G01660,AT2G01667,AT2G01670,AT2G01680,AT2G01710,AT2G01720,AT2G01730,AT2G01735,AT2G01750,AT2G01755,AT2G01760,AT2G01770,AT2G01780,AT2G01790,AT2G01810,AT2G01818,AT2G01830,AT2G01850,AT2G01860,AT2G01870,AT2G01890,AT2G01900,AT2G01905,AT2G01910,AT2G01913,AT2G01918,AT2G01920,AT2G01930,AT2G01940,AT2G01960,AT2G01970,AT2G01990,AT2G02000,AT2G02020,AT2G02023,AT2G02026,AT2G02030,AT2G02040,AT2G02060,AT2G02061,AT2G02080,AT2G02100,AT2G02103,AT2G02120,AT2G02140,AT2G02147,AT2G02148,AT2G02150,AT2G02160,AT2G02180,AT2G01280,AT2G01420,AT2G01440,AT2G01470,AT2G01505,AT2G01620,AT2G01690,AT2G01740,AT2G01800,AT2G01820,AT2G01950,AT2G01980,AT2G02010,AT2G02050,AT2G02070,AT2G02130,AT2G02170,AT2G01880,AT2G02090 |
| chr2:4114169 | ATCOPIA32-LTR/Copia | AT2G07110,AT2G07120,AT2G07140,AT2G07170,AT2G07190,AT2G07200,AT2G07215,AT2G07240,AT2G07280,AT2G07290,AT2G07310,AT2G07340,AT2G07440,AT2G07505,AT2G07560,AT2G07565,AT2G07640,AT2G07771,AT2G07773,AT2G07655,AT2G07776,AT2G07749,AT2G07777,AT2G07671,AT2G07779,AT2G07672,AT2G07613,AT2G07684,AT2G07617,AT2G07621,AT2G07673,AT2G07674,AT2G07751,AT2G07675,AT2G07676,AT2G07768,AT2G07678,AT2G07669,AT2G07625,AT2G07681,AT2G07626,AT2G07627,AT2G07772,AT2G07628,AT2G07774,AT2G07687,AT2G07629,AT2G07631,AT2G07632,AT2G07633,AT2G07689,AT2G07634,AT2G07691,AT2G07636,AT2G07692,AT2G07637,AT2G07695,AT2G07785,AT2G07599,AT2G07696,AT2G07698,AT2G07667,AT2G07701,A |

| | | |
|---|---|---|
| | | T2G07702,AT2G07638,AT2G07705,AT2G07706,AT2G07707,AT2G07708,AT2G07641,AT2G07642,AT2G07643,AT2G07644,AT2G07713,AT2G07646,AT2G07715,AT2G07718,AT2G07648,AT2G07719,AT2G07721,AT2G07722,AT2G07815,AT2G07724,AT2G07725,AT2G07727,AT2G07728,AT2G07820,AT2G07656,AT2G07732,AT2G07825,AT2G07827,AT2G07830,AT2G07787,AT2G07658,AT2G07659,AT2G07775,AT2G07661,AT2G07806,AT2G07738,AT2G07795,AT2G07739,AT2G07662,AT2G07665,AT2G07835,AT2G07741,AT2G07690,AT2G07750,AT2G07760,AT2G07800,AT2G07810,AT2G07981,AT2G09388,AT2G09838,AT2G09840,AT2G09970,AT2G09990,AT2G10020,AT2G10025,AT2G10260,AT2G10450,AT2G10455,AT2G10535,AT2G10545,AT2G10550,AT2G10553,AT2G10556,AT2G10557,AT2G10560,AT2G10602,AT2G10608,AT2G10615,AT2G07180,AT2G07360,AT2G07798,AT2G07652,AT2G07654,AT2G07734,AT2G07680,AT2G08986,AT2G10440,AT2G07623,AT2G07714 |
| chr2:7996680 | ATCOPIA30-LTR/Copia | AT2G18300,AT2G18320,AT2G18328,AT2G18330,AT2G18340,AT2G18350,AT2G18370,AT2G18380,AT2G18390,AT2G18400,AT2G18410,AT2G18420,AT2G18450,AT2G18460,AT2G18465,AT2G18470,AT2G18480,AT2G18360 |
| chr1:27601701 | ATCOPIA21-LTR/Copia | AT1G73260,AT1G73270,AT1G73280,AT1G73300,AT1G73310,AT1G73320,AT1G73325,AT1G73330,AT1G73350,AT1G73360,AT1G73370,AT1G73390,AT1G73400,AT1G73410,AT1G73440,AT1G73450,AT1G73470,AT1G73480,AT1G73490,AT1G73510,AT1G73530,AT1G73540,AT1G73550,AT1G73560,AT1G73570,AT1G73580,AT1G73590,AT1G73290,AT1G73340,AT1G73380,AT1G73430,AT1G73460,AT1G73500 |
| chr2:7995404 | HELITRONY1C-RC/Helitron | AT2G18328,AT2G18330,AT2G18340,AT2G18350,AT2G18370,AT2G18380,AT2G18390,AT2G18400,AT2G18410,AT2G18420,AT2G18450,AT2G18460,AT2G18465,AT2G18470,AT2G18480,AT2G18490,AT2G18510,AT2G18520,AT2G18530,AT2G18550,AT2G18570,AT2G18590,AT2G18600,AT2G18620,AT2G18630,AT2G18640,AT2G18650,AT2G18660,AT2G18680,AT2G18685,AT2G18690,AT2G18360,AT2G18500,AT2G18540,AT2G18560,AT2G18670,AT2G18700 |
| chr3:16601301 | TNAT2A-DNA | AT3G44910,AT3G44920,AT3G44930,AT3G44935,AT3G44940,AT3G44950,AT3G44960,AT3G44980,AT3G44990,AT3G45000,AT3G45010,AT3G45020,AT3G45030,AT3G45040,AT3G45050,AT3G45060,AT3G45070,AT3G45080,AT3G45090,AT3G45093,AT3G45100,AT3G45110,AT3G45140,AT3G45160,AT3G45170,AT3G45180,AT3 |

| | | |
|---|---|---|
| | | G45200,AT3G45210,AT3G45220,AT3G45230,AT3G45240,AT3G45243,AT3G45245,AT3G45248,AT3G45252,AT3G45275,AT3G45285,AT3G45290,AT3G44970,AT3G45150,AT3G45190,AT3G45260,AT3G45280,AT3G45130 |
| chr1:22226311 | ATREP14-RC/Helitron | AT1G60220,AT1G60230,AT1G60240,AT1G60250,AT1G60260,AT1G60270,AT1G60280,AT1G60300,AT1G60320,AT1G60340,AT1G60350,AT1G60360,AT1G60370,AT1G60380,AT1G60400,AT1G60410,AT1G60390 |
| chr5:13350109 | ATCOPIA15-LTR/Copia | AT5G35050,AT5G35067,AT5G35080,AT5G35090,AT5G35100,AT5G35110,AT5G35069,AT5G34940 |
| chr1:16120148 | ATHILA2-LTR/Gypsy | AT1G42960,AT1G42970,AT1G42980,AT1G42990,AT1G43000,AT1G43005,AT1G43010,AT1G43020,AT1G43040,AT1G43080,AT1G43090,AT1G43100,AT1G43130,AT1G43140,AT1G43145,AT1G43160,AT1G43170,AT1G43190,AT1G43245,AT1G43310,AT1G43320,AT1G43330,AT1G43415,AT1G43560,AT1G43580,AT1G43600,AT1G43610,AT1G43620,AT1G43630,AT1G43640,AT1G43650,AT1G43665,AT1G43666,AT1G43667,AT1G43670,AT1G43680,AT1G43700,AT1G43710,AT1G43720,AT1G43722,AT1G43730,AT1G43760,AT1G43770,AT1G43790,AT1G43800,AT1G43815,AT1G43825,AT1G43850,AT1G43860,AT1G43171,AT1G43260,AT1G43605,AT1G43690,AT1G43780,AT1G43810 |
| chr3:10761315 | DT1-DNA/Mariner | AT3G28670,AT3G28680,AT3G28690,AT3G28700,AT3G28710,AT3G28674 |
| chr1:540339 | ATCOPIA32B-LTR/Copia | AT1G02475,AT1G02490,AT1G02500,AT1G02510,AT1G02530,AT1G02540,AT1G02550,AT1G02560,AT1G02570,AT1G02575,AT1G02580,AT1G02590,AT1G02610,AT1G02620,AT1G02630,AT1G02520 |
| chr1:12817581 | ATCOPIA36-LTR/Copia | AT1G34840,AT1G34850,AT1G34855,AT1G34860,AT1G34910,AT1G34930,AT1G35030,AT1G35035,AT1G35140,AT1G35150,AT1G35160,AT1G35170,AT1G35181,AT1G35040,AT1G35180 |
| chr1:12817581 | ATCOPIA57-LTR/Copia | AT1G33770,AT1G33780,AT1G33790,AT1G33800,AT1G33810,AT1G33811,AT1G33820,AT1G33830,AT1G33840,AT1G33860,AT1G33870,AT1G33890,AT1G33900,AT1G33910,AT1G33930,AT1G33940,AT1G33945,AT1G33960,AT1G33970,AT1G33990,AT1G34000,AT1G34010,AT1G34015,AT1G34030,AT1G34040,AT1G34042,AT1G34046,AT1G34047,AT1G34049,AT1G34050,AT1G34060,AT1G34065,AT1G34095,AT1G34120,AT1G34130,AT1G34140,AT1G34150,AT1G34160,AT1G34170,AT1G34180,AT1G34200,AT1G34210,AT1G34220,AT1G34245,AT1G34260,AT1G34270,AT1G34290,AT1 |

| | | |
|---|---|---|
| | | G34300,AT1G34310,AT1G34315,AT1G34317,AT1G34320,AT1G34350,AT1G34355,AT1G34360,AT1G34370,AT1G34380,AT1G34390,AT1G34403,AT1G34410,AT1G34420,AT1G34430,AT1G34440,AT1G34470,AT1G34480,AT1G34490,AT1G34500,AT1G34510,AT1G34520,AT1G34540,AT1G34560,AT1G34570,AT1G34575,AT1G34580,AT1G34630,AT1G34640,AT1G34670,AT1G34750,AT1G34760,AT1G34780,AT1G34790,AT1G34792,AT1G34795,AT1G34800,AT1G34805,AT1G34807,AT1G34810,AT1G34812,AT1G34815,AT1G34817,AT1G34820,AT1G34822,AT1G34825,AT1G34827,AT1G34830,AT1G34840,AT1G34850,AT1G34855,AT1G34860,AT1G34910,AT1G34930,AT1G35030,AT1G35035,AT1G35140,AT1G35150,AT1G35160,AT1G35170,AT1G35181,AT1G35183,AT1G35190,AT1G35210,AT1G35215,AT1G35230,AT1G35240,AT1G35242,AT1G35250,AT1G35255,AT1G35260,AT1G35290,AT1G35310,AT1G35330,AT1G35340,AT1G35350,AT1G35353,AT1G35365,AT1G35375,AT1G35400,AT1G35410,AT1G35420,AT1G35430,AT1G35435,AT1G35440,AT1G35467,AT1G35470,AT1G35490,AT1G35500,AT1G35515,AT1G35516,AT1G35520,AT1G35537,AT1G35540,AT1G35550,AT1G35560,AT1G35610,AT1G35614,AT1G35617,AT1G35625,AT1G35630,AT1G35660,AT1G35680,AT1G35720,AT1G35730,AT1G35780,AT1G35820,AT1G35830,AT1G35850,AT1G35860,AT1G35880,AT1G35890,AT1G35910,AT1G36000,AT1G36005,AT1G36020,AT1G36030,AT1G36060,AT1G36078,AT1G36085,AT1G36095,AT1G36100,AT1G36150,AT1G36230,AT1G36272,AT1G36280,AT1G36310,AT1G36320,AT1G36325,AT1G36340,AT1G36380,AT1G36390,AT1G36580,AT1G36622,AT1G36623,AT1G36627,AT1G36640,AT1G36675,AT1G36730,AT1G36745,AT1G36756,AT1G36920,AT1G36922,AT1G36925,AT1G36940,AT1G36942,AT1G36950,AT1G36960,AT1G36970,AT1G36990,AT1G37000,AT1G37020,AT1G37113,AT1G37130,AT1G37140,AT1G37150,AT1G38065,AT1G38131,AT1G33850,AT1G33880,AT1G33950,AT1G33980,AT1G34020,AT1G34070,AT1G34110,AT1G34340,AT1G34400,AT1G34460,AT1G34550,AT1G34650,AT1G34770,AT1G35040,AT1G35180,AT1G35220,AT1G35320,AT1G35460,AT1G35510,AT1G35530,AT1G35580,AT1G35620,AT1G35670,AT1G35710,AT1G35750,AT1G35895,AT1G36050,AT1G36070,AT1G36160,AT1G36180,AT1G36240,AT1G36370,AT1G37010,AT1G33920,AT1G34190,AT1G36510,AT1G36980 |
| chr4:7881096 | ATGP3-LTR/Gypsy | AT4G13530,AT4G13540,AT4G13550,AT4G13570,AT4G13572,AT4G13575,AT4G13560,AT4G13577 |

| | | |
|---|---|---|
| chr4:5530600 | HELITRONY3-RC/Helitron | AT4G08685,AT4G08690,AT4G08691,AT4G08700,AT4G08670 |
| chr1:23588573 | VANDALNX2-DNA/MuDR | AT1G63206,AT1G63220,AT1G63230,AT1G63240,AT1G63245,AT1G63250,AT1G63260,AT1G63270,AT1G63280,AT1G63290,AT1G63295,AT1G63300,AT1G63310,AT1G63320,AT1G63330,AT1G63340;Note=Flavin-containing monooxygenase family protein,AT1G63360,AT1G63370,AT1G63380,AT1G63390,AT1G63400,AT1G63420,AT1G63430,AT1G63440,AT1G63450,AT1G63460,AT1G63470,AT1G63480,AT1G63490,AT1G63500,AT1G63522,AT1G63530,AT1G63535,AT1G63540,AT1G63550,AT1G63570,AT1G63580,AT1G63590,AT1G63600,AT1G63610,AT1G63615,AT1G63630,AT1G63650,AT1G63210,AT1G63350,AT1G63410,AT1G63520,AT1G63640 |
| chr4:972969 | HELITRONY1C-RC/Helitron | AT4G02150,AT4G02160,AT4G02170,AT4G02180,AT4G02190,AT4G02195,AT4G02200,AT4G02210,AT4G02220,AT4G02230,AT4G02235,AT4G02250,AT4G02260,AT4G02270,AT4G02280 |
| chr2:9843129 | ATMU6-DNA/MuDR | AT2G23080,AT2G23093,AT2G23096,AT2G23100,AT2G23110,AT2G23120,AT2G23118,AT2G23130,AT2G23142,AT2G23148,AT2G23150,AT2G23140,AT2G23090 |
| chr5:18849826 | ATCOPIA35-LTR/Copia | AT5G46340,AT5G46350,AT5G46360,AT5G46370,AT5G46380,AT5G46390,AT5G46395,AT5G46400,AT5G46420,AT5G46430,AT5G46440,AT5G46450,AT5G46470,AT5G46490,AT5G46510,AT5G46410,AT5G46460,AT5G46500 |
| chr4:10793777 | ATCOPIA28-LTR/Copia | AT4G19820,AT4G19830,AT4G19840,AT4G19850,AT4G19860,AT4G19865,AT4G19870,AT4G19880,AT4G19890,AT4G19900,AT4G19905,AT4G19910,AT4G19920,AT4G19925,AT4G19930,AT4G19940 |
| chr1:29398093 | ATHILA4C-LTR/Gypsy | AT1G78100,AT1G78120,AT1G78130,AT1G78140,AT1G78150,AT1G78170,AT1G78172,AT1G78180,AT1G78190,AT1G78200,AT1G78210,AT1G78220,AT1G78230,AT1G78240,AT1G78110,AT1G78160 |
| chr1:29398093 | VANDAL4-DNA/MuDR | AT1G78100,AT1G78120,AT1G78130,AT1G78140,AT1G78150,AT1G78170,AT1G78172,AT1G78180,AT1G78190,AT1G78200,AT1G78210,AT1G78220,AT1G78230,AT1G78240,AT1G78110,AT1G78160 |
| chr5:10476340 | ATLINE1A-LINE/L1 | AT5G28490,AT5G28491,AT5G28500,AT5G28510 |

| | | |
|---|---|---|
| chr1:26513099 | ATREP2-RC/Helitron | AT1G70330,AT1G70335,AT1G70340,AT1G70350,AT1G70370,AT1G70380,AT1G70390,AT1G70320,AT1G70360 |
| chr2:11414196 | ATCOPIA65-LTR/Copia | AT2G25470,AT2G25480,AT2G25490,AT2G25500,AT2G25510,AT2G25520,AT2G25530,AT2G25540,AT2G25560,AT2G25565,AT2G25570,AT2G25580,AT2G25590,AT2G25605,AT2G25610,AT2G25620,AT2G25625,AT2G25630,AT2G25650,AT2G25660,AT2G25670,AT2G25680,AT2G25685,AT2G25690,AT2G25700,AT2G25710,AT2G25720,AT2G25735,AT2G25740,AT2G25760,AT2G25770,AT2G25780,AT2G25790,AT2G25800,AT2G25810,AT2G25820,AT2G25830,AT2G25840,AT2G25870,AT2G25880,AT2G25890,AT2G25905,AT2G25910,AT2G25920,AT2G25940,AT2G25950,AT2G25964,AT2G25980,AT2G25990,AT2G26000,AT2G26010,AT2G26020,AT2G26040,AT2G26050,AT2G26060,AT2G26070,AT2G26080,AT2G26100,AT2G26110,AT2G26120,AT2G26130,AT2G26135,AT2G26140,AT2G26150,AT2G26160,AT2G26180,AT2G26190,AT2G26200,AT2G26230,AT2G26240,AT2G26250,AT2G26260,AT2G26270,AT2G26290,AT2G26300,AT2G26310,AT2G26320,AT2G26330,AT2G26340,AT2G26350,AT2G26370,AT2G26380,AT2G26390,AT2G26400,AT2G26410,AT2G26430,AT2G26440,AT2G26450,AT2G26470,AT2G26480,AT2G26490,AT2G26500,AT2G26510,AT2G26515,AT2G26520,AT2G26530,AT2G26550,AT2G26560,AT2G26570,AT2G26590,AT2G26600,AT2G26620,AT2G26640,AT2G26650,AT2G26660,AT2G26670,AT2G26680,AT2G26690,AT2G26695,AT2G26710,AT2G26720,AT2G26730,AT2G26740,AT2G26750,AT2G26770,AT2G26790,AT2G26800,AT2G26820,AT2G26830,AT2G25482,AT2G25600,AT2G25640,AT2G25730,AT2G25737,AT2G25850,AT2G25900,AT2G25930,AT2G25970,AT2G26030,AT2G26210,AT2G26280,AT2G26360,AT2G26420,AT2G26460,AT2G26540,AT2G26580,AT2G26610,AT2G26700,AT2G26780,AT2G26810,AT2G26170,AT2G26760 |
| chr3:16840925 | ATLANTYS2-LTR/Gypsy | AT3G45790,AT3G45800,AT3G45810,AT3G45820,AT3G45840,AT3G45851,AT3G45860,AT3G45870,AT3G45880,AT3G45890,AT3G45910,AT3G45830,AT3G45850,AT3G45900 |
| chr2:5800623 | ATHAT10-DNA/HAT | AT2G13840,AT2G13845,AT2G13895,AT2G13900,AT2G13905 |
| chr4:6409588 | TAG1-DNA/HAT | AT4G10300,AT4G10305,AT4G10310,AT4G10320,AT4G10330,AT4G10340,AT4G10350,AT4G10360,AT4G10370,AT4G10380 |
| chr3:17161175 | ATCOPIA19-LTR/Copia | AT3G45140,AT3G45160,AT3G45170,AT3G45180,AT3G45200,AT3G45210,AT3G45220,AT3G45230,AT3G45240,AT3G45243,AT3G4 |

| | | |
|---|---|---|
| | | 5245,AT3G45248,AT3G45252,AT3G45275,AT3G45285,AT3G45290,AT3G45300,AT3G45320,AT3G45330,AT3G45390,AT3G45400,AT3G45410,AT3G45420,AT3G45440,AT3G45443,AT3G45450,AT3G45470,AT3G45480,AT3G45490,AT3G45500,AT3G45510,AT3G45525,AT3G45530,AT3G45540,AT3G45555,AT3G45560,AT3G45570,AT3G45577,AT3G45580,AT3G45590,AT3G45600,AT3G45610,AT3G45620,AT3G45630,AT3G45640,AT3G45645,AT3G45660,AT3G45670,AT3G45673,AT3G45680,AT3G45700,AT3G45710,AT3G45720,AT3G45730,AT3G45740,AT3G45760,AT3G45770,AT3G45780,AT3G45790,AT3G45800,AT3G45810,AT3G45820,AT3G45840,AT3G45851,AT3G45860,AT3G45870,AT3G45880,AT3G45890,AT3G45910,AT3G45920,AT3G45930,AT3G45940,AT3G45950,AT3G45960,AT3G45970,AT3G45980,AT3G45990,AT3G46010,AT3G46020,AT3G46030,AT3G46040,AT3G46050,AT3G46060,AT3G46070,AT3G46080,AT3G46086,AT3G46090,AT3G46100,AT3G46110,AT3G46120,AT3G46130,AT3G46140,AT3G46150,AT3G46160,AT3G46170,AT3G46180,AT3G46190,AT3G46200,AT3G46210,AT3G46220,AT3G46230,AT3G46240,AT3G46260,AT3G46270,AT3G46290,AT3G46300,AT3G46310,AT3G46320,AT3G46340,AT3G46355,AT3G46360,AT3G46370,AT3G46380,AT3G46390,AT3G46400,AT3G46410,AT3G46420,AT3G46440,AT3G46450,AT3G46460,AT3G46470,AT3G46490,AT3G46510,AT3G46520,AT3G46530,AT3G46540,AT3G46560,AT3G46565,AT3G46570,AT3G46580,AT3G46600,AT3G46610,AT3G46613,AT3G46616,AT3G46617,AT3G46620,AT3G46630,AT3G46660,AT3G46666,AT3G46670,AT3G46680,AT3G46690,AT3G46700,AT3G46710,AT3G46720,AT3G46730,AT3G46735,AT3G46740,AT3G46750,AT3G46760,AT3G46770,AT3G46780,AT3G46790,AT3G46810,AT3G46820,AT3G46830,AT3G46845,AT3G46850,AT3G45150,AT3G45190,AT3G45260,AT3G45280,AT3G45310,AT3G45430,AT3G45460,AT3G45650,AT3G45750,AT3G45830,AT3G45850,AT3G45900,AT3G46280,AT3G46330,AT3G46430,AT3G46480,AT3G46500,AT3G46550,AT3G46590,AT3G46640,AT3G46650,AT3G46800,AT3G46840,AT3G45130,AT3G46000,AT3G46350,AT3G45690 |
| chr1:13210878 | ATREP3-RC/Helitron | AT1G35660,AT1G35680,AT1G35720,AT1G35730,AT1G35670,AT1G35710 |
| chr5:14686374 | ATCOPIA28-LTR/Copia | AT5G37130,AT5G37140,AT5G37150 |
| chr2:7230176 | ATCOPIA70-LTR/Copia | AT2G16640,AT2G16650,AT2G16660,AT2G16676,AT2G16700,AT2G16710 |

| | | |
|---|---|---|
| chr3:23071659 | ATRE1-LTR/Copia | AT3G62260,AT3G62270,AT3G62280,AT3G62300,AT3G62310,AT3G62320,AT3G62330,AT3G62340,AT3G62350,AT3G62360,AT3G62370,AT3G62380,AT3G62290 |
| chr1:19586307 | ATCOPIA81-LTR/Copia | AT1G52460,AT1G52470,AT1G52490,AT1G52495,AT1G52520,AT1G52530,AT1G52540,AT1G52550,AT1G52560,AT1G52565,AT1G52570,AT1G52580,AT1G52590,AT1G52600,AT1G52603,AT1G52618,AT1G52620,AT1G52630,AT1G52640,AT1G52660,AT1G52670,AT1G52680,AT1G52690,AT1G52695,AT1G52700,AT1G52710,AT1G52720,AT1G52740(HTA9),AT1G52750,AT1G52760,AT1G52770,AT1G52790,AT1G52800,AT1G52810,AT1G52820,AT1G52825,AT1G52827,AT1G52830,AT1G52857,AT1G52855,AT1G52870,AT1G52880,AT1G52890,AT1G52900,AT1G52905,AT1G52910,AT1G52930,AT1G52940,AT1G52970,AT1G52980,AT1G53000,AT1G53010,AT1G53023,AT1G53030,AT1G53035,AT1G53040,AT1G53060,AT1G53070,AT1G53080,AT1G53090,AT1G53100,AT1G53120,AT1G53130,AT1G53140,AT1G53160,AT1G53163,AT1G53165,AT1G53170,AT1G53180,AT1G53190,AT1G53210,AT1G53230,AT1G53240,AT1G53260,AT1G53265,AT1G53270,AT1G53280,AT1G53282,AT1G53285,AT1G53290,AT1G53300,AT1G53310,AT1G53320,AT1G53325,AT1G53330,AT1G53340,AT1G53345,AT1G53360,AT1G53366,AT1G53370,AT1G53380,AT1G53400,AT1G53420,AT1G53440,AT1G53450,AT1G53470,AT1G53480,AT1G53490,AT1G53500,AT1G53520,AT1G53530,AT1G53540,AT1G53541,AT1G53542,AT1G53550,AT1G53560,AT1G53580,AT1G53590,AT1G53600,AT1G52450,AT1G52500,AT1G52510,AT1G52650,AT1G52730,AT1G52920,AT1G52950,AT1G52990,AT1G53025,AT1G53050,AT1G53200,AT1G53250,AT1G53350,AT1G53430,AT1G53460,AT1G53510,AT1G53570,AT1G52780,AT1G53110,AT1G53390 |
| chr5:19235647 | ATCOPIA61-LTR/Copia | AT5G47350,AT5G47360,AT5G47370,AT5G47380,AT5G47390,AT5G47400,AT5G47430,AT5G47435,AT5G47440,AT5G47450,AT5G47460,AT5G47470,AT5G47480,AT5G47420,AT5G47455,AT5G47490 |
| chr2:19062038 | ATSINE2A-SINE | AT2G46400,AT2G46410,AT2G46430,AT2G46440,AT2G46450,AT2G46460,AT2G46470,AT2G46480,AT2G46490,AT2G46493,AT2G46494,AT2G46495,AT2G46505,AT2G46510,AT2G46420,AT2G46455,AT2G46500 |
| chr2:19142628 | ATHILA2-LTR/Gypsy | AT2G46360,AT2G46370,AT2G46380,AT2G46390,AT2G46400,AT2G46410,AT2G46430,AT2G46440,AT2G46450,AT2G46460,AT2G46470,AT2G46480,AT2G46490,AT2G46493,AT2G46494,AT2G46495,AT2G46505,AT2G46510,AT2G46520,AT2G46530,AT2G46535,A |

| | | |
|---|---|---|
| | | T2G46540,AT2G46550,AT2G46570,AT2G46567,AT2G46580,AT2G46590,AT2G46610,AT2G46620,AT2G46630,AT2G46640,AT2G46650,AT2G46660,AT2G46670,AT2G46680,AT2G46690,AT2G46710,AT2G46720,AT2G46735,AT2G46740,AT2G46750,AT2G46760,AT2G46765,AT2G46770,AT2G46780,AT2G46800,AT2G46810,AT2G46820,AT2G46830,AT2G46840,AT2G46860,AT2G46870,AT2G46375,AT2G46420,AT2G46455,AT2G46500,AT2G46560,AT2G46700,AT2G46790,AT2G46850,AT2G46600 |
| chr4:2718352 | VANDAL5-DNA/MuDR | AT4G05100,AT4G05110,AT4G05120,AT4G05140,AT4G05150,AT4G05160,AT4G05170,AT4G05180,AT4G05190,AT4G05200,AT4G05210,AT4G05220,AT4G05230,AT4G05240,AT4G05250,AT4G05260,AT4G05270,AT4G05310,AT4G05330,AT4G05340,AT4G05350,AT4G05360,AT4G05390,AT4G05400,AT4G05410,AT4G05430,AT4G05440,AT4G05450,AT4G05460,AT4G05470,AT4G05475,AT4G05490,AT4G05497,AT4G05130,AT4G05320,AT4G05380,AT4G05420 |
| chr2:10736036 | VANDAL2N1-DNA/MuDR | AT2G25180,AT2G25185,AT2G25190,AT2G25200,AT2G25210,AT2G25215,AT2G25125,AT2G25220,AT2G25240,AT2G25170(PKL),AT2G25230 |
| chr2:13189113 | BRODYAGA1A-DNA/MuDR | AT2G30940,AT2G30942,AT2G30950,AT2G30960,AT2G30970,AT2G30980,AT2G30985,AT2G31005,AT2G31010,AT2G31020,AT2G31018,AT2G31030,AT2G31040,AT2G31035,AT2G31050,AT2G31060,AT2G30990,AT2G31070 |
| chr1:21222585 | ATENSPM9-DNA/En-Spm | AT1G56600,AT1G56610,AT1G56620,AT1G56630,AT1G56650 |
| chr4:11307964 | TSCL-LINE? | AT4G21180,AT4G21190,AT4G21192,AT4G21210,AT4G21213,AT4G21215,AT4G21220,AT4G21240,AT4G21200,AT4G21230 |
| chr1:17853243 | ATREP6-RC/Helitron | AT1G48280,AT1G48285,AT1G48300,AT1G48310,AT1G48320,AT1G48325,AT1G48330,AT1G48350,AT1G48355,AT1G48360 |
| chr3:3937527 | ATCOPIA89-LTR/Copia | AT3G12050,AT3G12060,AT3G12070,AT3G12090,AT3G12100,AT3G12110,AT3G12120,AT3G12140,AT3G12145,AT3G12150,AT3G12160,AT3G12170,AT3G12180,AT3G12190,AT3G12200,AT3G12203,AT3G12210,AT3G12220,AT3G12240,AT3G12250,AT3G12260,AT3G12270,AT3G12290(MTHFD1),AT3G12300,AT3G12320,AT3G12340,AT3G12345,AT3G12360,AT3G12370,AT3G12380,AT3G12390,AT3G12400,AT3G12410,AT3G12430,AT3G12440,AT3G12460,AT3G12470,AT3G12480,AT3G12490,AT3G12500,AT3G12510,AT3G12520,AT3G12530,AT3G12540,AT3G12545,AT3G12560,AT3G12570,AT3G12580,AT3G12587,AT3G12600,AT3G12610,AT3G1262 |

| | | |
|---|---|---|
| | | 0,AT3G12630,AT3G12650,AT3G12670,AT3G12660,AT3G12680,AT3G12685,AT3G12700,AT3G12710,AT3G12720,AT3G12730,AT3G12740,AT3G12750,AT3G12760,AT3G12770,AT3G12775,AT3G12800,AT3G12810(PIE1),AT3G12820,AT3G12833,AT3G12835,AT3G12840,AT3G12850,AT3G12860,AT3G12870,AT3G12890,AT3G12900,AT3G12910,AT3G12915,AT3G12920,AT3G12940,AT3G12950,AT3G12955,AT3G12960,AT3G12970,AT3G12980,AT3G12990,AT3G13010,AT3G13020,AT3G13030,AT3G13040,AT3G13060,AT3G13062,AT3G13065,AT3G13075,AT3G13080,AT3G13090,AT3G13110,AT3G13120,AT3G13130,AT3G13150,AT3G12080,AT3G12130,AT3G12230,AT3G12280,AT3G12420,AT3G12550(FDM3),AT3G12590,AT3G12640,AT3G12690,AT3G12830,AT3G12880,AT3G12930,AT3G12977,AT3G13000,AT3G13050,AT3G13100,AT3G13140,AT3G12350,AT3G12780,AT3G13070 |
| chr4:2656886 | ATCOPIA76-LTR/Copia | AT4G05140,AT4G05150,AT4G05160,AT4G05170,AT4G05180,AT4G05130 |
| chr1:11453796 | VANDAL6-DNA/MuDR | AT1G31870,AT1G31880,AT1G31885,AT1G31910,AT1G31920,AT1G31940,AT1G31950,AT1G31860,AT1G31930 |
| chr5:13684120 | ATCOPIA24-LTR/Copia | AT5G35200,AT5G35210,AT5G35220,AT5G35230,AT5G35300,AT5G35320,AT5G35330,AT5G35360,AT5G35370,AT5G35375,AT5G35380,AT5G35400,AT5G35405,AT5G35410,AT5G35450,AT5G35460,AT5G35475,AT5G35480,AT5G35490,AT5G35510,AT5G35520,AT5G35525,AT5G35530,AT5G35540,AT5G35550,AT5G35570,AT5G35338,AT5G35390,AT5G35430,AT5G35560 |
| chr4:17321503 | ATCOPIA46-LTR/Copia | AT4G36680,AT4G36690,AT4G36700,AT4G36710,AT4G36720,AT4G36730,AT4G36740,AT4G36750,AT4G36760,AT4G36770,AT4G36790,AT4G36791,AT4G36795,AT4G36800,AT4G36810,AT4G36820,AT4G36840,AT4G36780,AT4G36830 |
| chr4:7839065 | ATCOPIA63-LTR/Copia | AT4G13350,AT4G13360,AT4G13370,AT4G13390,AT4G13395,AT4G13400,AT4G13410,AT4G13430,AT4G13440,AT4G13460(SUVH9),AT4G13480,AT4G13490,AT4G13500,AT4G13510,AT4G13530,AT4G13540,AT4G13550,AT4G13570,AT4G13572,AT4G13575,AT4G13580,AT4G13590,AT4G13600,AT4G13610,AT4G13615,AT4G13620,AT4G13630,AT4G13640,AT4G13650,AT4G13670,AT4G13680,AT4G13690,AT4G13700,AT4G13720,AT4G13730,AT4G13380,AT4G13420,AT4G13450,AT4G13560,AT4G13577,AT4G13660,AT4G13710,AT4G13520 |

| | | |
|---|---|---|
| chr2:7298605 | VANDAL14-DNA/MuDR | AT2G16760,AT2G16770,AT2G16780,AT2G16790,AT2G16800,AT2G16810,AT2G16835,AT2G16850,AT2G16860,AT2G16870,AT2G16880,AT2G16750 |
| chr5:17403448 | ATSINE2A-SINE | AT5G43300,AT5G43320,AT5G43330,AT5G43340,AT5G43360,AT5G43380,AT5G43390,AT5G43400,AT5G43401,AT5G43310,AT5G43350,AT5G43370 |
| chr1:25354273 | TSCL-LINE? | AT1G67265,AT1G67270,AT1G67280,AT1G67290,AT1G67300,AT1G67320,AT1G67325,AT1G67350,AT1G67360,AT1G67370,AT1G67390,AT1G67400,AT1G67410,AT1G67420,AT1G67430,AT1G67450,AT1G67455,AT1G67460,AT1G67470,AT1G67480,AT1G67510,AT1G67520,AT1G67530,AT1G67540,AT1G67560,AT1G67570,AT1G67590,AT1G67600,AT1G67620,AT1G67623,AT1G67635,AT1G67640,AT1G67660,AT1G67670,AT1G67680,AT1G67690,AT1G67700,AT1G67720,AT1G67730,AT1G67310,AT1G67330,AT1G67340,AT1G67440,AT1G67490,AT1G67550,AT1G67580,AT1G67630,AT1G67650,AT1G67710,AT1G67740,AT1G67500 |
| chr2:767986 | ATHILA8A-LTR/Gypsy | AT2G02700,AT2G02710,AT2G02720,AT2G02730,AT2G02750,AT2G02760,AT2G02765,AT2G02770,AT2G02780,AT2G02695,AT2G02740 |
| chr2:2151176 | ATHILA6A-LTR/Gypsy | AT2G05720,AT2G05752,AT2G05755,AT2G05753,AT2G05710 |
| chr3:18730206 | ATSINE4-SINE | AT3G50430,AT3G50440,AT3G50450,AT3G50460,AT3G50470,AT3G50480,AT3G50510,AT3G50520,AT3G50530,AT3G50540,AT3G50550,AT3G50500 |
| chr2:4007630 | ATCOPIA32B-LTR/Copia | AT2G07110,AT2G07120,AT2G07140,AT2G07170,AT2G07190,AT2G07200,AT2G07215,AT2G07240,AT2G07280,AT2G07290,AT2G07310,AT2G07340,AT2G07440,AT2G07505,AT2G07560,AT2G07565,AT2G07640,AT2G07771,AT2G07773,AT2G07655,AT2G07776,AT2G07749,AT2G07777,AT2G07671,AT2G07779,AT2G07672,AT2G07613,AT2G07684,AT2G07617,AT2G07621,AT2G07673,AT2G07674,AT2G07751,AT2G07675,AT2G07676,AT2G07768,AT2G07678,AT2G07669,AT2G07625,AT2G07681,AT2G07626,AT2G07627,AT2G07772,AT2G07628,AT2G07774,AT2G07687,AT2G07629,AT2G07631,AT2G07632,AT2G07633,AT2G07689,AT2G07634,AT2G07691,AT2G07636,AT2G07692,AT2G07637,AT2G07695,AT2G07785,AT2G07599,AT2G07696,AT2G07698,AT2G07667,AT2G07701,AT2G07702,AT2G07638,AT2G07705,AT2G07706,AT2G07707,AT2G07708,AT2G07641,AT2G07642,AT2G07643,AT2G07644,AT2G07713,AT2G07646,AT2G07715,AT2G07718,AT2G07648,AT2G0771 |

| | | |
|---|---|---|
| | | 9,AT2G07721,AT2G07722,AT2G07815,AT2G07724,AT2G07725,AT2G07727,AT2G07728,AT2G07820,AT2G07656,AT2G07732,AT2G07825,AT2G07827,AT2G07830,AT2G07787,AT2G07658,AT2G07659,AT2G07775,AT2G07661,AT2G07806,AT2G07738,AT2G07795,AT2G07739,AT2G07662,AT2G07665,AT2G07835,AT2G07741,AT2G07690,AT2G07750,AT2G07760,AT2G07800,AT2G07810,AT2G07981,AT2G09388,AT2G09838,AT2G09840,AT2G09970,AT2G09990,AT2G10020,AT2G10025,AT2G10260,AT2G10450,AT2G10455,AT2G10535,AT2G10545,AT2G10550,AT2G10553,AT2G10556,AT2G10557,AT2G10560,AT2G10602,AT2G10608,AT2G10615,AT2G10625,AT2G10920,AT2G10930,AT2G10931,AT2G10940,AT2G10950,AT2G10955,AT2G10965,AT2G10970,AT2G10975,AT2G11005,AT2G11010,AT2G11015,AT2G11025,AT2G07180,AT2G07360,AT2G07798,AT2G07652,AT2G07654,AT2G07734,AT2G07680,AT2G08986,AT2G10440,AT2G11000,AT2G07623,AT2G07714 |
| chr3:15887401 | VANDAL16 | AT3G43990,AT3G44006,AT3G44010,AT3G44020,AT3G44050,AT3G44060,AT3G44070,AT3G44080,AT3G44090,AT3G44100,AT3G44110,AT3G44115,AT3G44120,AT3G44130,AT3G44140,AT3G44150,AT3G44160,AT3G44170,AT3G44180,AT3G44190,AT3G44210,AT3G44220,AT3G44230,AT3G44235,AT3G44240,AT3G44250,AT3G44260,AT3G44261,AT3G44265,AT3G44280,AT3G44300,AT3G44310,AT3G44320,AT3G44326,AT3G44200,AT3G44290,AT3G44330 |
| chr3:15507824 | ATCOPIA34-LTR/Copia | AT3G43470,AT3G43480,AT3G43490,AT3G43500,AT3G43505,AT3G43520,AT3G43540,AT3G43550,AT3G43570,AT3G43572,AT3G43574,AT3G43580,AT3G43583,AT3G43590,AT3G43610,AT3G43440,AT3G43503,AT3G43600 |
| chr3:15507824 | TNAT1A-DNA | AT3G43432,AT3G43470,AT3G43480,AT3G43490,AT3G43500,AT3G43505,AT3G43520,AT3G43540,AT3G43550,AT3G43570,AT3G43572,AT3G43574,AT3G43580,AT3G43583,AT3G43590,AT3G43610,AT3G43440,AT3G43503,AT3G43600 |
| chr4:6736283 | ATREP13-RC/Helitron | AT4G10660,AT4G10670,AT4G10680,AT4G10695,AT4G10700,AT4G10720,AT4G10730,AT4G10740,AT4G10760,AT4G10767,AT4G10770,AT4G10790,AT4G10800,AT4G10810,AT4G10820,AT4G10840,AT4G10843,AT4G10850,AT4G10860,AT4G10870,AT4G10880,AT4G10890,AT4G10895,AT4G10910,AT4G10920,AT4G10925,AT4G10930,AT4G10950,AT4G10955,AT4G10970,AT4G11000,AT4G11020,AT4G11030,AT4G11040,AT4G11060,AT4G11070,AT4G11080,AT4G11090,AT4G10710(SPT16),AT4G10750,AT4G10780,AT4G10960,AT4G11010,AT4G11050 |

| | | |
|---|---|---|
| chr4:15344984 | ATHILA8A-LTR/Gypsy | AT4G31400,AT4G31405,AT4G31410,AT4G31420,AT4G31430,AT4G31440,AT4G31441,AT4G31450,AT4G31460,AT4G31470,AT4G31480,AT4G31500,AT4G31510,AT4G31520,AT4G31550,AT4G31560,AT4G31580,AT4G31590,AT4G31600,AT4G31615,AT4G31620,AT4G31630,AT4G31640,AT4G31650,AT4G31670,AT4G31680,AT4G31685,AT4G31690,AT4G31700,AT4G31715,AT4G31720,AT4G31390,AT4G31490,AT4G31530,AT4G31540,AT4G31570,AT4G31610,AT4G31710,AT4G31660 |
| chr5:20202759 | BRODYAGA1-DNA/MuDR | AT5G49690,AT5G49700,AT5G49710,AT5G49720,AT5G49740,AT5G49750,AT5G49730,AT5G49760,AT5G49680 |
| chr3:10978600 | TA11-LINE/L1 | AT3G28922,AT3G28925,AT3G28940,AT3G28956,AT3G28958,AT3G28960,AT3G28970,AT3G28930,AT3G28950,AT3G28980 |
| chr4:10339233 | ATCOPIA29-LTR/Copia | AT4G18810,AT4G18820,AT4G18823,AT4G18860,AT4G18870,AT4G18880,AT4G18900,AT4G18905,AT4G18910,AT4G18920,AT4G18940,AT4G18950,AT4G18970,AT4G18975,AT4G18980,AT4G19000,AT4G19003,AT4G19006,AT4G19010,AT4G19030,AT4G19035,AT4G19038,AT4G18830,AT4G18840,AT4G18930,AT4G18960,AT4G18990,AT4G19020(CMT2),AT4G18890 |
| chr3:15869093 | ARNOLDY1-DNA/MuDR | AT3G44060,AT3G44070,AT3G44080,AT3G44090,AT3G44100,AT3G44110,AT3G44115,AT3G44120,AT3G44130 |
| chr3:22903228 | ATCOPIA31-LTR/Copia | AT3G61670,AT3G61680,AT3G61690,AT3G61700,AT3G61720,AT3G61723,AT3G61730,AT3G61750,AT3G61760,AT3G61770,AT3G61780,AT3G61800,AT3G61810,AT3G61820,AT3G61826,AT3G61829,AT3G61840,AT3G61850,AT3G61860,AT3G61870,AT3G61880,AT3G61890,AT3G61898,AT3G61900,AT3G61910,AT3G61920,AT3G61930,AT3G61950,AT3G61960,AT3G61962,AT3G61710,AT3G61740,AT3G61790,AT3G61830,AT3G61940 |
| chr3:11117812 | ATCOPIA27-LTR/Copia | AT3G29034,AT3G29035,AT3G29040,AT3G29050,AT3G29060,AT3G29070,AT3G29075,AT3G29080,AT3G29090,AT3G29100,AT3G29110,AT3G29130,AT3G29140,AT3G29152,AT3G29160,AT3G29170,AT3G29173,AT3G29180,AT3G29185,AT3G29190,AT3G29200,AT3G29240,AT3G29250,AT3G29230 |
| chr3:22347547 | ATCOPIA22-LTR/Copia | AT3G60400,AT3G60410,AT3G60415,AT3G60420,AT3G60440,AT3G60450,AT3G60460,AT3G60470,AT3G60490,AT3G60500,AT3G60510,AT3G60520,AT3G60480 |
| chr1:6985756 | ATLANTYS2-LTR/Gypsy | AT1G20100,AT1G20110,AT1G20120,AT1G20132,AT1G20135,AT1G20140,AT1G20150,AT1G20160,AT1G20180,AT1G20190,AT1G20200,AT1G20130 |

| | | |
|---|---|---|
| chr2:17291660 | ATDNAI27T9C-DNA/MuDR | AT2G40700,AT2G40710,AT2G40711,AT2G40715,AT2G40720,AT2G40740,AT2G40750,AT2G40745,AT2G40760,AT2G40765,AT2G40770,AT2G40780,AT2G40800,AT2G40810,AT2G40815,AT2G40830,AT2G40840,AT2G40850,AT2G40880,AT2G40890,AT2G40900,AT2G40910,AT2G40920,AT2G40925,AT2G40930,AT2G40935,AT2G40940,AT2G40955,AT2G40960,AT2G40970,AT2G40980,AT2G40990,AT2G40995,AT2G41997,AT2G41000,AT2G41010,AT2G41020,AT2G41040,AT2G41050,AT2G41060,AT2G41070,AT2G41080,AT2G41082,AT2G41090,AT2G41105,AT2G41110,AT2G41120,AT2G41130,AT2G41140,AT2G41160,AT2G41170,AT2G41180,AT2G41200,AT2G41210,AT2G41225,AT2G41230,AT2G41231,AT2G41240,AT2G41250,AT2G41260,AT2G41280,AT2G41290,AT2G41300,AT2G41310,AT2G41330,AT2G41340,AT2G41342,AT2G41350,AT2G41355,AT2G41370,AT2G41375,AT2G41390,AT2G41400,AT2G41410,AT2G41415,AT2G41417,AT2G41420,AT2G41430,AT2G41440,AT2G41445,AT2G41450,AT2G41460,AT2G41470,AT2G41473,AT2G41475,AT2G41480,AT2G41490,AT2G41500,AT2G41505,AT2G41510,AT2G41515,AT2G41530,AT2G41540,AT2G41550,AT2G41590,AT2G41600,AT2G41610,AT2G41630,AT2G41640,AT2G41650,AT2G41660,AT2G41670,AT2G41680,AT2G41690,AT2G41710,AT2G41720,AT2G41730,AT2G41750,AT2G41760,AT2G41770,AT2G41780,AT2G41800,AT2G41810,AT2G41830,AT2G41835,AT2G41840,AT2G41850,AT2G41860,AT2G41870,AT2G41880,AT2G41890,AT2G41900,AT2G41905,AT2G41910,AT2G41920,AT2G41930,AT2G41940,AT2G41945,AT2G41950,AT2G41970,AT2G41980,AT2G41990,AT2G42000,AT2G42005,AT2G42010,AT2G42030,AT2G42040,AT2G42060,AT2G42070,AT2G42065,AT2G42080,AT2G42090,AT2G42100,AT2G42110,AT2G42130,AT2G42140,AT2G42150,AT2G42160,AT2G42180,AT2G42190,AT2G42200,AT2G42210,AT2G42220,AT2G42240,AT2G42250,AT2G42260,AT2G42270,AT2G40730,AT2G40790,AT2G40860,AT2G40950,AT2G41100,AT2G41150,AT2G41220,AT2G41360,AT2G41380,AT2G41451,AT2G41520,AT2G41560,AT2G41620,AT2G41700,AT2G41705,AT2G41740,AT2G41820,AT2G41960,AT2G42170,AT2G42230,AT2G42280,AT2G40820,AT2G41190,AT2G41790,AT2G42120 |
| chr5:10298842 | ATCOPIA92-LTR/Copia | AT5G28290,AT5G28295,AT5G28300,AT5G28310,AT5G28340,AT5G28345,AT5G28370,AT5G28380,AT5G28390,AT5G28400,AT5G28410,AT5G28420,AT5G28442,AT5G28450,AT5G28460,AT5G28463,AT5G28462,AT5G28320,AT5G28350 |

| | | |
|---|---|---|
| chr3:6774161 | ATCOPIA63-LTR/Copia | AT3G19500,AT3G19508,AT3G19515,AT3G19520,AT3G19530,AT3G19550,AT3G19552,AT3G19510,AT3G19540 |
| chr4:7990444 | ATREP5-RC/Helitron | AT4G13730,AT4G13760,AT4G13770,AT4G13790,AT4G13800,AT4G13810,AT4G13750,AT4G13780 |
| chr4:11341911 | HELITRONY1B-RC/Helitron | AT4G21250,AT4G21260,AT4G21270,AT4G21280,AT4G21310,AT4G21320,AT4G21323,AT4G21326,AT4G21330,AT4G21350,AT4G21300,AT4G21340 |
| chr1:20175651 | ATHPOGON3-DNA/Pogo | AT1G54000,AT1G54010,AT1G54020,AT1G54040,AT1G54050,AT1G54060,AT1G54070,AT1G54090,AT1G54095,AT1G53990,AT1G54030,AT1G54080 |
| chr5:9263425 | ATHATN1-DNA/HAT | AT5G26310,AT5G26330,AT5G26340,AT5G26570,AT5G26594,AT5G26667,AT5G26673,AT5G26320,AT5G26360 |
| chr5:4957575 | ATHILA6B-LTR/Gypsy | AT5G15220,AT5G15230,AT5G15250,AT5G15260,AT5G15265,AT5G15270,AT5G15290,AT5G15300,AT5G15310,AT5G15320,AT5G15240,AT5G15280,AT5G15330 |
| chr3:7794973 | RathE2_cons-RathE2_cons | AT3G22085,AT3G22090,AT3G22100,AT3G22104,AT3G22110,AT3G22120,AT3G22125,AT3G22142 |
| chr1:16317933 | ATHAT1-DNA/HAT | AT1G43245,AT1G43310,AT1G43320,AT1G43330,AT1G43415,AT1G43560,AT1G43260 |
| chr3:15677944 | ATHATN6-DNA/HAT | AT3G43800,AT3G43833,AT3G43837,AT3G43810 |
| chr1:23535159 | ATENSPM7-DNA/En-Spm | AT1G63010,AT1G63020(NRPD1),AT1G63030,AT1G63050,AT1G63060,AT1G63070,AT1G63080,AT1G63090,AT1G63100,AT1G63105,AT1G63110,AT1G63120,AT1G63140,AT1G63150,AT1G63160,AT1G63170,AT1G63190,AT1G63200,AT1G63205,AT1G63206,AT1G63220,AT1G63230,AT1G63240,AT1G63245,AT1G63250,AT1G63260,AT1G63270,AT1G63280,AT1G63290,AT1G63295,AT1G63300,AT1G63310,AT1G63320,AT1G63330,AT1G63340;Note=Flavin-containing monooxygenase family protein,AT1G63360,AT1G63370,AT1G63380,AT1G63390,AT1G63400,AT1G63420,AT1G63430,AT1G63440,AT1G63450,AT1G63460,AT1G63470,AT1G63480,AT1G63490,AT1G63500,AT1G63522,AT1G63530,AT1G63535,AT1G63540,AT1G63550,AT1G63055,AT1G63057,AT1G63130,AT1G63180,AT1G63210,AT1G63350,AT1G63410,AT1G63520 |
| chr5:15191802 | RP1_AT-DNA | AT5G37950,AT5G37960,AT5G37970,AT5G37980,AT5G37990,AT5G38000,AT5G38010,AT5G38020,AT5G38040,AT5G38060,AT5G3 |

| | | |
|---|---|---|
| | | 8070,AT5G38080,AT5G38090,AT5G38100,AT5G38110,AT5G38120,AT5G38130,AT5G38150,AT5G38160,AT5G38170,AT5G38180,AT5G38030,AT5G38050,AT5G38140,AT5G38190 |
| chr4:6382376 | ATCOPIA31-LTR/Copia | AT4G10120,AT4G10130,AT4G10150,AT4G10160,AT4G10170,AT4G10180,AT4G10190,AT4G10200,AT4G10210,AT4G10220,AT4G10230,AT4G10240,AT4G10250,AT4G10260,AT4G10270,AT4G10280,AT4G10290,AT4G10300,AT4G10305,AT4G10310,AT4G10320,AT4G10330,AT4G10340,AT4G10350,AT4G10360,AT4G10370,AT4G10140,AT4G10265 |
| chr5:15189300 | ATDNA12T3_2-DNA | AT5G38040,AT5G38060,AT5G38070,AT5G38080,AT5G38090,AT5G38100,AT5G38110,AT5G38120,AT5G38130,AT5G38150,AT5G38160,AT5G38170,AT5G38180,AT5G38030,AT5G38050,AT5G38140,AT5G38190 |
| chr3:10494234 | ATCOPIA50-LTR/Copia | AT3G28170,AT3G28180,AT3G28155 |
| chr3:10494234 | ATCOPIA74-LTR/Copia | AT3G27999,AT3G28007,AT3G28020,AT3G28030,AT3G28050,AT3G28060,AT3G28070,AT3G28100,AT3G28120,AT3G28130,AT3G28140,AT3G28150,AT3G28170,AT3G28180,AT3G28190,AT3G28193,AT3G28210,AT3G28216,AT3G28220,AT3G28223,AT3G28230,AT3G28243,AT3G28250,AT3G28270,AT3G28280,AT3G28040,AT3G28080,AT3G28200,AT3G28155 |
| chr1:19484366 | ATCOPIA59-LTR/Copia | AT1G51190,AT1G51200,AT1G51220,AT1G51230,AT1G51240,AT1G51250,AT1G51270,AT1G51290,AT1G51300,AT1G51310,AT1G51320,AT1G51330,AT1G51340,AT1G51355,AT1G51360,AT1G51370,AT1G51380,AT1G51390,AT1G51400,AT1G51402,AT1G51405,AT1G51410,AT1G51420,AT1G51430,AT1G51450,AT1G51460,AT1G51470,AT1G51480,AT1G51490,AT1G51500,AT1G51520,AT1G51530,AT1G51538,AT1G51540,AT1G51550,AT1G51570,AT1G51580,AT1G51590,AT1G51600,AT1G51620,AT1G51630,AT1G51640,AT1G51660,AT1G51670,AT1G51680,AT1G51700,AT1G51710,AT1G51730,AT1G51740,AT1G51745(PWWP2),AT1G51760,AT1G51780,AT1G51790,AT1G51800,AT1G51805,AT1G51810,AT1G51820,AT1G51823,AT1G51830,AT1G51840,AT1G51850,AT1G51860,AT1G51870,AT1G51880,AT1G51890,AT1G51900,AT1G51910,AT1G51913,AT1G51915,AT1G51920,AT1G51930,AT1G51940,AT1G51950,AT1G51960,AT1G51965,AT1G51970,AT1G51980,AT1G51990,AT1G52030,AT1G52040,AT1G52050,AT1G52060,AT1G52080,AT1G52100,AT1G52110,AT1G52120,AT1G52130,AT1G52140,AT1G52150,AT1G52155,AT1G52160,AT1G52180,AT1G52190,AT1G52191,AT1G52200,AT1G52220,AT1G52230,AT1G52240,AT1G52245,A |

| | | |
|---|---|---|
| | | T1G52260,AT1G52280,AT1G52290,AT1G52300,AT1G52315,AT1G52320,AT1G52325,AT1G52330,AT1G52340,AT1G52342,AT1G52343,AT1G52360,AT1G52370,AT1G52380,AT1G52390,AT1G52400,AT1G52410,AT1G52415,AT1G52430,AT1G52440,AT1G52460,AT1G52470,AT1G52490,AT1G52495,AT1G52520,AT1G52530,AT1G52540,AT1G52550,AT1G52560,AT1G52565,AT1G52570,AT1G52580,AT1G52590,AT1G52600,AT1G52603,AT1G52618,AT1G52620,AT1G52630,AT1G52640,AT1G52660,AT1G52670,AT1G52680,AT1G52690,AT1G52695,AT1G52700,AT1G52710,AT1G52720,AT1G52740(HTA9),AT1G52750,AT1G52760,AT1G52770,AT1G52790,AT1G52800,AT1G52810,AT1G52820,AT1G52825,AT1G52827,AT1G52830,AT1G52857,AT1G52855,AT1G52870,AT1G52880,AT1G52890,AT1G52900,AT1G52905,AT1G52910,AT1G52930,AT1G52940,AT1G52970,AT1G52980,AT1G53000,AT1G53010,AT1G53023,AT1G53030,AT1G53035,AT1G53040,AT1G53060,AT1G53070,AT1G53080,AT1G53090,AT1G53100,AT1G53120,AT1G53130,AT1G53140,AT1G53160,AT1G53163,AT1G53165,AT1G53170,AT1G53180,AT1G53190,AT1G53210,AT1G53230,AT1G53240,AT1G53260,AT1G53265,AT1G53270,AT1G53280,AT1G53282,AT1G53285,AT1G53290,AT1G53300,AT1G53310,AT1G53320,AT1G53325,AT1G53330,AT1G53340,AT1G53345,AT1G53360,AT1G53366,AT1G53370,AT1G53380,AT1G53400,AT1G53420,AT1G53440,AT1G53450,AT1G53470,AT1G53480,AT1G53490,AT1G53500,AT1G53520,AT1G53530,AT1G53540,AT1G53541,AT1G53542,AT1G53550,AT1G53560,AT1G51210,AT1G51350,AT1G51440,AT1G51510,AT1G51560,AT1G51650,AT1G51690,AT1G51720,AT1G51770,AT1G52000,AT1G52070,AT1G52270,AT1G52310,AT1G52450,AT1G52500,AT1G52510,AT1G52650,AT1G52730,AT1G52920,AT1G52950,AT1G52990,AT1G53025,AT1G53050,AT1G53200,AT1G53250,AT1G53350,AT1G53430,AT1G53460,AT1G53510,AT1G51260,AT1G51610,AT1G52420,AT1G52780,AT1G53110,AT1G53390 |
| chr4:7956078 | ATLINE1_5-LINE/L1 | AT4G13500,AT4G13510,AT4G13530,AT4G13540,AT4G13550,AT4G13570,AT4G13572,AT4G13575,AT4G13580,AT4G13590,AT4G13600,AT4G13610,AT4G13615,AT4G13620,AT4G13630,AT4G13640,AT4G13650,AT4G13670,AT4G13680,AT4G13690,AT4G13700,AT4G13720,AT4G13730,AT4G13560,AT4G13577,AT4G13660,AT4G13710,AT4G13520 |
| chr4:14244202 | ATCOPIA46-LTR/Copia | AT4G28540,AT4G28550,AT4G28556,AT4G28560,AT4G28570,AT4G28580,AT4G28590,AT4G28600,AT4G28610,AT4G28620,AT4G28640,AT4G28650,AT4G28660,AT4G28670,AT4G28690,AT4G28700,AT4G28703,AT4G28720,AT4G28730,AT4G28740,AT4G28750,A |

| | | |
|---|---|---|
| | | T4G28755,AT4G28760,AT4G28770,AT4G28775,AT4G28780,AT4G28790,AT4G28800,AT4G28811,AT4G28815,AT4G28820,AT4G28830,AT4G28840,AT4G28850,AT4G28860,AT4G28870,AT4G28890,AT4G28910,AT4G28920,AT4G28930,AT4G28940,AT4G28950,AT4G28980,AT4G28990,AT4G29000,AT4G29010,AT4G29020,AT4G29030,AT4G29033,AT4G29035,AT4G29037,AT4G29040(RPT2a),AT4G29050,AT4G29070,AT4G29080,AT4G28630,AT4G28706,AT4G28710,AT4G28880,AT4G29060,AT4G28680 |
| chr2:2148268 | ATGP1-LTR/Gypsy | AT2G05720,AT2G05752,AT2G05755,AT2G05753,AT2G05710 |
| chr5:7670238 | VANDAL18NB-DNA/MuDR | AT5G22900,AT5G22910,AT5G22920,AT5G22940,AT5G22950,AT5G22970,AT5G22980,AT5G22990,AT5G23000,AT5G23010,AT5G23020,AT5G23030,AT5G23035,AT5G23040,AT5G23060,AT5G23070,AT5G23080,AT5G23100,AT5G23115,AT5G23120,AT5G23140,AT5G23150,AT5G23160,AT5G23180,AT5G23190,AT5G23200,AT5G23210,AT5G23212,AT5G23220,AT5G22930,AT5G22960,AT5G23090,AT5G23110,AT5G23130,AT5G23170,AT5G23050 |
| chr2:4943212 | ATHATN1-DNA/HAT | AT2G12400,AT2G12405 |
| chr5:26121244 | ATCOPIA29-LTR/Copia | AT5G65310,AT5G65320,AT5G65330,AT5G65340,AT5G65350,AT5G65360,AT5G65370,AT5G65380,AT5G65390,AT5G65410,AT5G65400 |
| chr2:9835396 | ARNOLD2-DNA/MuDR | AT2G23067,AT2G23070,AT2G23080,AT2G23093,AT2G23096,AT2G23100,AT2G23110,AT2G23120,AT2G23118,AT2G23130,AT2G23142,AT2G23140,AT2G23090 |
| chr5:26428195 | AT9MU1-DNA/MuDR | AT5G64620,AT5G64630(FAS2),AT5G64650,AT5G64660,AT5G64667,AT5G64680,AT5G64687,AT5G64690,AT5G64700,AT5G64720,AT5G64730,AT5G64750,AT5G64760,AT5G64770,AT5G64780,AT5G64800,AT5G64810,AT5G64813,AT5G64816,AT5G64820,AT5G64830,AT5G64850,AT5G64860,AT5G64870,AT5G64880,AT5G64890,AT5G64900,AT5G64905,AT5G64910,AT5G64920,AT5G64930,AT5G64950,AT5G64960,AT5G64970,AT5G64980,AT5G64990,AT5G65000,AT5G65005,AT5G65010,AT5G65020,AT5G65030,AT5G65040,AT5G65060,AT5G65070,AT5G65090,AT5G65100,AT5G65110,AT5G65120,AT5G65140,AT5G65158,AT5G65160,AT5G65165,AT5G65170,AT5G65180,AT5G65200,AT5G65205,AT5G65207,AT5G65210,AT5G65220,AT5G65230,AT5G65240,AT5G65250,AT5G65260,AT5G65274,AT5G65280,AT5G65290,AT5G65300,AT5G65310,AT5G65320,AT5G65330,AT5G65340,AT5G65350,AT5G65360,A |

| | | |
|---|---|---|
| | | T5G65370,AT5G65380,AT5G65390,AT5G65410,AT5G65420,AT5G65440,AT5G65450,AT5G65470,AT5G65480,AT5G65490,AT5G65495,AT5G65510,AT5G65520,AT5G65530,AT5G65550,AT5G65560,AT5G65570,AT5G65580,AT5G65590,AT5G65600,AT5G65609,AT5G65610,AT5G65613,AT5G65620,AT5G65630,AT5G65640,AT5G65650,AT5G65660,AT5G65670,AT5G65690,AT5G65687,AT5G65685,AT5G65700,AT5G65710,AT5G65730,AT5G65740,AT5G65750,AT5G65760,AT5G65790,AT5G65800,AT5G65810,AT5G65820,AT5G65830,AT5G65850,AT5G65860,AT5G65870,AT5G65880,AT5G65890,AT5G65900,AT5G65920,AT5G65925,AT5G65930,AT5G65950,AT5G65960,AT5G65970,AT5G65990,AT5G66000,AT5G66005,AT5G66010,AT5G66020,AT5G66030,AT5G66050,AT5G66052,AT5G66053,AT5G66055,AT5G66060,AT5G66070,AT5G66080,AT5G66090,AT5G66110,AT5G66120,AT5G66130,AT5G66140,AT5G66160,AT5G66170,AT5G66180,AT5G66190,AT5G66200,AT5G66210,AT5G66230,AT5G66220,AT5G66240,AT5G66250,AT5G66260,AT5G66270,AT5G66290,AT5G66300,AT5G66310,AT5G66320,AT5G66330,AT5G66335,AT5G66340,AT5G66350,AT5G66360,AT5G66370,AT5G66380,AT5G66390,AT5G66400,AT5G66410,AT5G66420,AT5G66440,AT5G66450,AT5G66460,AT5G66480,AT5G66490,AT5G66500,AT5G66510,AT5G66530,AT5G66520,AT5G66550,AT5G66560,AT5G66570,AT5G66580,AT5G66600,AT5G66595,AT5G66607,AT5G64640,AT5G64670,AT5G64710,AT5G64740,AT5G64790,AT5G64840,AT5G65050,AT5G65080,AT5G65130,AT5G65225,AT5G65400,AT5G65430,AT5G65460,AT5G65500,AT5G65540,AT5G65683,AT5G65720,AT5G65770,AT5G65780,AT5G65910,AT5G65940,AT5G65980,AT5G66040,AT5G66100,AT5G66150,AT5G66430,AT5G66470,AT5G66540,AT5G66590,AT5G64940,AT5G65270,AT5G65840,AT5G66280 |
| chr5:26428195 | AT9NMU1-DNA/MuDR | AT5G64620,AT5G64630(FAS2),AT5G64650,AT5G64660,AT5G64667,AT5G64680,AT5G64687,AT5G64690,AT5G64700,AT5G64720,AT5G64730,AT5G64750,AT5G64760,AT5G64770,AT5G64780,AT5G64800,AT5G64810,AT5G64813,AT5G64816,AT5G64820,AT5G64830,AT5G64850,AT5G64860,AT5G64870,AT5G64880,AT5G64890,AT5G64900,AT5G64905,AT5G64910,AT5G64920,AT5G64930,AT5G64950,AT5G64960,AT5G64970,AT5G64980,AT5G64990,AT5G65000,AT5G65005,AT5G65010,AT5G65020,AT5G65030,AT5G65040,AT5G65060,AT5G65070,AT5G65090,AT5G65100,AT5G65110,AT5G65120,AT5G65140,AT5G65158,AT5G65160,AT5G65165,AT5G65170,AT5G65180,AT5G65200,AT5G65205,AT5G65207,AT5G65210,AT5G65220,AT5G65230,AT5G65240,AT5G65250,AT5G6 |

| | | |
|---|---|---|
| | | 5260,AT5G65274,AT5G65280,AT5G65290,AT5G65300,AT5G65310,AT5G65320,AT5G65330,AT5G65340,AT5G65350,AT5G65360,AT5G65370,AT5G65380,AT5G65390,AT5G65410,AT5G65420,AT5G65440,AT5G65450,AT5G65470,AT5G65480,AT5G65490,AT5G65495,AT5G65510,AT5G65520,AT5G65530,AT5G65550,AT5G65560,AT5G65570,AT5G65580,AT5G65590,AT5G65600,AT5G65609,AT5G65610,AT5G65613,AT5G65620,AT5G65630,AT5G65640,AT5G65650,AT5G65660,AT5G65670,AT5G65690,AT5G65687,AT5G65685,AT5G65700,AT5G65710,AT5G65730,AT5G65740,AT5G65750,AT5G65760,AT5G65790,AT5G65800,AT5G65810,AT5G65820,AT5G65830,AT5G65850,AT5G65860,AT5G65870,AT5G65880,AT5G65890,AT5G65900,AT5G65920,AT5G65925,AT5G65930,AT5G65950,AT5G65960,AT5G65970,AT5G65990,AT5G66000,AT5G66005,AT5G66010,AT5G66020,AT5G66030,AT5G66050,AT5G66052,AT5G66053,AT5G66055,AT5G66060,AT5G66070,AT5G66080,AT5G66090,AT5G66110,AT5G66120,AT5G66130,AT5G66140,AT5G66160,AT5G66170,AT5G66180,AT5G66190,AT5G66200,AT5G66210,AT5G66230,AT5G66220,AT5G66240,AT5G66250,AT5G66260,AT5G66270,AT5G66290,AT5G66300,AT5G66310,AT5G66320,AT5G66330,AT5G66335,AT5G66340,AT5G66350,AT5G66360,AT5G66370,AT5G66380,AT5G66390,AT5G66400,AT5G66410,AT5G66420,AT5G66440,AT5G66450,AT5G66460,AT5G66480,AT5G66490,AT5G66500,AT5G66510,AT5G66530,AT5G66520,AT5G66550,AT5G66560,AT5G66570,AT5G66580,AT5G66600,AT5G66595,AT5G66607,AT5G64640,AT5G64670,AT5G64710,AT5G64740,AT5G64790,AT5G64840,AT5G65050,AT5G65080,AT5G65130,AT5G65225,AT5G65400,AT5G65430,AT5G65460,AT5G65500,AT5G65540,AT5G65683,AT5G65720,AT5G65770,AT5G65780,AT5G65910,AT5G65940,AT5G65980,AT5G66040,AT5G66100,AT5G66150,AT5G66430,AT5G66470,AT5G66540,AT5G66590,AT5G64940,AT5G65270,AT5G65840,AT5G66280 |
| chr3:15775672 | ATCOPIA62-LTR/Copia | AT3G43910,AT3G43920(DCL3),AT3G43940,AT3G43950,AT3G43960,AT3G43980,AT3G43990,AT3G44006,AT3G44010,AT3G44020,AT3G44050,AT3G44060,AT3G44070,AT3G44080,AT3G44090,AT3G43930,AT3G43970 |
| chr3:15102607 | ATCOPIA11-LTR/Copia | AT3G43110,AT3G43120 |
| chr5:4967255 | ATHATN4-DNA/HAT | AT5G15220,AT5G15230,AT5G15250,AT5G15260,AT5G15265,AT5G15270,AT5G15290,AT5G15300,AT5G15310,AT5G15320,AT5G15240,AT5G15280,AT5G15330 |

| | | |
|---|---|---|
| chr2:7454593 | ATCOPIA71-LTR/Copia | AT2G17060,AT2G17070,AT2G17080,AT2G17090,AT2G17110,AT2G17120,AT2G17130,AT2G17140,AT2G17160,AT2G17170,AT2G17180,AT2G17190,AT2G17200,AT2G17210,AT2G17150 |
| chr5:22246184 | HELITRONY1E-RC/Helitron | AT5G54585,AT5G54590,AT5G54610,AT5G54620,AT5G54630,AT5G54640,AT5G54660,AT5G54670,AT5G54680,AT5G54690,AT5G54700,AT5G54710,AT5G54720,AT5G54730,AT5G54740,AT5G54745,AT5G54750,AT5G54760,AT5G54770,AT5G54790,AT5G54800,AT5G54820,AT5G54830,AT5G54840,AT5G54855,AT5G54860,AT5G54870,AT5G54880,AT5G54890,AT5G54900,AT5G54920,AT5G54930,AT5G54940,AT5G54950,AT5G54960,AT5G54970,AT5G54980,AT5G54990,AT5G55000,AT5G55010,AT5G55020,AT5G55050,AT5G55060,AT5G55080,AT5G55090,AT5G55110,AT5G55120,AT5G55125,AT5G55130,AT5G54600,AT5G54650,AT5G54780,AT5G54810,AT5G54910,AT5G55040,AT5G55070,AT5G55100,AT5G54850 |
| chr5:14142812 | ATCOPIA91-LTR/Copia | AT5G35920,AT5G35926,AT5G35927,AT5G35940,AT5G35945,AT5G35960,AT5G35980,AT5G35995,AT5G36000,AT5G36001,AT5G36080,AT5G36100,AT5G36110,AT5G36120,AT5G36130,AT5G36140,AT5G36160,AT5G36180,AT5G36190,AT5G36200,AT5G36210,AT5G36225,AT5G36228,AT5G36230,AT5G36240,AT5G36250,AT5G35930,AT5G35950,AT5G35970,AT5G36150,AT5G36170,AT5G36220,AT5G36260 |
| chr1:16273576 | ATCOPIA72-LTR/Copia | AT1G43160,AT1G43170,AT1G43190,AT1G43245,AT1G43310,AT1G43320,AT1G43330,AT1G43415,AT1G43560,AT1G43580,AT1G43600,AT1G43171,AT1G43260,AT1G43605 |
| chr5:2442479 | ATCOPIA18-LTR/Copia | AT5G07210,AT5G07220,AT5G07225,AT5G07230,AT5G07240,AT5G07250,AT5G07260,AT5G07270,AT5G07280,AT5G07300,AT5G07310,AT5G07330,AT5G07340,AT5G07350,AT5G07360,AT5G07370,AT5G07380,AT5G07390,AT5G07410,AT5G07420,AT5G07430,AT5G07440,AT5G07460,AT5G07470,AT5G07475,AT5G07480,AT5G07490,AT5G07500,AT5G07510,AT5G07520,AT5G07530,AT5G07540,AT5G07545,AT5G07550,AT5G07560,AT5G07571,AT5G07572,AT5G07580,AT5G07590,AT5G07600,AT5G07610,AT5G07630,AT5G07640,AT5G07650,AT5G07670,AT5G07680,AT5G07690,AT5G07700,AT5G07710,AT5G07720,AT5G07740,AT5G07760,AT5G07770,AT5G07780,AT5G07790,AT5G07800,AT5G07810,AT5G07820,AT5G07290,AT5G07320,AT5G07400,AT5G07570,AT5G07620,AT5G07660,AT5G07730,AT5G07450 |

| | | |
|---|---|---|
| chr1:22601456 | ATHILA4C-LTR/Gypsy | AT1G61250,AT1G61255,AT1G61270,AT1G61280,AT1G61290,AT1G61310,AT1G61320,AT1G61260,AT1G61300 |
| chr1:24432078 | ATENSPM1-DNA/En-Spm | AT1G65650,AT1G65660,AT1G65670,AT1G65681,AT1G65690,AT1G65700,AT1G65710,AT1G65720,AT1G65735,AT1G65740,AT1G65680,AT1G65730 |
| chr2:13194438 | ATCOPIA44-LTR/Copia | AT2G30970,AT2G30980,AT2G30985,AT2G31005,AT2G31010,AT2G31020,AT2G31018,AT2G31030,AT2G31040,AT2G31035,AT2G30990 |
| chr3:14167801 | ATLANTYS1-LTR/Gypsy | AT3G32920,AT3G32930,AT3G32940,AT3G32960,AT3G32980,AT3G33187,AT3G33293,AT3G33393,AT3G33494,AT3G33528,AT3G33520(ARP6),AT3G41762,AT3G42050,AT3G42075,AT3G42130,AT3G42140,AT3G42148,AT3G42150,AT3G42153,AT3G42155,AT3G42160,AT3G42170,AT3G42180,AT3G42310,AT3G42390,AT3G42473,AT3G42550,AT3G42560,AT3G42565,AT3G42570,AT3G42628,AT3G42630,AT3G42640,AT3G42660,AT3G42670(CLSY1),AT3G33530,AT3G42060 |
| chr5:7896154 | ATCOPIA32-LTR/Copia | AT5G23405,AT5G23411,AT5G23420,AT5G23440,AT5G23450,AT5G23400,AT5G23430 |
| chr2:12255099 | HELITRON2-RC/Helitron | AT2G28570,AT2G28580,AT2G28590,AT2G28600,AT2G28610,AT2G28620,AT2G28560,AT2G28605 |
| chr3:17117971 | LIMPET1-DNA/MuDR | AT3G46490,AT3G46510,AT3G46520,AT3G46530,AT3G46480,AT3G46500 |
| chr4:11241351 | ATCOPIA11-LTR/Copia | AT4G21010,AT4G21020,AT4G21030,AT4G21040,AT4G21050,AT4G21060,AT4G21063,AT4G21065,AT4G21070,AT4G21080,AT4G21090,AT4G21100,AT4G21105,AT4G21110 |
| chr1:21736735 | ARNOLDY2-DNA/MuDR | AT1G58450,AT1G58460,AT1G58470,AT1G58525,AT1G58520,AT1G58602 |
| chr5:16048359 | ATCOPIA18A-LTR/Copia | AT5G40030,AT5G40040,AT5G40050,AT5G40060,AT5G40070,AT5G40090,AT5G40100,AT5G40120,AT5G40140,AT5G40150,AT5G40153,AT5G40155,AT5G40160,AT5G40080 |
| chr1:10120008 | ATCOPIA71-LTR/Copia | AT1G28765,AT1G28960,AT1G29000,AT1G29005,AT1G29010,AT1G29020,AT1G29030,AT1G29040,AT1G29041,AT1G29050,AT1G29060,AT1G29025 |
| chr3:21416229 | SADHU-Unassigned | AT3G57770,AT3G57780,AT3G57785,AT3G57787,AT3G57790,AT3G57800,AT3G57810,AT3G57830,AT3G57840,AT3G57850,AT3G57860,AT3G57880,AT3G57870 |

| | | |
|---|---|---|
| chr5:25473574 | VANDAL10-DNA/MuDR | AT5G63470,AT5G63480,AT5G63490,AT5G63500,AT5G63510,AT5G63530,AT5G63550,AT5G63560,AT5G63570,AT5G63580,AT5G63590,AT5G63595,AT5G63610,AT5G63620,AT5G63625,AT5G63630,AT5G63640,AT5G63650,AT5G63660,AT5G63670,AT5G63540,AT5G63600,AT5G63520 |
| chr4:1559762 | VANDAL5-DNA/MuDR | AT4G03470,AT4G03480,AT4G03490,AT4G03495,AT4G03505,AT4G03510,AT4G03520,AT4G03540,AT4G03550,AT4G03560,AT4G03565,AT4G03566,AT4G03570,AT4G03580,AT4G03590,AT4G03600,AT4G03620,AT4G03625,AT4G03630,AT4G03635,AT4G03500,AT4G03610 |
| chr2:9277465 | ATENSPM7-DNA/En-Spm | AT2G21580,AT2G21595,AT2G21600,AT2G21610,AT2G21620,AT2G21630,AT2G21640,AT2G21650,AT2G21655,AT2G21660,AT2G21680,AT2G21690,AT2G21720,AT2G21725,AT2G21727,AT2G21730,AT2G21740,AT2G21750,AT2G21780,AT2G21790,AT2G21800,AT2G21590,AT2G21710,AT2G21770 |
| chr5:20204392 | ATCOPIA82-LTR/Copia | AT5G49690,AT5G49700,AT5G49710,AT5G49720,AT5G49740,AT5G49750,AT5G49770,AT5G49780,AT5G49800,AT5G49810,AT5G49820,AT5G49840,AT5G49850,AT5G49860,AT5G49870,AT5G49730,AT5G49760,AT5G49830,AT5G49680 |
| chr2:1557162 | VANDAL10-DNA/MuDR | AT2G04300,AT2G04305,AT2G04340,AT2G04350,AT2G04360,AT2G04378,AT2G04380,AT2G04390,AT2G04395,AT2G04400,AT2G04410,AT2G04420,AT2G04425,AT2G04440,AT2G04450,AT2G04480,AT2G04495,AT2G04500,AT2G04430 |
| chr4:13391455 | DT1-DNA/Mariner | AT4G26020,AT4G26030,AT4G26040,AT4G26050,AT4G26055,AT4G26060,AT4G26070,AT4G26090,AT4G26100,AT4G26110,AT4G26130,AT4G26140,AT4G26145,AT4G26160,AT4G26170(ET1),AT4G26180,AT4G26190,AT4G26200,AT4G26210,AT4G26220,AT4G26230,AT4G26250,AT4G26260,AT4G26280,AT4G26288,AT4G26290,AT4G26300,AT4G26310,AT4G26320,AT4G26330,AT4G26340,AT4G26350,AT4G26370,AT4G26380,AT4G26390,AT4G26400,AT4G26410,AT4G26415,AT4G26420,AT4G26430,AT4G26440,AT4G26450,AT4G26455,AT4G26466,AT4G26460,AT4G26470,AT4G26480,AT4G26483,AT4G26485,AT4G26490,AT4G26500,AT4G26520,AT4G26530,AT4G26540,AT4G26550,AT4G26555,AT4G26080,AT4G26150,AT4G26240,AT4G26270,AT4G26510,AT4G26120 |
| chr3:15392159 | ATLINE1_6-LINE/L1 | AT3G43420,AT3G43430,AT3G43432,AT3G43470,AT3G43480,AT3G43490,AT3G43500,AT3G43505,AT3G43520,AT3G43540,AT3G43550,AT3G43570,AT3G43572,AT3G43574,AT3G43580,AT3G43583,AT3G43590,AT3G43610,AT3G43440,AT3G43503,AT3G43600 |

Table S7. Summary of significant peaks in TE family copy number GWAS analysis.

| Chromosome | Lead SNP | P value | TE family | Peak type |
|---|---|---|---|---|
| chr1 | 10607332 | 2.60E-16 | ATCOPIA83-LTR/Copia | same family |
| chr1 | 11011882 | 1.44E-65 | ATCOPIA97-LTR/Copia | same family |
| chr1 | 11156643 | 3.16E-14 | ATCOPIA52-LTR/Copia | same family |
| chr1 | 11156643 | 8.98E-38 | ATCOPIA51-LTR/Copia | same family |
| chr1 | 11156879 | 7.45E-12 | ATCOPIA25-LTR/Copia | same family |
| chr1 | 11157043 | 1.53E-10 | ATCOPIA79-LTR/Copia | same family |
| chr1 | 11177724 | 3.86E-58 | ATCOPIA50-LTR/Copia | same family |
| chr1 | 12015691 | 4.39E-16 | ATENSPM1A-DNA/En-Spm | same family |
| chr1 | 12143760 | 4.62E-34 | ATREP14-RC/Helitron | same family |
| chr1 | 12652897 | 4.57E-45 | ATHATN3A-DNA/HAT | same family |
| chr1 | 12893547 | 4.59E-16 | ATHILA7-LTR/Gypsy | same family |
| chr1 | 13097162 | 3.06E-22 | ATHPOGON2-DNA/Pogo | same family |
| chr1 | 13109197 | 9.75E-15 | ATCOPIA33-LTR/Copia | others |
| chr1 | 13276232 | 9.03E-18 | VANDAL9-DNA/MuDR | same family |
| chr1 | 13323600 | 5.74E-27 | ATCOPIA84-LTR/Copia | same family |
| chr1 | 13434878 | 1.03E-80 | ATMU11-DNA/MuDR | same family |
| chr1 | 13633201 | 2.53E-38 | ATCOPIA89-LTR/Copia | same family |
| chr1 | 13633429 | 1.36E-09 | ATHILA4B_LTR-LTR/Gypsy | same family |
| chr1 | 13717668 | 1.14E-25 | ATHPOGO-DNA/Pogo | others |
| chr1 | 13808790 | 1.22E-26 | ATGP9LTR-LTR/Gypsy | same family |
| chr1 | 14212288 | 3.27E-12 | ATCOPIA57-LTR/Copia | others |
| chr1 | 14243988 | 1.60E-13 | ATHATN1-DNA/HAT | same family |
| chr1 | 14286125 | 1.03E-67 | ATREP18-DNA | same family |
| chr1 | 14334057 | 2.53E-10 | ATHILA7-LTR/Gypsy | same family |
| chr1 | 14446668 | 1.23E-11 | ATGP5-LTR/Gypsy | others |
| chr1 | 15561355 | 2.58E-11 | ATCOPIA57-LTR/Copia | others |
| chr1 | 15590950 | 3.09E-10 | ATHILA7-LTR/Gypsy | others |
| chr1 | 15596008 | 2.76E-13 | ATGP5-LTR/Gypsy | same family |
| chr1 | 15598871 | 7.42E-12 | ATHILA0_I-LTR/Gypsy | same family |
| chr1 | 15601600 | 2.59E-63 | ATREP18-DNA | same family |
| chr1 | 15890614 | 3.45E-31 | ATHATN1-DNA/HAT | same family |
| chr1 | 15923352 | 2.46E-10 | ATHPOGO-DNA/Pogo | others |
| chr1 | 15983870 | 1.64E-13 | ATHAT3-DNA/HAT | others |
| chr1 | 16271132 | 6.36E-32 | ATCOPIA33-LTR/Copia | same family |
| chr1 | 16379185 | 6.40E-26 | ATDNA1T9A-DNA/MuDR | same family |
| chr1 | 16549604 | 2.66E-22 | ATCOPIA93-LTR/Copia | same family |
| chr1 | 16581090 | 1.88E-11 | ATCOPIA81-LTR/Copia | same family |
| chr1 | 16593969 | 3.26E-09 | ATCOPIA44-LTR/Copia | others |
| chr1 | 16639248 | 2.04E-17 | ATCOPIA25-LTR/Copia | same family |
| chr1 | 16790732 | 1.82E-10 | ATCOPIA4-LTR/Copia | same family |
| chr1 | 16891707 | 4.33E-95 | VANDAL7-DNA/MuDR | same family |
| chr1 | 17019880 | 1.59E-54 | VANDALNX1-DNA/MuDR | same family |

| | | | | |
|---|---|---|---|---|
| chr1 | 17029326 | 9.44E-72 | ATCOPIA79-LTR/Copia | same family |
| chr1 | 17187455 | 1.95E-17 | ATTIRX1D-DNA | same family |
| chr1 | 17526477 | 7.80E-98 | ATCOPIA38A-LTR/Copia | same family |
| chr1 | 17555714 | 1.70E-11 | ATHPOGON1-DNA/Pogo | same family |
| chr1 | 17821876 | 4.98E-11 | ATDNA2T9B-DNA/MuDR | same family |
| chr1 | 17827103 | 1.35E-39 | ATHATN3A-DNA/HAT | same family |
| chr1 | 17840214 | 8.59E-16 | ATMU7-DNA/MuDR | same family |
| chr1 | 18297667 | 2.18E-18 | ATRE1-LTR/Copia | same family |
| chr1 | 18519237 | 2.98E-12 | ATHAT1-DNA/HAT | others |
| chr1 | 18550316 | 2.12E-10 | ATTIR16T3A-DNA | same family |
| chr1 | 18550465 | 1.29E-17 | ATCOPIA52-LTR/Copia | others |
| chr1 | 19003520 | 7.64E-22 | ATCOPIA68-LTR/Copia | same family |
| chr1 | 19241179 | 7.91E-13 | ATTIRX1D-DNA | same family |
| chr1 | 20175306 | 3.43E-23 | ATHATN10-DNA/HAT | same family |
| chr1 | 20471994 | 4.23E-18 | ATGP2N-LTR/Gypsy | same family |
| chr1 | 2071605 | 2.14E-11 | ATCOPIA59-LTR/Copia | others |
| chr1 | 21105081 | 5.65E-15 | ATHPOGON1-DNA/Pogo | same family |
| chr1 | 21244304 | 6.05E-55 | ATCOPIA26-LTR/Copia | same family |
| chr1 | 21341250 | 1.47E-210 | ATMUN2-DNA/MuDR | same family |
| chr1 | 2161368 | 6.22E-18 | ATCOPIA58-LTR/Copia | same family |
| chr1 | 22109873 | 1.55E-25 | ATCOPIA5-LTR/Copia | same family |
| chr1 | 22111484 | 7.13E-09 | ATN9_1-DNA/MuDR | others |
| chr1 | 22563850 | 1.37E-09 | ATMU6N1-DNA/MuDR | same family |
| chr1 | 22616313 | 5.48E-156 | ATCOPIA60-LTR/Copia | same family |
| chr1 | 23097954 | 6.24E-17 | ATREP19-DNA | same family |
| chr1 | 23423591 | 1.23E-14 | ATMUN1-DNA/MuDR | same family |
| chr1 | 24319975 | 1.48E-12 | ATCOPIA11-LTR/Copia | same family |
| chr1 | 2543710 | 2.52E-15 | ATHAT1-DNA/HAT | same family |
| chr1 | 26582825 | 7.17E-196 | ATHATN5-DNA/HAT | same family |
| chr1 | 28525544 | 2.71E-28 | ATMU6-DNA/MuDR | same family |
| chr1 | 29474237 | 5.58E-13 | ATMU3-DNA/MuDR | same family |
| chr1 | 3186493 | 3.19E-14 | ATCOPIA12-LTR/Copia | same family |
| chr1 | 3362042 | 2.31E-24 | ATCOPIA78-LTR/Copia | same family |
| chr1 | 3636643 | 6.50E-15 | ATCOPIA53-LTR/Copia | others |
| chr1 | 4208687 | 7.72E-09 | ATMU3N1-DNA/MuDR | others |
| chr1 | 5916747 | 2.41E-12 | ATMU7-DNA/MuDR | same family |
| chr1 | 5918999 | 1.09E-20 | ATMU3-DNA/MuDR | same family |
| chr1 | 6310478 | 1.88E-12 | ATCOPIA58-LTR/Copia | same family |
| chr1 | 8383571 | 1.42E-57 | ATHATN9-DNA/HAT | same family |
| chr1 | 8516338 | 2.86E-16 | ATHILA7-LTR/Gypsy | same family |
| chr1 | 8719142 | 6.00E-19 | ATCOPIA12-LTR/Copia | same family |
| chr1 | 9367779 | 1.54E-16 | ATCOPIA16-LTR/Copia | same family |
| chr1 | 9939725 | 2.44E-21 | HARBINGER-DNA/Harbinger | same family |
| chr2 | 10076422 | 6.69E-14 | ATHAT3-DNA/HAT | others |
| chr2 | 10083642 | 3.92E-11 | VANDAL6-DNA/MuDR | same family |
| chr2 | 10360797 | 1.06E-16 | ATCOPIA74-LTR/Copia | same family |
| chr2 | 10432025 | 2.47E-10 | BOMZH1-DNA/MuDR | same family |

| | | | | |
|---|---|---|---|---|
| chr2 | 10487686 | 1.23E-13 | ATCOPIA13-LTR/Copia | same family |
| chr2 | 10487686 | 1.80E-22 | ATCOPIA12-LTR/Copia | same family |
| chr2 | 10891327 | 1.16E-65 | ATCOPIA39-LTR/Copia | others |
| chr2 | 11238460 | 1.09E-14 | ATCOPIA89-LTR/Copia | same family |
| chr2 | 11400550 | 8.02E-23 | ATHATN7-DNA/HAT | same family |
| chr2 | 12589468 | 5.02E-09 | ATCOPIA63-LTR/Copia | others |
| chr2 | 1359238 | 2.74E-201 | TA1-2-LTR/Copia | same family |
| chr2 | 1380259 | 4.22E-11 | ATN9_1-DNA/MuDR | same family |
| chr2 | 13922516 | 1.21E-19 | ATGP3-LTR/Gypsy | others |
| chr2 | 14407341 | 1.34E-26 | ATENSPM1-DNA/En-Spm | same family |
| chr2 | 1556737 | 7.30E-12 | ATCOPIA10-LTR/Copia | same family |
| chr2 | 15592321 | 5.49E-23 | ATMU7-DNA/MuDR | same family |
| chr2 | 1570882 | 1.05E-11 | ATGP5-LTR/Gypsy | same family |
| chr2 | 1657991 | 3.61E-10 | ATMU3N1-DNA/MuDR | same family |
| chr2 | 1684493 | 1.62E-20 | ATDNA2T9B-DNA/MuDR | same family |
| chr2 | 173191 | 3.37E-09 | ATMU3-DNA/MuDR | same family |
| chr2 | 18106620 | 1.32E-19 | RathE2_cons-RathE2_cons | same family |
| chr2 | 18712298 | 1.16E-74 | ATCOPIA57-LTR/Copia | same family |
| chr2 | 1970254 | 4.72E-99 | ATCOPIA64-LTR/Copia | same family |
| chr2 | 2038048 | 5.31E-12 | ATN9_1-DNA/MuDR | same family |
| chr2 | 2072254 | 8.52E-16 | ATGP3-LTR/Gypsy | same family |
| chr2 | 2312467 | 9.70E-18 | VANDALNX2-DNA/MuDR | same family |
| chr2 | 2502209 | 1.65E-09 | ATENSPM4-DNA/En-Spm | same family |
| chr2 | 2731919 | 3.19E-10 | ATLINE1_4-LINE/L1 | same family |
| chr2 | 2778128 | 1.94E-16 | ATCOPIA68-LTR/Copia | same family |
| chr2 | 2882240 | 8.36E-38 | ATCOPIA8A-LTR/Copia | same family |
| chr2 | 2907791 | 1.06E-12 | ATCOPIA13-LTR/Copia | same family |
| chr2 | 2952411 | 2.20E-51 | ATMU9-DNA/MuDR | same family |
| chr2 | 3059644 | 1.95E-17 | ATCOPIA20-LTR/Copia | same family |
| chr2 | 3115454 | 1.03E-12 | ARNOLD4-DNA/MuDR | same family |
| chr2 | 3120130 | 2.33E-16 | ATCOPIA32-LTR/Copia | same family |
| chr2 | 3189562 | 1.74E-10 | DT1-DNA/Mariner | others |
| chr2 | 3189562 | 2.27E-13 | VANDAL16-DNA/MuDR | others |
| chr2 | 3189562 | 4.63E-10 | ATHILA4B_LTR-LTR/Gypsy | others |
| chr2 | 4034562 | 1.48E-09 | ATENSPM1-DNA/En-Spm | same family |
| chr2 | 4071403 | 1.06E-49 | ATCOPIA42-LTR/Copia | same family |
| chr2 | 4091777 | 7.21E-27 | ATCOPIA35-LTR/Copia | same family |
| chr2 | 4152747 | 9.11E-10 | ATREP16-DNA/MuDR | same family |
| chr2 | 4328548 | 1.03E-09 | ARNOLD4-DNA/MuDR | same family |
| chr2 | 4348901 | 4.83E-12 | ATENSPM1A-DNA/En-Spm | others |
| chr2 | 4603036 | 1.20E-12 | ATCOPIA34-LTR/Copia | same family |
| chr2 | 4685333 | 5.23E-10 | ATHILA0_I-LTR/Gypsy | same family |
| chr2 | 5166963 | 4.34E-22 | VANDAL9-DNA/MuDR | others |
| chr2 | 5393279 | 1.05E-09 | ATCOPIA27-LTR/Copia | same family |
| chr2 | 5672297 | 6.08E-23 | VANDALNX2-DNA/MuDR | same family |
| chr2 | 5835322 | 4.20E-09 | ATREP12-RC/Helitron | others |
| chr2 | 5906544 | 1.13E-09 | ATHILA7A-LTR/Gypsy | others |

| | | | | |
|---|---|---|---|---|
| chr2 | 613432 | 4.61E-17 | RathE2_cons-RathE2_cons | same family |
| chr2 | 623834 | 4.44E-13 | ATCOPIA63-LTR/Copia | same family |
| chr2 | 6419026 | 1.29E-13 | ATCOPIA25-LTR/Copia | same family |
| chr2 | 6421720 | 5.85E-114 | ATCOPIA9-LTR/Copia | same family |
| chr2 | 6481956 | 2.85E-09 | SIMPLEHAT2-DNA/HAT | same family |
| chr2 | 6485147 | 3.81E-09 | DT1-DNA/Mariner | same family |
| chr2 | 6508164 | 1.71E-28 | ATRE1-LTR/Copia | same family |
| chr2 | 6864213 | 3.58E-09 | ATLINE1_6-LINE/L1 | same family |
| chr2 | 6889853 | 1.26E-19 | VANDAL18NA-DNA/MuDR | same family |
| chr2 | 6906197 | 5.64E-40 | ATCOPIA80-LTR/Copia | others |
| chr2 | 6909271 | 8.11E-142 | ATCOPIA38-LTR/Copia | same family |
| chr2 | 6953263 | 2.13E-21 | ATCOPIA38B-LTR/Copia | same family |
| chr2 | 7058161 | 1.37E-18 | ATCOPIA61-LTR/Copia | same family |
| chr2 | 71953 | 2.93E-12 | ATCOPIA27-LTR/Copia | others |
| chr2 | 7223612 | 4.08E-202 | ATCOPIA70-LTR/Copia | same family |
| chr2 | 7423364 | 5.54E-10 | VANDAL18NB-DNA/MuDR | same family |
| chr2 | 7438905 | 3.62E-10 | ATREP14-RC/Helitron | same family |
| chr2 | 7448151 | 1.81E-16 | ATCOPIA71-LTR/Copia | same family |
| chr2 | 751219 | 6.80E-11 | ATTIRTA1-DNA/Tc1 | same family |
| chr2 | 7598016 | 6.93E-67 | ATCOPIA39-LTR/Copia | same family |
| chr2 | 7894427 | 1.06E-20 | ATCOPIA48-LTR/Copia | same family |
| chr2 | 8088582 | 4.67E-14 | ATMUN1-DNA/MuDR | same family |
| chr2 | 8367770 | 2.84E-16 | VANDAL15-DNA/MuDR | same family |
| chr2 | 8560185 | 2.17E-12 | ATCOPIA69-LTR/Copia | same family |
| chr2 | 905303 | 3.75E-13 | ATCOPIA7-LTR/Copia | others |
| chr2 | 9060259 | 3.67E-18 | VANDAL5A-DNA/MuDR | same family |
| chr2 | 907295 | 7.13E-09 | ATMUN1-DNA/MuDR | others |
| chr2 | 9117263 | 3.09E-45 | ATCOPIA74-LTR/Copia | same family |
| chr2 | 935080 | 2.29E-15 | ATCOPIA56-LTR/Copia | same family |
| chr2 | 9443780 | 1.06E-15 | RathE2_cons-RathE2_cons | same family |
| chr2 | 9489659 | 1.75E-11 | ATCOPIA9-LTR/Copia | same family |
| chr2 | 9808470 | 2.19E-56 | ATCOPIA87-LTR/Copia | same family |
| chr3 | 10106026 | 5.68E-52 | ATCOPIA49-LTR/Copia | same family |
| chr3 | 10450322 | 8.34E-38 | ATHATN1-DNA/HAT | same family |
| chr3 | 10522685 | 7.88E-37 | ATCOPIA37-LTR/Copia | same family |
| chr3 | 10544809 | 2.20E-120 | ATCOPIA31A-LTR/Copia | same family |
| chr3 | 10880665 | 1.07E-16 | ATCOPIA3-LTR/Copia | same family |
| chr3 | 10923253 | 1.56E-20 | ATLINE1_4-LINE/L1 | same family |
| chr3 | 10923253 | 6.17E-11 | ATLINE1_6-LINE/L1 | same family |
| chr3 | 11021693 | 1.24E-24 | ATCOPIA92-LTR/Copia | same family |
| chr3 | 11021693 | 1.34E-62 | ATCOPIA5-LTR/Copia | same family |
| chr3 | 11021693 | 3.48E-16 | ATCOPIA84-LTR/Copia | same family |
| chr3 | 11081241 | 7.82E-24 | VANDAL18NB-DNA/MuDR | same family |
| chr3 | 11122535 | 6.02E-96 | ATCOPIA14-LTR/Copia | same family |
| chr3 | 11182577 | 1.25E-14 | VANDAL6-DNA/MuDR | same family |
| chr3 | 11651213 | 1.38E-27 | ATCOPIA63-LTR/Copia | same family |
| chr3 | 11697052 | 1.87E-29 | ATCOPIA16-LTR/Copia | same family |

| | | | | |
|---|---|---|---|---|
| chr3 | 11702428 | 3.46E-60 | ATCOPIA89-LTR/Copia | same family |
| chr3 | 11813775 | 1.56E-43 | ATHAT7-DNA/HAT | others |
| chr3 | 11859836 | 1.75E-14 | BOMZH1-DNA/MuDR | same family |
| chr3 | 11952384 | 1.32E-61 | ATGP6-LTR/Gypsy | same family |
| chr3 | 12116218 | 2.38E-11 | ATENSPM4-DNA/En-Spm | same family |
| chr3 | 12380198 | 1.57E-10 | Unassigned-Unassigned | same family |
| chr3 | 12446418 | 2.86E-37 | ATCOPIA63-LTR/Copia | others |
| chr3 | 12468067 | 2.30E-12 | ATCOPIA43-LTR/Copia | same family |
| chr3 | 12641372 | 6.56E-16 | ATGP8-LTR/Gypsy | others |
| chr3 | 13574424 | 7.97E-10 | ATENSPM11-DNA/En-Spm | same family |
| chr3 | 13577002 | 1.81E-11 | ATDNA1T9A-DNA/MuDR | others |
| chr3 | 14328918 | 2.03E-16 | ATGP9B-LTR/Gypsy | others |
| chr3 | 14868498 | 8.73E-17 | ATCOPIA34-LTR/Copia | same family |
| chr3 | 15386215 | 8.60E-55 | ATCOPIA33-LTR/Copia | same family |
| chr3 | 15402284 | 8.06E-10 | ATGP5-LTR/Gypsy | same family |
| chr3 | 15598975 | 3.67E-20 | ATCOPIA43-LTR/Copia | same family |
| chr3 | 15617992 | 4.75E-22 | ATCOPIA16-LTR/Copia | same family |
| chr3 | 15768731 | 1.14E-41 | ATCOPIA62-LTR/Copia | same family |
| chr3 | 15963467 | 2.68E-20 | ATCOPIA14-LTR/Copia | same family |
| chr3 | 16067491 | 2.91E-11 | ATMU5-DNA/MuDR | others |
| chr3 | 16395324 | 3.03E-45 | ATCOPIA10-LTR/Copia | same family |
| chr3 | 16433227 | 7.85E-14 | ATCOPIA1-LTR/Copia | same family |
| chr3 | 16461148 | 1.30E-12 | ATCOPIA59-LTR/Copia | others |
| chr3 | 16604525 | 4.17E-22 | ATHAT2-DNA/HAT | same family |
| chr3 | 16700656 | 1.69E-53 | ATCOPIA32-LTR/Copia | same family |
| chr3 | 16764373 | 4.24E-26 | ATCOPIA13-LTR/Copia | same family |
| chr3 | 16814862 | 1.83E-31 | ATCOPIA81-LTR/Copia | same family |
| chr3 | 16888129 | 1.56E-43 | ATCOPIA45-LTR/Copia | same family |
| chr3 | 16978660 | 4.05E-11 | ATCOPIA79-LTR/Copia | others |
| chr3 | 17118337 | 1.05E-10 | ATCOPIA47-LTR/Copia | same family |
| chr3 | 17225920 | 4.46E-14 | ATHILA4D_LTR-LTR/Gypsy | same family |
| chr3 | 17606534 | 2.70E-12 | DT1-DNA/Mariner | same family |
| chr3 | 18181126 | 2.83E-44 | ATHPOGON2-DNA/Pogo | same family |
| chr3 | 18458178 | 1.48E-09 | ATN9_1-DNA/MuDR | same family |
| chr3 | 18787205 | 5.47E-29 | ATCOPIA52-LTR/Copia | same family |
| chr3 | 18925937 | 3.43E-13 | ATTIRTA1-DNA/Tc1 | same family |
| chr3 | 19884704 | 2.71E-25 | VANDAL11-DNA/MuDR | same family |
| chr3 | 21766850 | 2.88E-10 | SADHU-Unassigned | same family |
| chr3 | 21982234 | 2.21E-24 | ATHATN10-DNA/HAT | same family |
| chr3 | 2866904 | 9.06E-10 | VANDAL16-DNA/MuDR | same family |
| chr3 | 2960095 | 3.79E-14 | ATMU6N1-DNA/MuDR | same family |
| chr3 | 3694956 | 1.70E-11 | ATLINE1_4-LINE/L1 | others |
| chr3 | 4378113 | 1.24E-14 | SADHU-Unassigned | same family |
| chr3 | 6684040 | 5.46E-63 | ATCOPIA54-LTR/Copia | same family |
| chr3 | 7637497 | 2.50E-15 | ATCOPIA13-LTR/Copia | same family |
| chr3 | 8105155 | 4.36E-12 | ATDNAI26T9-DNA/MuDR | same family |
| chr3 | 8841794 | 2.17E-20 | VANDAL2-DNA/MuDR | same family |

| | | | | |
|---|---|---|---|---|
| chr3 | 9232206 | 1.11E-31 | ATCOPIA65-LTR/Copia | same family |
| chr3 | 9243249 | 3.49E-13 | ATCOPIA18-LTR/Copia | others |
| chr3 | 9276243 | 3.98E-13 | ATLINE1_4-LINE/L1 | same family |
| chr3 | 9508890 | 2.11E-10 | ATCOPIA24-LTR/Copia | others |
| chr3 | 9646870 | 2.84E-17 | RathE2_cons-RathE2_cons | same family |
| chr4 | 10247683 | 6.16E-203 | ATHATN9-DNA/HAT | same family |
| chr4 | 10930204 | 1.25E-10 | SIMPLEHAT2-DNA/HAT | same family |
| chr4 | 1095114 | 8.83E-17 | ATCOPIA38B-LTR/Copia | same family |
| chr4 | 10975231 | 3.85E-33 | ATCOPIA47-LTR/Copia | same family |
| chr4 | 11803853 | 2.61E-09 | ATHATN3A-DNA/HAT | others |
| chr4 | 11833351 | 1.52E-17 | HARBINGER-DNA/Harbinger | same family |
| chr4 | 12901450 | 5.92E-12 | ATTIRX1C-DNA | same family |
| chr4 | 14266531 | 7.91E-123 | ATCOPIA46-LTR/Copia | same family |
| chr4 | 14910847 | 2.65E-09 | ATREP12-RC/Helitron | others |
| chr4 | 1542814 | 8.00E-16 | SADHU-Unassigned | others |
| chr4 | 15568558 | 8.86E-17 | ATGP9B-LTR/Gypsy | same family |
| chr4 | 16831625 | 1.37E-12 | ATMU3-DNA/MuDR | others |
| chr4 | 17798768 | 4.95E-20 | ATCOPIA71-LTR/Copia | same family |
| chr4 | 18426760 | 1.57E-70 | ATREP9-RC/Helitron | same family |
| chr4 | 2034021 | 2.53E-21 | ATRE1-LTR/Copia | same family |
| chr4 | 2188759 | 1.57E-31 | ATCOPIA93-LTR/Copia | same family |
| chr4 | 2198043 | 1.68E-14 | ATCOPIA58-LTR/Copia | same family |
| chr4 | 2204593 | 2.22E-25 | ATCOPIA44-LTR/Copia | same family |
| chr4 | 2234944 | 8.42E-21 | RathE3_cons-RathE3_cons | same family |
| chr4 | 2245163 | 1.59E-15 | VANDAL18-DNA/MuDR | same family |
| chr4 | 2304027 | 4.71E-40 | ATCOPIA10-LTR/Copia | same family |
| chr4 | 2324899 | 1.04E-21 | ATLINE1_5-LINE/L1 | same family |
| chr4 | 2570520 | 2.99E-13 | DT1-DNA/Mariner | same family |
| chr4 | 2595271 | 7.19E-20 | ATCOPIA1-LTR/Copia | same family |
| chr4 | 2604435 | 1.14E-12 | ATCOPIA69-LTR/Copia | same family |
| chr4 | 2615926 | 3.01E-18 | ATHPOGON1-DNA/Pogo | same family |
| chr4 | 2641470 | 3.49E-09 | ATCOPIA56-LTR/Copia | same family |
| chr4 | 2656183 | 1.20E-18 | ATTIRX1D-DNA | same family |
| chr4 | 2687921 | 1.56E-35 | ATCOPIA8B-LTR/Copia | same family |
| chr4 | 4084529 | 1.01E-15 | ATGP6-LTR/Gypsy | same family |
| chr4 | 4171045 | 2.34E-16 | ATGP7-LTR/Gypsy | same family |
| chr4 | 4739600 | 4.17E-15 | ATHAT2-DNA/HAT | same family |
| chr4 | 4839451 | 9.50E-13 | ARNOLD3-DNA/MuDR | same family |
| chr4 | 4887044 | 1.98E-11 | ATHILA0_I-LTR/Gypsy | same family |
| chr4 | 5252651 | 2.53E-29 | ATHATN4-DNA/HAT | same family |
| chr4 | 5267074 | 5.79E-09 | RathE3_cons-RathE3_cons | others |
| chr4 | 5303557 | 2.15E-13 | ATCOPIA44-LTR/Copia | same family |
| chr4 | 5303730 | 5.99E-22 | ATHATN8-DNA/HAT | same family |
| chr4 | 5303730 | 6.23E-123 | ATCOPIA55-LTR/Copia | same family |
| chr4 | 5425380 | 3.17E-16 | ATCOPIA93-LTR/Copia | same family |
| chr4 | 5439622 | 1.08E-44 | ATCOPIA37-LTR/Copia | same family |
| chr4 | 5492861 | 1.90E-09 | VANDALNX2-DNA/MuDR | same family |

| | | | | |
|---|---|---|---|---|
| chr4 | 5614580 | 1.26E-10 | ATGP9B-LTR/Gypsy | same family |
| chr4 | 5643994 | 8.35E-11 | ATLINE1_1-LINE/L1 | same family |
| chr4 | 5790554 | 7.73E-11 | BOMZH1-DNA/MuDR | same family |
| chr4 | 5921463 | 3.21E-27 | ATRE1-LTR/Copia | same family |
| chr4 | 5937137 | 1.73E-28 | ATCOPIA69-LTR/Copia | same family |
| chr4 | 6123153 | 6.45E-22 | BOMZH2-DNA/MuDR | same family |
| chr4 | 6353065 | 1.56E-31 | ATCOPIA88-LTR/Copia | same family |
| chr4 | 637062 | 1.08E-13 | ATLINE1_6-LINE/L1 | same family |
| chr4 | 6453697 | 3.78E-09 | ATGP3-LTR/Gypsy | same family |
| chr4 | 6466396 | 3.88E-82 | ATCOPIA50-LTR/Copia | same family |
| chr4 | 6507387 | 2.19E-55 | TA1_AT-LTR/Gypsy | same family |
| chr4 | 6594104 | 4.11E-55 | ATCOPIA7-LTR/Copia | same family |
| chr4 | 6914365 | 5.15E-134 | ATCOPIA22-LTR/Copia | same family |
| chr4 | 6971989 | 2.96E-16 | ATCOPIA57-LTR/Copia | same family |
| chr4 | 7005996 | 5.12E-09 | ATREP16-DNA/MuDR | others |
| chr4 | 7300169 | 2.76E-12 | ATLINE1_1-LINE/L1 | same family |
| chr4 | 7304640 | 2.69E-11 | ATCOPIA48-LTR/Copia | same family |
| chr4 | 7330268 | 1.83E-09 | ATLINE1_6-LINE/L1 | same family |
| chr4 | 7592733 | 1.63E-40 | ATCOPIA34-LTR/Copia | same family |
| chr4 | 7809793 | 7.36E-12 | ATHPOGON1-DNA/Pogo | same family |
| chr4 | 7831727 | 1.18E-09 | ATREP14-RC/Helitron | others |
| chr4 | 7897390 | 3.02E-30 | ATREP8-RC/Helitron | same family |
| chr4 | 7947027 | 4.72E-16 | ATCOPIA79-LTR/Copia | others |
| chr4 | 8587578 | 5.49E-47 | ATTIRX1D-DNA | same family |
| chr4 | 8914809 | 4.11E-35 | ATCOPIA48-LTR/Copia | same family |
| chr5 | 10022210 | 9.60E-10 | ATCOPIA31-LTR/Copia | others |
| chr5 | 10102514 | 5.74E-10 | VANDAL14-DNA/MuDR | others |
| chr5 | 10179768 | 8.67E-15 | ATGP6-LTR/Gypsy | others |
| chr5 | 10397122 | 2.12E-12 | ATCOPIA68-LTR/Copia | same family |
| chr5 | 10587991 | 3.62E-12 | ATMU8-DNA/MuDR | same family |
| chr5 | 10648398 | 1.01E-27 | ATCOPIA44-LTR/Copia | same family |
| chr5 | 10719195 | 1.25E-11 | ATHATN10-DNA/HAT | same family |
| chr5 | 10837909 | 5.75E-11 | ATENSPM11-DNA/En-Spm | same family |
| chr5 | 12315675 | 9.04E-12 | ATHILA4D_LTR-LTR/Gypsy | same family |
| chr5 | 12517374 | 6.85E-13 | HARBINGER-DNA/Harbinger | same family |
| chr5 | 12606542 | 2.40E-55 | ATCOPIA88-LTR/Copia | same family |
| chr5 | 12904117 | 2.54E-27 | ATCOPIA43-LTR/Copia | others |
| chr5 | 13083991 | 4.45E-10 | ATGP8-LTR/Gypsy | same family |
| chr5 | 133889 | 1.00E-09 | Unassigned-Unassigned | same family |
| chr5 | 13737229 | 3.36E-75 | ATCOPIA24-LTR/Copia | same family |
| chr5 | 13792376 | 5.20E-10 | ATHAT3-DNA/HAT | same family |
| chr5 | 13810698 | 7.05E-17 | ATHATN7-DNA/HAT | same family |
| chr5 | 13877920 | 3.46E-17 | ATCOPIA67-LTR/Copia | same family |
| chr5 | 13942794 | 4.54E-29 | ATCOPIA12-LTR/Copia | same family |
| chr5 | 13949340 | 9.95E-44 | ATCOPIA75-LTR/Copia | same family |
| chr5 | 13993917 | 5.29E-34 | ATCOPIA56-LTR/Copia | same family |
| chr5 | 14607204 | 1.06E-12 | ATCOPIA57-LTR/Copia | same family |

| | | | | |
|---|---|---|---|---|
| chr5 | 14649603 | 1.33E-09 | VANDAL18NB-DNA/MuDR | same family |
| chr5 | 14650025 | 1.74E-17 | ATCOPIA62-LTR/Copia | same family |
| chr5 | 14822011 | 1.11E-10 | ATDNA2T9C-DNA/MuDR | same family |
| chr5 | 14993632 | 3.18E-14 | ATREP17-DNA/MuDR | same family |
| chr5 | 15015836 | 4.33E-14 | BOMZH1-DNA/MuDR | same family |
| chr5 | 15175171 | 9.19E-21 | ATCOPIA87-LTR/Copia | same family |
| chr5 | 15334040 | 1.96E-13 | Unassigned-Unassigned | same family |
| chr5 | 15645894 | 1.12E-46 | ATCOPIA25-LTR/Copia | same family |
| chr5 | 15659612 | 3.63E-34 | ATCOPIA83-LTR/Copia | same family |
| chr5 | 15703009 | 1.19E-26 | RathE2_cons-RathE2_cons | same family |
| chr5 | 16030044 | 2.92E-10 | ATDNAI26T9-DNA/MuDR | others |
| chr5 | 16971608 | 1.18E-10 | ATMU3N1-DNA/MuDR | same family |
| chr5 | 17089240 | 1.57E-09 | ATCOPIA61-LTR/Copia | same family |
| chr5 | 17538386 | 9.69E-200 | ATCOPIA77-LTR/Copia | same family |
| chr5 | 17744373 | 3.57E-22 | ATDNAI27T9B-DNA/MuDR | same family |
| chr5 | 17968425 | 6.88E-23 | ATCOPIA71-LTR/Copia | same family |
| chr5 | 18174460 | 3.06E-58 | VANDAL18NA-DNA/MuDR | same family |
| chr5 | 18230709 | 6.84E-39 | ATCOPIA23-LTR/Copia | same family |
| chr5 | 18490555 | 4.28E-49 | ATCOPIA31-LTR/Copia | same family |
| chr5 | 18642012 | 6.55E-14 | ATENSPM4-DNA/En-Spm | same family |
| chr5 | 18930031 | 2.74E-13 | ATCOPIA58-LTR/Copia | same family |
| chr5 | 19176529 | 1.53E-19 | ATLINE1_4-LINE/L1 | same family |
| chr5 | 19358287 | 4.56E-19 | Unassigned-Unassigned | same family |
| chr5 | 19501005 | 4.71E-49 | ATCOPIA42-LTR/Copia | same family |
| chr5 | 19528022 | 2.92E-21 | RP1_AT-DNA | same family |
| chr5 | 1964239 | 3.93E-17 | ATCOPIA63-LTR/Copia | others |
| chr5 | 20730598 | 1.79E-15 | ATENSPM1-DNA/En-Spm | same family |
| chr5 | 21562910 | 3.41E-10 | ATCOPIA73-LTR/Copia | others |
| chr5 | 21852815 | 2.32E-24 | ATCOPIA16-LTR/Copia | same family |
| chr5 | 22956136 | 3.26E-25 | ATCOPIA61-LTR/Copia | same family |
| chr5 | 22957638 | 2.13E-18 | ATCOPIA62-LTR/Copia | same family |
| chr5 | 24003873 | 2.11E-09 | ATREP12-RC/Helitron | same family |
| chr5 | 3258283 | 7.67E-19 | ATGP9LTR-LTR/Gypsy | same family |
| chr5 | 5947440 | 2.71E-15 | ATGP9B-LTR/Gypsy | same family |
| chr5 | 6408237 | 5.60E-63 | ATCOPIA89-LTR/Copia | same family |
| chr5 | 6430050 | 1.63E-31 | ATCOPIA53-LTR/Copia | same family |
| chr5 | 6433221 | 4.63E-251 | ATCOPIA90-LTR/Copia | same family |
| chr5 | 6906427 | 1.04E-17 | ATMU3N1-DNA/MuDR | same family |
| chr5 | 6909846 | 8.44E-14 | ATCOPIA36-LTR/Copia | same family |
| chr5 | 7041065 | 1.17E-24 | VANDAL20-DNA/MuDR | same family |
| chr5 | 7434205 | 3.28E-15 | ATENSPM4-DNA/En-Spm | same family |
| chr5 | 8123700 | 9.04E-12 | ATMUN1-DNA/MuDR | same family |
| chr5 | 8210570 | 6.37E-30 | ATDNA1T9A-DNA/MuDR | same family |
| chr5 | 8406755 | 2.80E-18 | ATN9_1-DNA/MuDR | same family |
| chr5 | 8441442 | 7.96E-23 | ATREP18-DNA | others |
| chr5 | 8445178 | 8.14E-09 | ATCOPIA57-LTR/Copia | others |
| chr5 | 8523828 | 1.17E-11 | VANDAL14-DNA/MuDR | same family |

| | | | | |
|---|---|---|---|---|
| chr5 | 8686371 | 1.30E-61 | ATCOPIA24-LTR/Copia | same family |
| chr5 | 8690188 | 6.31E-16 | ATCOPIA35-LTR/Copia | same family |
| chr5 | 8927979 | 7.48E-14 | ATCOPIA27-LTR/Copia | same family |
| chr5 | 9029774 | 3.62E-19 | ATCOPIA38B-LTR/Copia | same family |
| chr5 | 9136863 | 1.56E-50 | ATCOPIA80-LTR/Copia | others |
| chr5 | 9210060 | 1.47E-11 | ATHILA7A-LTR/Gypsy | same family |
| chr5 | 9276156 | 2.84E-19 | ATCOPIA63-LTR/Copia | same family |
| chr5 | 9708994 | 1.86E-35 | ATMU5-DNA/MuDR | same family |

Note: In peak type, same family represents a peak contain TEs of the same family, others represents a peak not contain TEs of the same family.

Table S8. Summary of causal TEs associated with TE family copy number variation.

| **Lead SNP** | **TE family** | **Causal TE** |
|---|---|---|
| chr2:4328548 | ARNOLD4-DNA/MuDR | AT2TE18120 |
| chr4:5303557 | ATCOPIA44-LTR/Copia | AT4TE20845 |
| chr1:19003520 | ATCOPIA68-LTR/Copia | AT1TE62960 |
| chr1:14243988 | ATHATN1-DNA/HAT | AT1TE46880, AT1TE46890, AT1TE46895, AT1TE46900, AT1TE46905 |
| chr1:15890614 | ATHATN1-DNA/HAT | AT1TE52290 |

Table S9. Summary of candidate genes in the GWAS analysis of TE family copy number.

| Lead SNP | TE family | Gene |
|---|---|---|
| chr2:4328548 | ARNOLD4-DNA/MuDR | AT2G10940,AT2G10950,AT2G10955,AT2G10965,AT2G10970,AT2G10975,AT2G11005,AT2G11000 |
| chr3:9243249 | ATCOPIA18-LTR/Copia | AT3G25430,AT3G25460,AT3G25470,AT3G25480,AT3G25490,AT3G25493,AT3G25500,AT3G25505,AT3G25510,AT3G25440 |
| chr3:9508890 | ATCOPIA24-LTR/Copia | AT3G25910,AT3G25920,AT3G25930,AT3G25940,AT3G25950,AT3G25960,AT3G25970,AT3G25980,AT3G25990,AT3G26000,AT3G26010,AT3G26020,AT3G26040,AT3G26050,AT3G26060,AT3G26070,AT3G26080,AT3G26030 |
| chr2:71953 | ATCOPIA27-LTR/Copia | AT2G01050,AT2G01060,AT2G01070,AT2G01080,AT2G01090,AT2G01110,AT2G01120,AT2G01130,AT2G01140,AT2G01150,AT2G01170,AT2G01100 |
| chr1:13109197 | ATCOPIA33-LTR/Copia | AT1G35140,AT1G35150,AT1G35160,AT1G35170,AT1G35181,AT1G35183,AT1G35190,AT1G35210,AT1G35215,AT1G35230,AT1G35240,AT1G35242,AT1G35250,AT1G35255,AT1G35260,AT1G35290,AT1G35310,AT1G35330,AT1G35340,AT1G35350,AT1G35353,AT1G35365,AT1G35375,AT1G35400,AT1G35410,AT1G35420,AT1G35430,AT1G35435,AT1G35440,AT1G35467,AT1G35470,AT1G35490,AT1G35500,AT1G35515,AT1G35516,AT1G35520,AT1G35537,AT1G35540,AT1G35550,AT1G35560,AT1G35610,AT1G35614,AT1G35617,AT1G35625,AT1G35630,AT1G35660,AT1G35680,AT1G35180,AT1G35220,AT1G35320,AT1G35460,AT1G35510,AT1G35530,AT1G35580,AT1G35620,AT1G35670,AT1G35710 |
| chr2:10891327 | ATCOPIA39-LTR/Copia | AT2G25470,AT2G25480,AT2G25490,AT2G25500,AT2G25510,AT2G25520,AT2G25530,AT2G25540,AT2G25560,AT2G25565,AT2G25570,AT2G25580,AT2G25590,AT2G25605,AT2G25610,AT2G25620,AT2G25625,AT2G25630,AT2G25482,AT2G25600,AT2G25640 |
| chr5:12904117 | ATCOPIA43-LTR/Copia | AT5G32670,AT5G33210,AT5G33290,AT5G33300,AT5G33320,AT5G33330,AT5G33340,AT5G33370,AT5G33393,AT5G33390,AT5G33406,AT5G33806,AT5G33898,AT5G34581,AT5G34780,AT5G34830,AT5G34829,AT5G34869,AT5G34828,AT5G34850,AT5G34870,AT5G34882,AT5G34881,AT5G34883,AT5G34885,AT5G34887,AT5G34905,AT5G34908,AT5G34930,AT5G35050,AT5G35067,AT5G35080,AT5G35090,AT5G35100,AT5G35110,AT5G35120,AT5G33280,AT5G33355,AT5G35069,AT5G34940 |
| chr4:5303557 | ATCOPIA44-LTR/Copia | AT4G07940,AT4G07950,AT4G07960,AT4G07965,AT4G07990,AT4G07995,AT4G08025,AT4G08028,AT4G08039,AT4G08040,AT4G08097,AT4G08140,AT4G08150,AT4G08170,AT4G08180,AT4G08190,AT4G08210,AT4G08230,AT4G08240,AT4G08250,AT4G08260,AT4G08263,AT4G08267,AT4G08270,AT4G08280,AT4G08290,AT4G08300,AT4G08310,AT4G08320,AT4G08330,AT4G08360,AT4G08370,AT4G08380,AT4G08390,AT4G08395,AT4G08406,AT4G08430,AT4G08450,AT4G08455,AT4G08460,AT4G08470,AT4G08160,AT4G08350,AT4G08400,AT4G08410 |
| chr1:18550465 | ATCOPIA52-LTR/Copia | AT1G50040,AT1G50060,AT1G50080,AT1G50090,AT1G50110,AT1G50030,AT1G50050,AT1G50120 |

| | | |
|---|---|---|
| chr1:3636643 | ATCOPIA53-LTR/Copia | AT1G10875,AT1G10880,AT1G10890,AT1G10910,AT1G10920,AT1G10870,AT1G10900,AT1G10930 |
| chr1:14212288 | ATCOPIA57-LTR/Copia | AT1G37140,AT1G37150,AT1G38065,AT1G38131 |
| chr1:15561355 | ATCOPIA57-LTR/Copia | AT1G41820,AT1G41830,AT1G40390 |
| chr3:16461148 | ATCOPIA59-LTR/Copia | AT3G45000,AT3G45010,AT3G45020,AT3G45030,AT3G45040,AT3G45050,AT3G45060,AT3G45070 |
| chr1:2071605 | ATCOPIA59-LTR/Copia | AT1G06670,AT1G06690,AT1G06700,AT1G06710,AT1G06730,AT1G06740,AT1G06750,AT1G06760,AT1G06780,AT1G06680,AT1G06720,AT1G06770 |
| chr3:12446418 | ATCOPIA63-LTR/Copia | AT3G30720,AT3G30725,AT3G30730,AT3G30770,AT3G30775,AT3G30739 |
| chr5:1964239 | ATCOPIA63-LTR/Copia | AT5G06360,AT5G06370,AT5G06390,AT5G06400,AT5G06410,AT5G06420,AT5G06430,AT5G06450,AT5G06460,AT5G06470,AT5G06480,AT5G06490,AT5G06380,AT5G06500,AT5G06440 |
| chr1:19003520 | ATCOPIA68-LTR/Copia | AT1G51200,AT1G51220,AT1G51230,AT1G51240,AT1G51250,AT1G51270,AT1G51290,AT1G51300,AT1G51310,AT1G51210,AT1G51260 |
| chr5:21562910 | ATCOPIA73-LTR/Copia | AT5G53140,AT5G53150,AT5G53170,AT5G53180,AT5G53190,AT5G53200,AT5G53210,AT5G53220,AT5G53230,AT5G53240,AT5G53160 |
| chr4:7947027 | ATCOPIA79-LTR/Copia | AT4G13620,AT4G13630,AT4G13640,AT4G13650,AT4G13670,AT4G13680,AT4G13690,AT4G13700,AT4G13660 |
| chr2:905303 | ATCOPIA7-LTR/Copia | AT2G03050,AT2G03070,AT2G03110,AT2G03120,AT2G03130,AT2G03150,AT2G03160,AT2G03170,AT2G03180,AT2G03190,AT2G03210,AT2G03220,AT2G03240,AT2G03230,AT2G03060,AT2G03140,AT2G03200,AT2G03090 |
| chr5:9136863 | ATCOPIA80-LTR/Copia | AT5G26050,AT5G26060,AT5G26070,AT5G26080,AT5G26100,AT5G26110,AT5G26130,AT5G26140,AT5G26150,AT5G26160,AT5G26170,AT5G26180,AT5G26190,AT5G26200,AT5G26220,AT5G26230,AT5G26090,AT5G26120,AT5G26210 |
| chr2:6906197 | ATCOPIA80-LTR/Copia | AT2G15450,AT2G15460,AT2G15470,AT2G15480,AT2G15490,AT2G15500,AT2G15530,AT2G15535,AT2G15560,AT2G15570,AT2G15580,AT2G15590,AT2G15610,AT2G15620,AT2G15640,AT2G15660,AT2G15670,AT2G15680,AT2G15695,AT2G15710,AT2G15730,AT2G15740,AT2G15760,AT2G15770,AT2G15780,AT2G15790,AT2G15820,AT2G15830,AT2G15860,AT2G15880,AT2G15890,AT2G15910,AT2G15960,AT2G15970,AT2G15980,AT2G16005,AT2G16015,AT2G16016,AT2G16018,AT2G16019,AT2G16020,AT2G16030,AT2G16040,AT2G16050,AT2G16060,AT2G16070,AT2G16090,AT2G16120,AT2G16130,AT2G16190,AT2G16200,AT2G16220,AT2G16225,AT2G16230,AT2G16250,AT2G16270,AT2G16280,AT2G16290,AT2G16300,AT2G16340,AT2G16360,AT2G16370,AT2G16380,AT2G16385,AT2G16390,AT2G16400,AT2G16405,AT2G16430,AT2G16450,AT2G16460,AT2G16490,AT2G16500,AT2G16505,AT2G16520,AT2G16530,AT2G16535,AT2G16570,AT2G16580,AT2G16575,AT2G16586,AT2G16592,AT2G16594,AT2G16595,AT2G16600,AT2G16620,AT2G16630,AT2G16640,AT2G16650,AT2G16660,AT2G16676,AT2G16700,AT2G16710,AT2G16720,AT2G16730,AT2G16740,AT2G16760,AT2G16770,AT2G15690,AT2G15900,AT2G16210,AT2G16440,AT2G16485,AT2G16510,AT2G16750,AT2G15630,AT2G16365 |
| chr3:13577002 | ATDNA1T9A-DNA/MuDR | AT3G32960,AT3G32980 |

| | | |
|---|---|---|
| chr5:16030044 | ATDNAI26T9-DNA/MuDR | AT5G39940,AT5G39960,AT5G39970,AT5G39980,AT5G39990,AT5G40000,AT5G40010,AT5G40030,AT5G40040,AT5G40050,AT5G40060,AT5G40070,AT5G40090,AT5G40100,AT5G40020,AT5G40080,AT5G39950 |
| chr2:4348901 | ATENSPM1A-DNA/En-Spm | AT2G10975,AT2G11005,AT2G11010,AT2G11015,AT2G11025,AT2G11000 |
| chr2:13922516 | ATGP3-LTR/Gypsy | AT2G32780,AT2G32785,AT2G32788,AT2G32790,AT2G32800,AT2G32810,AT2G32830,AT2G32835,AT2G32840,AT2G32850,AT2G32870,AT2G32880,AT2G32885,AT2G32890,AT2G32900,AT2G32905,AT2G32920,AT2G32820,AT2G32860,AT2G32910 |
| chr1:14446668 | ATGP5-LTR/Gypsy | AT1G37113,AT1G37130,AT1G37140,AT1G37150,AT1G38065,AT1G38131 |
| chr5:10179768 | ATGP6-LTR/Gypsy | AT5G28190,AT5G28210,AT5G28235,AT5G28237,AT5G28220 |
| chr3:12641372 | ATGP8-LTR/Gypsy | AT3G30840,AT3G30841,AT3G30845,AT3G30842 |
| chr3:14328918 | ATGP9B-LTR/Gypsy | AT3G42160,AT3G42170,AT3G42180 |
| chr1:18519237 | ATHAT1-DNA/HAT | AT1G49970,AT1G49975,AT1G49980,AT1G50000,AT1G50010,AT1G50020,AT1G49990,AT1G50030 |
| chr1:15983870 | ATHAT3-DNA/HAT | AT1G42525,AT1G42550,AT1G42560,AT1G42540 |
| chr3:11813775 | ATHAT7-DNA/HAT | AT3G30180 |
| chr1:15890614 | ATHATN1-DNA/HAT | AT1G41880,AT1G41920,AT1G42080,AT1G42190,AT1G42440,AT1G42470,AT1G42480,AT1G42430 |
| chr1:14243988 | ATHATN1-DNA/HAT | AT1G36940,AT1G36942,AT1G36950,AT1G36960,AT1G36970,AT1G36990,AT1G37000,AT1G37020,AT1G37113,AT1G37130,AT1G37140,AT1G37150,AT1G38065,AT1G38131,AT1G37010,AT1G36980 |
| chr1:15590950 | ATHILA7-LTR/Gypsy | AT1G41820,AT1G41830 |
| chr1:15923352 | ATHPOGO-DNA/Pogo | AT1G41830,AT1G41880,AT1G41920,AT1G42080,AT1G42190,AT1G42440,AT1G42470,AT1G42480,AT1G42430 |
| chr1:13717668 | ATHPOGO-DNA/Pogo | AT1G35820,AT1G35830,AT1G35850,AT1G35860,AT1G35880,AT1G35890,AT1G35910,AT1G36000,AT1G36005,AT1G36020,AT1G36030,AT1G36060,AT1G36078,AT1G36085,AT1G36095,AT1G36100,AT1G36150,AT1G36230,AT1G36272,AT1G36280,AT1G36310,AT1G36320,AT1G36325,AT1G36340,AT1G36380,AT1G36390,AT1G36580,AT1G36622,AT1G36623,AT1G36627,AT1G36640,AT1G36675,AT1G36730,AT1G36745,AT1G36756,AT1G36920,AT1G36922,AT1G36925,AT1G36940,AT1G36942,AT1G36950,AT1G36960,AT1G36970,AT1G36990,AT1G37000,AT1G37020,AT1G35895,AT1G36050,AT1G36070,AT1G36160,AT1G36180,AT1G36240,AT1G36370,AT1G37010,AT1G36510,AT1G36980 |
| chr3:3694956 | ATLINE1_4-LINE/L1 | AT3G11660,AT3G11670,AT3G11680,AT3G11690,AT3G11700,AT3G11720,AT3G11730,AT3G11710 |
| chr2:5835322 | ATREP12-RC/Helitron | AT2G13720,AT2G13760,AT2G13770,AT2G13790,AT2G13800,AT2G13810,AT2G13820,AT2G13840,AT2G13845,AT2G13895,AT2G13900,AT2G13905,AT2G13950,AT2G13960,AT2G13965,AT2G13980,AT2G13985,AT2G14000,AT2G14045,AT2G14050,AT2G14070,AT2G14080,AT2G14095,AT2G14100,AT2G14110,AT2G14120,AT2G14160,AT2G14210,AT2G14247,AT2G14255,AT2G14260,AT2G14265,AT2G14282,AT2G14285,AT2G14288,AT2G14289,AT2G14290,AT2G14365,AT2G14378,AT2G14060,AT2G14170,AT2G14270 |
| chr5:8441442 | ATREP18-DNA | AT5G24655,AT5G24660,AT5G24670 |

| | | |
|---|---|---|
| chr4:1542814 | SADHU-Unassigned | AT4G03440,AT4G03443,AT4G03450,AT4G03470,AT4G03480,AT4G03490,AT4G03495,AT4G03505,AT4G03510,AT4G03520,AT4G03540,AT4G03460,AT4G03500 |
| chr2:5166963 | VANDAL9-DNA/MuDR | AT2G12475,AT2G12480,AT2G12550,AT2G12646 |

Table S10. Summary of the overlapped candidate genes in the two stage GWAS analysis.

| Gene | Lead SNP (expression) | Lead SNP (copy number) |
|---|---|---|
| AT1G10870 | chr1:3627108:ATCOPIA23 | chr1:3636643:ATCOPIA53-LTR/Copia |
| AT1G10875 | chr1:3627108:ATCOPIA23 | chr1:3636643:ATCOPIA53-LTR/Copia |
| AT1G10880 | chr1:3627108:ATCOPIA23 | chr1:3636643:ATCOPIA53-LTR/Copia |
| AT1G10890 | chr1:3627108:ATCOPIA23 | chr1:3636643:ATCOPIA53-LTR/Copia |
| AT1G10900 | chr1:3627108:ATCOPIA23 | chr1:3636643:ATCOPIA53-LTR/Copia |
| AT1G10910 | chr1:3627108:ATCOPIA23 | chr1:3636643:ATCOPIA53-LTR/Copia |
| AT1G10920 | chr1:3627108:ATCOPIA23 | chr1:3636643:ATCOPIA53-LTR/Copia |
| AT1G10930 | chr1:3627108:ATCOPIA23 | chr1:3636643:ATCOPIA53-LTR/Copia |
| AT1G35140 | chr1:12817581:ATCOPIA36,chr1:12817581:ATCOPIA57 | chr1:13109197:ATCOPIA33-LTR/Copia |
| AT1G35150 | chr1:12817581:ATCOPIA36,chr1:12817581:ATCOPIA57 | chr1:13109197:ATCOPIA33-LTR/Copia |
| AT1G35160 | chr1:12817581:ATCOPIA36,chr1:12817581:ATCOPIA57 | chr1:13109197:ATCOPIA33-LTR/Copia |
| AT1G35170 | chr1:12817581:ATCOPIA36,chr1:12817581:ATCOPIA57 | chr1:13109197:ATCOPIA33-LTR/Copia |
| AT1G35180 | chr1:12817581:ATCOPIA36,chr1:12817581:ATCOPIA57 | chr1:13109197:ATCOPIA33-LTR/Copia |
| AT1G35181 | chr1:12817581:ATCOPIA36,chr1:12817581:ATCOPIA57 | chr1:13109197:ATCOPIA33-LTR/Copia |
| AT1G35183 | chr1:12817581:ATCOPIA57 | chr1:13109197:ATCOPIA33-LTR/Copia |
| AT1G35190 | chr1:12817581:ATCOPIA57 | chr1:13109197:ATCOPIA33-LTR/Copia |
| AT1G35210 | chr1:12817581:ATCOPIA57 | chr1:13109197:ATCOPIA33-LTR/Copia |
| AT1G35215 | chr1:12817581:ATCOPIA57 | chr1:13109197:ATCOPIA33-LTR/Copia |
| AT1G35220 | chr1:12817581:ATCOPIA57 | chr1:13109197:ATCOPIA33-LTR/Copia |
| AT1G35230 | chr1:12817581:ATCOPIA57 | chr1:13109197:ATCOPIA33-LTR/Copia |
| AT1G35240 | chr1:12817581:ATCOPIA57 | chr1:13109197:ATCOPIA33-LTR/Copia |
| AT1G35242 | chr1:12817581:ATCOPIA57 | chr1:13109197:ATCOPIA33-LTR/Copia |
| AT1G35250 | chr1:12817581:ATCOPIA57 | chr1:13109197:ATCOPIA33-LTR/Copia |
| AT1G35255 | chr1:12817581:ATCOPIA57 | chr1:13109197:ATCOPIA33-LTR/Copia |
| AT1G35260 | chr1:12817581:ATCOPIA57 | chr1:13109197:ATCOPIA33-LTR/Copia |
| AT1G35290 | chr1:12817581:ATCOPIA57 | chr1:13109197:ATCOPIA33-LTR/Copia |
| AT1G35310 | chr1:12817581:ATCOPIA57 | chr1:13109197:ATCOPIA33-LTR/Copia |
| AT1G35320 | chr1:12817581:ATCOPIA57 | chr1:13109197:ATCOPIA33-LTR/Copia |

| | | |
|---|---|---|
| AT1G35330 | chr1:12817581:ATCOPIA57 | chr1:13109197:ATCOPIA33-LTR/Copia |
| AT1G35620 | chr1:13255134:ENDOVIR1 | chr1:13109197:ATCOPIA33-LTR/Copia |
| AT1G35625 | chr1:13255134:ENDOVIR1 | chr1:13109197:ATCOPIA33-LTR/Copia |
| AT1G35630 | chr1:13255134:ENDOVIR1 | chr1:13109197:ATCOPIA33-LTR/Copia |
| AT1G35660 | chr1:13210172:ATCOPIA43,chr1:13210878:ATREP3,chr1:13255134:ENDOVIR1 | chr1:13109197:ATCOPIA33-LTR/Copia |
| AT1G35670 | chr1:13210172:ATCOPIA43,chr1:13210878:ATREP3,chr1:13255134:ENDOVIR1 | chr1:13109197:ATCOPIA33-LTR/Copia |
| AT1G35680 | chr1:13210172:ATCOPIA43,chr1:13210878:ATREP3,chr1:13255134:ENDOVIR1 | chr1:13109197:ATCOPIA33-LTR/Copia |
| AT1G35710 | chr1:13210172:ATCOPIA43,chr1:13210878:ATREP3,chr1:13255134:ENDOVIR1 | chr1:13109197:ATCOPIA33-LTR/Copia |
| AT1G35880 | chr1:13352257:HELITRON1 | chr1:13717668:ATHPOGO-DNA/Pogo |
| AT1G35890 | chr1:13352257:HELITRON1 | chr1:13717668:ATHPOGO-DNA/Pogo |
| AT1G35895 | chr1:13352257:HELITRON1 | chr1:13717668:ATHPOGO-DNA/Pogo |
| AT1G35910 | chr1:13352257:HELITRON1 | chr1:13717668:ATHPOGO-DNA/Pogo |
| AT1G36730 | chr1:13950766:BRODYAGA1A | chr1:13717668:ATHPOGO-DNA/Pogo |
| AT1G36745 | chr1:13950766:BRODYAGA1A | chr1:13717668:ATHPOGO-DNA/Pogo |
| AT1G36756 | chr1:13950766:BRODYAGA1A | chr1:13717668:ATHPOGO-DNA/Pogo |
| AT1G36920 | chr1:13950766:BRODYAGA1A | chr1:13717668:ATHPOGO-DNA/Pogo |
| AT1G36922 | chr1:13950766:BRODYAGA1A | chr1:13717668:ATHPOGO-DNA/Pogo |
| AT1G36925 | chr1:13950766:BRODYAGA1A | chr1:13717668:ATHPOGO-DNA/Pogo |
| AT1G36940 | chr1:13950766:BRODYAGA1A | chr1:14243988:ATHATN1-DNA/HAT,chr1:13717668:ATHPOGO-DNA/Pogo |
| AT1G36942 | chr1:13950766:BRODYAGA1A | chr1:14243988:ATHATN1-DNA/HAT,chr1:13717668:ATHPOGO-DNA/Pogo |
| AT1G36950 | chr1:13950766:BRODYAGA1A | chr1:14243988:ATHATN1-DNA/HAT,chr1:13717668:ATHPOGO-DNA/Pogo |
| AT1G36960 | chr1:13950766:BRODYAGA1A | chr1:14243988:ATHATN1-DNA/HAT,chr1:13717668:ATHPOGO-DNA/Pogo |
| AT1G36970 | chr1:13950766:BRODYAGA1A | chr1:14243988:ATHATN1-DNA/HAT,chr1:13717668:ATHPOGO-DNA/Pogo |
| AT1G36980 | chr1:13950766:BRODYAGA1A | chr1:14243988:ATHATN1-DNA/HAT,chr1:13717668:ATHPOGO-DNA/Pogo |
| AT1G36990 | chr1:13950766:BRODYAGA1A | chr1:14243988:ATHATN1-DNA/HAT,chr1:13717668:ATHPOGO-DNA/Pogo |

| | | |
|---|---|---|
| AT1G37000 | chr1:13950766:BRODYAGA1A | chr1:14243988:ATHATN1-DNA/HAT,chr1:13717668:ATHPOGO-DNA/Pogo |
| AT1G37010 | chr1:13950766:BRODYAGA1A | chr1:14243988:ATHATN1-DNA/HAT,chr1:13717668:ATHPOGO-DNA/Pogo |
| AT1G37020 | chr1:13950766:BRODYAGA1A | chr1:14243988:ATHATN1-DNA/HAT,chr1:13717668:ATHPOGO-DNA/Pogo |
| AT1G51200 | chr1:19484366:ATCOPIA59 | chr1:19003520:ATCOPIA68-LTR/Copia |
| AT1G51210 | chr1:19484366:ATCOPIA59 | chr1:19003520:ATCOPIA68-LTR/Copia |
| AT1G51220 | chr1:19484366:ATCOPIA59 | chr1:19003520:ATCOPIA68-LTR/Copia |
| AT1G51230 | chr1:19484366:ATCOPIA59 | chr1:19003520:ATCOPIA68-LTR/Copia |
| AT1G51240 | chr1:19484366:ATCOPIA59 | chr1:19003520:ATCOPIA68-LTR/Copia |
| AT1G51250 | chr1:19484366:ATCOPIA59 | chr1:19003520:ATCOPIA68-LTR/Copia |
| AT1G51260 | chr1:19484366:ATCOPIA59 | chr1:19003520:ATCOPIA68-LTR/Copia |
| AT1G51270 | chr1:19484366:ATCOPIA59 | chr1:19003520:ATCOPIA68-LTR/Copia |
| AT1G51290 | chr1:19484366:ATCOPIA59 | chr1:19003520:ATCOPIA68-LTR/Copia |
| AT1G51300 | chr1:19484366:ATCOPIA59 | chr1:19003520:ATCOPIA68-LTR/Copia |
| AT1G51310 | chr1:19484366:ATCOPIA59 | chr1:19003520:ATCOPIA68-LTR/Copia |
| AT2G10950 | chr2:4007630:ATCOPIA32B,chr2:4114169:ATCOPIA32 | chr2:4328548:ARNOLD4-DNA/MuDR |
| AT2G10955 | chr2:4007630:ATCOPIA32B,chr2:4114169:ATCOPIA32 | chr2:4328548:ARNOLD4-DNA/MuDR |
| AT2G10965 | chr2:4007630:ATCOPIA32B,chr2:4114169:ATCOPIA32 | chr2:4328548:ARNOLD4-DNA/MuDR |
| AT2G10970 | chr2:4007630:ATCOPIA32B,chr2:4114169:ATCOPIA32 | chr2:4328548:ARNOLD4-DNA/MuDR |
| AT2G10975 | chr2:4007630:ATCOPIA32B,chr2:4114169:ATCOPIA32 | chr2:4328548:ARNOLD4-DNA/MuDR,chr2:4348901:ATENSPM1A-DNA/En-Spm |
| AT2G11000 | chr2:4007630:ATCOPIA32B,chr2:4114169:ATCOPIA32 | chr2:4328548:ARNOLD4-DNA/MuDR,chr2:4348901:ATENSPM1A-DNA/En-Spm |
| AT2G11005 | chr2:4007630:ATCOPIA32B,chr2:4114169:ATCOPIA32 | chr2:4328548:ARNOLD4-DNA/MuDR,chr2:4348901:ATENSPM1A-DNA/En-Spm |
| AT2G11010 | chr2:4007630:ATCOPIA32B,chr2:4114169:ATCOPIA32 | chr2:4348901:ATENSPM1A-DNA/En-Spm |
| AT2G11015 | chr2:4007630:ATCOPIA32B,chr2:4114169:ATCOPIA32 | chr2:4348901:ATENSPM1A-DNA/En-Spm |
| AT2G11025 | chr2:4114169:ATCOPIA32 | chr2:4348901:ATENSPM1A-DNA/En-Spm |
| AT2G13840 | chr2:5800623:ATHAT10 | chr2:5835322:ATREP12-RC/Helitron |
| AT2G13845 | chr2:5800623:ATHAT10 | chr2:5835322:ATREP12-RC/Helitron |
| AT2G13895 | chr2:5800623:ATHAT10 | chr2:5835322:ATREP12-RC/Helitron |

| | | |
|---|---|---|
| AT2G13900 | chr2:5800623:ATHAT10 | chr2:5835322:ATREP12-RC/Helitron |
| AT2G13905 | chr2:5800623:ATHAT10 | chr2:5835322:ATREP12-RC/Helitron |
| AT2G14210 | chr2:6023757:ATCOPIA77 | chr2:5835322:ATREP12-RC/Helitron |
| AT2G14247 | chr2:6023757:ATCOPIA77 | chr2:5835322:ATREP12-RC/Helitron |
| AT2G14255 | chr2:6023757:ATCOPIA77 | chr2:5835322:ATREP12-RC/Helitron |
| AT2G14260 | chr2:6023757:ATCOPIA77 | chr2:5835322:ATREP12-RC/Helitron |
| AT2G14265 | chr2:6023757:ATCOPIA77 | chr2:5835322:ATREP12-RC/Helitron |
| AT2G14270 | chr2:6023757:ATCOPIA77 | chr2:5835322:ATREP12-RC/Helitron |
| AT2G14282 | chr2:6023757:ATCOPIA77 | chr2:5835322:ATREP12-RC/Helitron |
| AT2G16640 | chr2:7230176:ATCOPIA70 | chr2:6906197:ATCOPIA80-LTR/Copia |
| AT2G16650 | chr2:7230176:ATCOPIA70 | chr2:6906197:ATCOPIA80-LTR/Copia |
| AT2G16660 | chr2:7230176:ATCOPIA70 | chr2:6906197:ATCOPIA80-LTR/Copia |
| AT2G16676 | chr2:7230176:ATCOPIA70 | chr2:6906197:ATCOPIA80-LTR/Copia |
| AT2G16700 | chr2:7230176:ATCOPIA70 | chr2:6906197:ATCOPIA80-LTR/Copia |
| AT2G16710 | chr2:7230176:ATCOPIA70 | chr2:6906197:ATCOPIA80-LTR/Copia |
| AT2G16750 | chr2:7298605:VANDAL14 | chr2:6906197:ATCOPIA80-LTR/Copia |
| AT2G16760 | chr2:7298605:VANDAL14 | chr2:6906197:ATCOPIA80-LTR/Copia |
| AT2G16770 | chr2:7298605:VANDAL14 | chr2:6906197:ATCOPIA80-LTR/Copia |
| AT3G32960 | chr3:13493764:ARNOLD3 | chr3:13577002:ATDNA1T9A-DNA/MuDR |
| AT3G32980 | chr3:13493764:ARNOLD3 | chr3:13577002:ATDNA1T9A-DNA/MuDR |
| AT3G42160 | chr3:14167801:ATLANTYS1 | chr3:14328918:ATGP9B-LTR/Gypsy |
| AT3G45000 | chr3:16601301:TNAT2A | chr3:16461148:ATCOPIA59-LTR/Copia |
| AT3G45010 | chr3:16601301:TNAT2A | chr3:16461148:ATCOPIA59-LTR/Copia |
| AT3G45020 | chr3:16601301:TNAT2A | chr3:16461148:ATCOPIA59-LTR/Copia |
| AT3G45030 | chr3:16601301:TNAT2A | chr3:16461148:ATCOPIA59-LTR/Copia |
| AT3G45040 | chr3:16601301:TNAT2A | chr3:16461148:ATCOPIA59-LTR/Copia |
| AT3G45050 | chr3:16601301:TNAT2A | chr3:16461148:ATCOPIA59-LTR/Copia |
| AT3G45060 | chr3:16601301:TNAT2A | chr3:16461148:ATCOPIA59-LTR/Copia |
| AT3G45070 | chr3:16601301:TNAT2A | chr3:16461148:ATCOPIA59-LTR/Copia |
| AT4G03460 | chr4:1559762:VANDAL5 | chr4:1542814:SADHU-Unassigned |

| | | |
|---|---|---|
| AT4G03470 | chr4:1559762:VANDAL5 | chr4:1542814:SADHU-Unassigned |
| AT4G03480 | chr4:1559762:VANDAL5 | chr4:1542814:SADHU-Unassigned |
| AT4G03490 | chr4:1559762:VANDAL5 | chr4:1542814:SADHU-Unassigned |
| AT4G03495 | chr4:1559762:VANDAL5 | chr4:1542814:SADHU-Unassigned |
| AT4G03500 | chr4:1559762:VANDAL5 | chr4:1542814:SADHU-Unassigned |
| AT4G03505 | chr4:1559762:VANDAL5 | chr4:1542814:SADHU-Unassigned |
| AT4G03510 | chr4:1559762:VANDAL5 | chr4:1542814:SADHU-Unassigned |
| AT4G03520 | chr4:1559762:VANDAL5 | chr4:1542814:SADHU-Unassigned |
| AT4G03540 | chr4:1559762:VANDAL5 | chr4:1542814:SADHU-Unassigned |
| AT4G13620 | chr4:7956078:ATLINE1_5 | chr4:7947027:ATCOPIA79-LTR/Copia |
| AT4G13630 | chr4:7956078:ATLINE1_5 | chr4:7947027:ATCOPIA79-LTR/Copia |
| AT4G13640 | chr4:7956078:ATLINE1_5 | chr4:7947027:ATCOPIA79-LTR/Copia |
| AT4G13650 | chr4:7956078:ATLINE1_5 | chr4:7947027:ATCOPIA79-LTR/Copia |
| AT4G13660 | chr4:7956078:ATLINE1_5 | chr4:7947027:ATCOPIA79-LTR/Copia |
| AT4G13670 | chr4:7956078:ATLINE1_5 | chr4:7947027:ATCOPIA79-LTR/Copia |
| AT4G13680 | chr4:7956078:ATLINE1_5 | chr4:7947027:ATCOPIA79-LTR/Copia |
| AT4G13690 | chr4:7956078:ATLINE1_5 | chr4:7947027:ATCOPIA79-LTR/Copia |
| AT4G13700 | chr4:7956078:ATLINE1_5 | chr4:7947027:ATCOPIA79-LTR/Copia |
| AT5G35050 | chr5:13350109:ATCOPIA15 | chr5:12904117:ATCOPIA43-LTR/Copia |
| AT5G35067 | chr5:13350109:ATCOPIA15 | chr5:12904117:ATCOPIA43-LTR/Copia |
| AT5G35069 | chr5:13350109:ATCOPIA15 | chr5:12904117:ATCOPIA43-LTR/Copia |
| AT5G40030 | chr5:16048359:ATCOPIA18A | chr5:16030044:ATDNAI26T9-DNA/MuDR |
| AT5G40040 | chr5:16048359:ATCOPIA18A | chr5:16030044:ATDNAI26T9-DNA/MuDR |
| AT5G40050 | chr5:16048359:ATCOPIA18A | chr5:16030044:ATDNAI26T9-DNA/MuDR |
| AT5G40060 | chr5:16048359:ATCOPIA18A | chr5:16030044:ATDNAI26T9-DNA/MuDR |
| AT5G40070 | chr5:16048359:ATCOPIA18A | chr5:16030044:ATDNAI26T9-DNA/MuDR |
| AT5G40080 | chr5:16048359:ATCOPIA18A | chr5:16030044:ATDNAI26T9-DNA/MuDR |
| AT5G40090 | chr5:16048359:ATCOPIA18A | chr5:16030044:ATDNAI26T9-DNA/MuDR |
| AT5G40100 | chr5:16048359:ATCOPIA18A | chr5:16030044:ATDNAI26T9-DNA/MuDR |

Table S11. Summary of the candidate genes whose LoF mutation is associated with TE expression level or copy number variation.

| Gene | GWAS type | TE family | LoF | Non-LoF | FDR | Freq |
|---|---|---|---|---|---|---|
| AT3G30730 | copy number | ATCOPIA63-LTR/Copia | 3.58 | 3.88 | 8.80E-04 | 0.00 |
| AT3G43950 | expression | ATCOPIA62-LTR/Copia | 250.20 | 443.58 | 2.10E-04 | 1.00 |
| AT2G10556 | expression | ATCOPIA32B-LTR/Copia | 5.63 | 29.32 | 1.96E-03 | 0.00 |
| AT1G35310 | copy number | ATCOPIA33-LTR/Copia | 1.82 | 1.95 | 9.81E-03 | 0.00 |
| AT4G11170 | expression | ATCOPIA17-LTR/Copia | 29.32 | 93.20 | 9.35E-04 | 0.52 |
| AT4G11170 | expression | ATHILA4D_LTR-LTR/Gypsy | 10.56 | 18.33 | 6.87E-07 | 0.52 |
| AT1G35710 | expression | ATCOPIA43-LTR/Copia | 91.86 | 20.17 | 4.25E-23 | 0.00 |
| AT1G35710 | expression | ATREP3-RC/Helitron | 284.84 | 204.81 | 4.17E-08 | 0.00 |
| AT2G09970 | expression | ATCOPIA32-LTR/Copia | 164.03 | 99.77 | 1.23E-05 | 0.00 |
| AT2G09388 | expression | ATCOPIA32-LTR/Copia | 137.37 | 106.46 | 5.21E-03 | 0.37 |
| AT2G11015 | expression | ATCOPIA32B-LTR/Copia | 28.81 | 15.28 | 2.83E-03 | 0.00 |
| AT5G35510 | expression | ATCOPIA24-LTR/Copia | 91.84 | 25.38 | 9.35E-04 | 1.00 |
| AT2G10545 | expression | ATCOPIA32-LTR/Copia | 72.91 | 152.45 | 8.02E-03 | 0.82 |
| AT2G25590 | expression | ATCOPIA65-LTR/Copia | 6.05 | 9.09 | 8.89E-03 | 0.00 |
| AT4G05210 | expression | VANDAL5-DNA/MuDR | 107.01 | 26.78 | 9.81E-03 | 0.00 |
| AT1G52560 | expression | ATCOPIA81-LTR/Copia | 36.90 | 32.70 | 5.73E-04 | 0.00 |
| AT3G32920 | expression | ATLANTYS1-LTR/Gypsy | 296.19 | 580.07 | 8.32E-13 | 0.12 |
| AT2G40850 | expression | ATDNAI27T9C-DNA/MuDR | 13.21 | 5.23 | 6.26E-03 | 0.08 |
| AT3G30770 | copy number | ATCOPIA63-LTR/Copia | 3.63 | 3.94 | 2.83E-03 | 0.12 |
| AT4G08267 | copy number | ATCOPIA44-LTR/Copia | 4.03 | 3.60 | 2.83E-03 | 0.00 |
| AT1G12190 | expression | ATGP3-LTR/Gypsy | 91.67 | 227.48 | 6.20E-03 | 0.33 |
| AT1G61280 | expression | ATHILA4C-LTR/Gypsy | 449.14 | 200.66 | 8.27E-04 | 0.00 |
| AT3G28170 | expression | ATCOPIA50-LTR/Copia | 19.17 | 97.08 | 1.73E-04 | 0.84 |
| AT3G28170 | expression | ATCOPIA74-LTR/Copia | 14.34 | 170.45 | 5.52E-09 | 0.84 |
| AT2G41590 | expression | ATDNAI27T9C-DNA/MuDR | 18.66 | 6.61 | 2.97E-03 | 0.77 |
| AT2G17060 | expression | ATCOPIA71-LTR/Copia | 206.94 | 11.73 | 4.43E-04 | 0.06 |
| AT3G43970 | expression | ATCOPIA62-LTR/Copia | 280.50 | 481.30 | 1.24E-05 | 0.86 |
| AT1G36627 | copy number | ATHPOGO-DNA/Pogo | 2.77 | 2.92 | 2.12E-03 | 0.00 |
| AT5G44830 | expression | ATDNA2T9B-DNA/MuDR | 11.86 | 3.70 | 1.18E-04 | 0.00 |
| AT5G54745 | expression | HELITRONY1E-RC/Helitron | 119.47 | 48.41 | 2.75E-08 | 0.00 |
| AT2G10955 | expression | ATCOPIA32B-LTR/Copia | 44.60 | 19.41 | 3.93E-04 | 0.18 |
| AT4G03580 | expression | VANDAL5-DNA/MuDR | 19.97 | 36.11 | 4.83E-03 | 0.63 |
| AT1G54030 | expression | ATHPOGON3-DNA/Pogo | 23.40 | 14.65 | 1.70E-05 | 0.92 |
| AT5G47400 | expression | ATCOPIA61-LTR/Copia | 91.90 | 158.51 | 6.96E-11 | 0.05 |
| AT5G36080 | expression | ATCOPIA91-LTR/Copia | 54.56 | 126.11 | 7.55E-03 | 0.00 |
| AT3G44006 | expression | ATCOPIA62-LTR/Copia | 267.82 | 446.06 | 8.39E-03 | 0.88 |
| AT3G42160 | expression | ARNOLD3-DNA/MuDR | 21.80 | 7.66 | 3.87E-03 | 0.00 |
| AT3G42160 | expression | ATLANTYS1-LTR/Gypsy | 744.14 | 351.23 | 1.70E-05 | 0.00 |
| AT4G10890 | expression | ATCOPIA17-LTR/Copia | 30.00 | 173.57 | 5.21E-03 | 1.00 |
| AT3G28020 | expression | ATCOPIA74-LTR/Copia | 18.68 | 62.46 | 8.27E-04 | 0.16 |

| | | | | | | |
|---|---|---|---|---|---|---|
| AT3G30739 | copy number | ATCOPIA63-LTR/Copia | 3.57 | 3.95 | 6.61E-05 | 0.12 |
| AT4G28920 | expression | ATCOPIA46-LTR/Copia | 47.37 | 154.63 | 1.26E-03 | 0.96 |
| AT5G38150 | expression | RP1_AT-DNA | 140.91 | 23.51 | 5.57E-04 | 0.01 |
| AT5G38080 | expression | ATDNA12T3_2-DNA | 278.89 | 34.63 | 4.35E-03 | 1.00 |
| AT5G35930 | expression | ATCOPIA91-LTR/Copia | 52.31 | 136.99 | 5.72E-08 | 0.00 |
| AT4G10360 | expression | TAG1-DNA/HAT | 2.34 | 0.90 | 4.10E-06 | 0.00 |
| AT1G42440 | copy number | ATHATN1-DNA/HAT | 2.22 | 1.92 | 1.20E-03 | 1.00 |
| AT3G50480 | expression | ATSINE4-SINE | 68.94 | 24.86 | 2.83E-03 | 0.00 |
| AT5G35926 | expression | ATCOPIA91-LTR/Copia | 80.52 | 200.36 | 6.01E-04 | 1.00 |
| AT2G25480 | copy number | ATCOPIA39-LTR/Copia | 0.42 | 0.19 | 4.64E-04 | 0.00 |
| AT1G35140 | expression | ATCOPIA36-LTR/Copia | 236.72 | 1177.03 | 1.57E-22 | 0.00 |
| AT1G35140 | expression | ATCOPIA57-LTR/Copia | 15.78 | 63.53 | 1.09E-19 | 0.00 |
| AT5G36190 | expression | ATCOPIA91-LTR/Copia | 214.83 | 86.45 | 7.47E-06 | 0.00 |
| AT5G35950 | expression | ATCOPIA91-LTR/Copia | 56.44 | 112.59 | 2.24E-03 | 0.49 |
| AT2G41470 | expression | ATDNAI27T9C-DNA/MuDR | 16.21 | 4.62 | 1.27E-05 | 0.00 |
| AT3G44080 | expression | ATCOPIA62-LTR/Copia | 290.90 | 614.89 | 2.23E-06 | 1.00 |
| AT3G44090 | expression | ARNOLDY1-DNA/MuDR | 30.45 | 23.47 | 4.87E-03 | 0.01 |
| AT3G44090 | expression | ATCOPIA62-LTR/Copia | 221.53 | 393.20 | 1.27E-05 | 0.01 |
| AT1G35030 | expression | ATCOPIA36-LTR/Copia | 601.04 | 879.49 | 4.02E-04 | 0.82 |
| AT1G35030 | expression | ATCOPIA57-LTR/Copia | 34.16 | 50.94 | 8.27E-04 | 0.82 |
| AT5G35375 | expression | ATCOPIA24-LTR/Copia | 61.13 | 29.04 | 9.35E-04 | 0.99 |
| AT2G14265 | expression | ATCOPIA77-LTR/Copia | 159.90 | 63.52 | 2.06E-04 | 0.98 |
| AT1G34760 | expression | ATCOPIA57-LTR/Copia | 15.33 | 53.12 | 4.43E-04 | 0.00 |
| AT1G35040 | expression | ATCOPIA36-LTR/Copia | 104.97 | 1087.78 | 5.36E-16 | 0.00 |
| AT1G35040 | expression | ATCOPIA57-LTR/Copia | 9.84 | 59.24 | 1.41E-14 | 0.00 |
| AT5G36160 | expression | ATCOPIA91-LTR/Copia | 37.86 | 123.33 | 8.09E-04 | 0.93 |
| AT1G35240 | expression | ATCOPIA57-LTR/Copia | 25.43 | 60.56 | 1.14E-07 | 0.01 |
| AT5G54820 | expression | HELITRONY1E-RC/Helitron | 178.52 | 54.73 | 1.72E-11 | 0.00 |
| AT1G37113 | copy number | ATHATN1-DNA/HAT | 2.18 | 1.90 | 6.01E-04 | 0.99 |

Note: LoF (non-LoF), TE family expression level or TE copy number or total polymorphic TE number of accessions with (without) LoF mutations. Freq, the allele frequency of LoF in Yangtze River basin population.
